# Engineered ACE2 receptor traps potently neutralize SARS-CoV-2

**DOI:** 10.1101/2020.07.31.231746

**Authors:** Anum Glasgow, Jeff Glasgow, Daniel Limonta, Paige Solomon, Irene Lui, Yang Zhang, Matthew A. Nix, Nicholas J. Rettko, Shion A. Lim, Shoshana Zha, Rachel Yamin, Kevin Kao, Oren S. Rosenberg, Jeffrey V. Ravetch, Arun P. Wiita, Kevin K. Leung, Xin X. Zhou, Tom C. Hobman, Tanja Kortemme, James A. Wells

## Abstract

An essential mechanism for SARS-CoV-1 and -2 infection begins with the viral spike protein binding to the human receptor protein angiotensin-converting enzyme II (ACE2). Here we describe a stepwise engineering approach to generate a set of affinity optimized, enzymatically inactivated ACE2 variants that potently block SARS-CoV-2 infection of cells. These optimized receptor traps tightly bind the receptor binding domain (RBD) of the viral spike protein and prevent entry into host cells. We first computationally designed the ACE2-RBD interface using a two-stage flexible protein backbone design process that improved affinity for the RBD by up to 12-fold. These designed receptor variants were affinity matured an additional 14-fold by random mutagenesis and selection using yeast surface display. The highest affinity variant contained seven amino acid changes and bound to the RBD 170-fold more tightly than wild-type ACE2. With the addition of the natural ACE2 collectrin domain and fusion to a human Fc domain for increased stabilization and avidity, the most optimal ACE2 receptor traps neutralized SARS-CoV-2 pseudotyped lentivirus and authentic SARS-CoV-2 virus with half-maximal inhibitory concentrations (IC50) in the 10-100 ng/ml range. Engineered ACE2 receptor traps offer a promising route to fighting infections by SARS-CoV-2 and other ACE2-utilizing coronaviruses, with the key advantage that viral resistance would also likely impair viral entry. Moreover, such traps can be pre-designed for viruses with known entry receptors for faster therapeutic response without the need for neutralizing antibodies isolated or generated from convalescent patients.

---

There is an urgent need for broadly effective therapeutics to treat SARS-CoV-2 infections during the ongoing COVID-19 pandemic (1, 2). Antibodies isolated from convalescent patient sera and recombinant antibodies cloned from the B-cells of recovered patients have been effective in past and recent pandemics, and much of the ongoing drug development effort is based on these approaches (3–8). However, strategies for antibody development necessarily follow widespread viral spread and infection, which costs precious time in a rapidly developing pandemic.

Protein engineering approaches to identify binders to viral entry proteins offer a rapid alternative, without the prerequisite for an infected population. In the first step of a SARS-CoV-1 or CoV-2 infection, the receptor binding domain (RBD) of the trimeric spike protein on the surface of the virus binds to the membrane-bound receptor angiotensin-converting enzyme II (ACE2) to enter human cells (3, 4, 8). Most neutralizing antibodies to SARS-CoV-1 and CoV-2 block viral entry by binding to the ACE2 binding site on the RBD. Ongoing efforts by our lab and others use *in vitro* methods, such as phage display or yeast display, from naïve libraries to generate recombinant antibodies or other formatted domains to block viral entry (9, 10).

As an alternate strategy, we pursued development of ACE2 “receptor traps”: affinity-optimized soluble variants of the ACE2 extracellular domain that block the viral spike protein from binding cellular ACE2 and facilitating entry (11). This approach has the potential advantage that viral resistance to an ACE2 receptor trap would also inhibit the ability of the virus to enter via binding to the ACE2 entry receptor. Receptor traps would also be useful for both pandemic SARS-CoV-1 and CoV-2 as well as other emerging variant strains that use ACE2 as a common entry port. Furthermore, the soluble extracellular domain of wild-type (WT) human recombinant ACE2 (APN01) was found to be safe in healthy volunteers (12) and in a small cohort of patients with acute respiratory distress syndrome (13) by virtue of ACE2’s intrinsic angiotensin converting activity, which is not required for viral entry. APN01 is currently in phase II clinical trials in Europe for treatment of SARS-CoV-2 (14) (NCT04335136). However, we and others have shown that WT ACE2 binds the SARS-CoV-2 spike RBD with only modest affinity (K_D_ ∼15 nM) (15–17). ACE2 is therefore a good candidate for affinity optimization, especially because potent blocking antibodies to the spike protein can be isolated with binding affinity (K_D_) values in the mid- to low-pM range (3, 4, 6, 7, 9, 18–20).

Here we improve the binding affinity of ACE2 for the monomeric spike RBD by 170-fold using a hybrid computational and experimental protein engineering approach. We demonstrate that after fusion to a human IgG Fc domain and the natural collectrin domain of ACE2, our most effective ACE2-Fc variant has a half-maximal inhibitory concentration (IC50) of 28 ng/ml in pseudotyped SARS-CoV-2 neutralization assays and comparable neutralization in authentic SARS-CoV-2 infection assays, reducing viral replication to almost undetectable levels. ACE2 receptor traps are promising therapeutic candidates, especially given the potential for viral escape mutations to impact antibody efficacy (5, 21) and low neutralizing antibody levels in a subset of recovered patients (6).

## Results

We re-engineered the soluble extracellular domain of ACE2 (residues 18-614, ACE2(614)) to bind the RBD of the SARS-CoV-2 spike protein using a combined computational/experimental protein engineering strategy (Figure 1). First, we computationally redesigned ACE2(614) using the Rosetta macromolecular modeling suite, introducing sets of mutations that improved the K_D_ of an ACE2(614)-Fc fusion protein for the SARS-CoV-2 spike RBD from 3- to 11-fold over the WT ACE2(614)-Fc protein in bio-layer interferometry (BLI) binding assays. Then, we affinity-matured the improved ACE2(614) designs in a pooled yeast display format. Additional mutations discovered through yeast display conferred a further 14-fold improvement in the apparent binding affinity (K_D,app_) for the RBD over the computationally designed parent ACE2(614), as measured on the surface of yeast. The final ACE2 variants have K_D,app_ close to 100 pM for the monomeric spike RBD.

**Figure 1.**
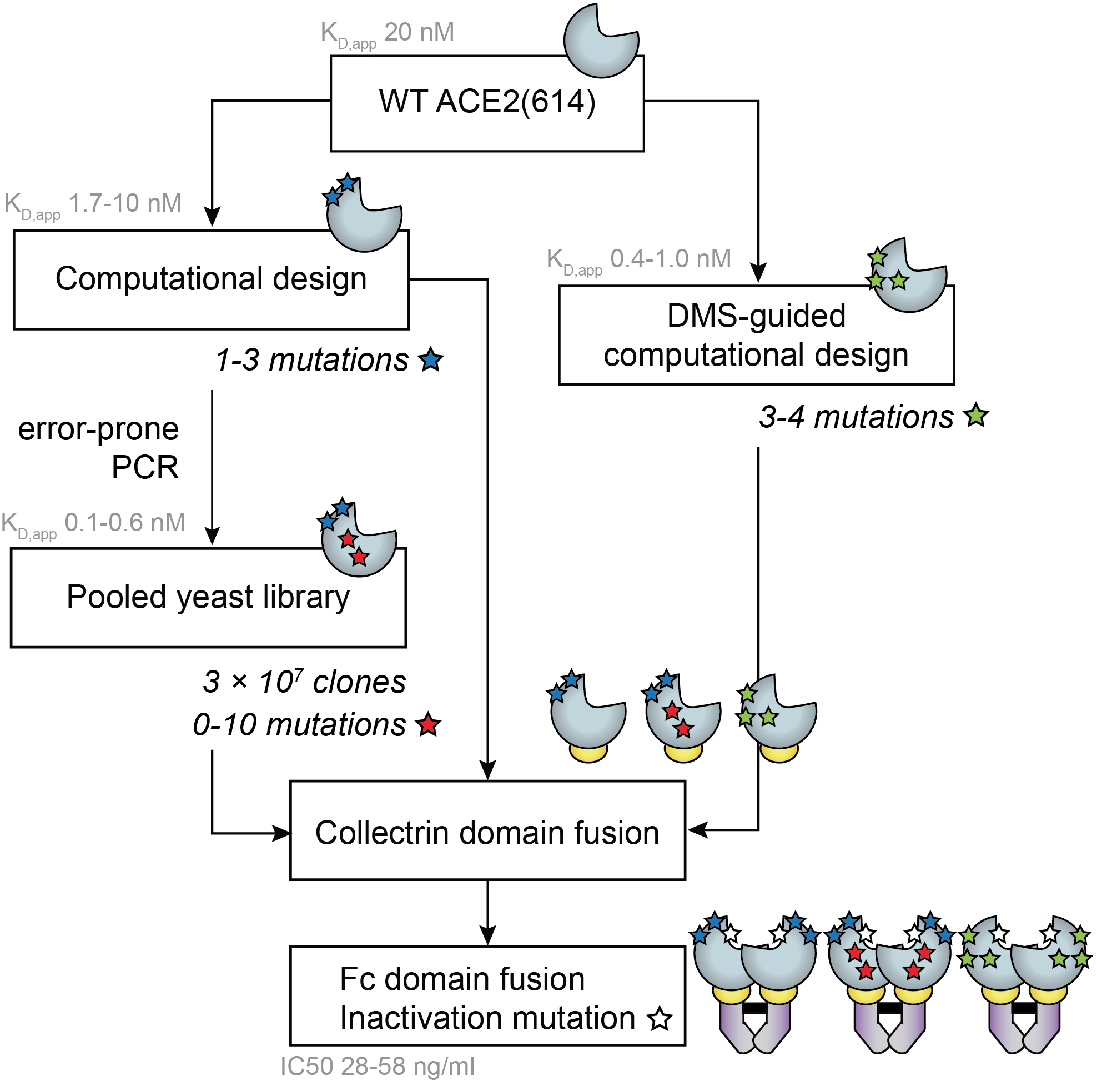
Protein engineering strategy to generate ACE2 receptor traps. Two independent computational design strategies were used to predict mutations to ACE2(614) (light blue shape) that enhance its affinity for the SARS-CoV-2 spike RBD: (1) saturation mutagenesis at hotspot positions determined by computational alanine scanning, followed by local ACE2 redesign (blue star mutations), and (2) combining individual mutations from computational saturation mutagenesis and experimental deep mutational scanning data (green star mutations). Four ACE2 variants determined from design strategy (1) were mutagenized and screened for binding to the RBD. ACE2 variants from the yeast library with additional mutations that improved the binding affinity for the RBD were isolated (red star mutations). Engineered ACE2(614) variants were fused to the ACE2 collectrin domain (residues 615-740, yellow ovals). ACE2(740) variants were expressed as Fc-fusions (purple shapes) with an additional mutation to inactivate ACE2 peptidase activity (white star mutation). K_D,app_ values represent apparent binding affinities measured between yeast surface-displayed ACE2(614) variants and the monomeric RBD. IC50 values represent half-maximal inhibitory concentrations measured for the four most potent neutralizing variants in SARS-CoV-2 pseudotyped virus assays.

High-resolution ACE2-RBD structures (22, 23) show a large, flat binding interface primarily comprising the N-terminal helices of ACE2 (residues 18-90), with secondary interaction sites spanning residues 324-361 (Figure 2A). To computationally redesign ACE2(614) for increased binding affinity with the RBD, we first determined which amino acid sidechains are most crucial to the ACE2-RBD interaction (“hotspots”) by performing a computational alanine scan on the binding interface using an established method in Rosetta (24, 25). Then, we systematically redesigned a subset of hotspot residues and their local environment to generate models for new interfaces, and selected the lowest (best) scoring ACE2 designs for testing (Figure 2A-C).

**Figure 2.**
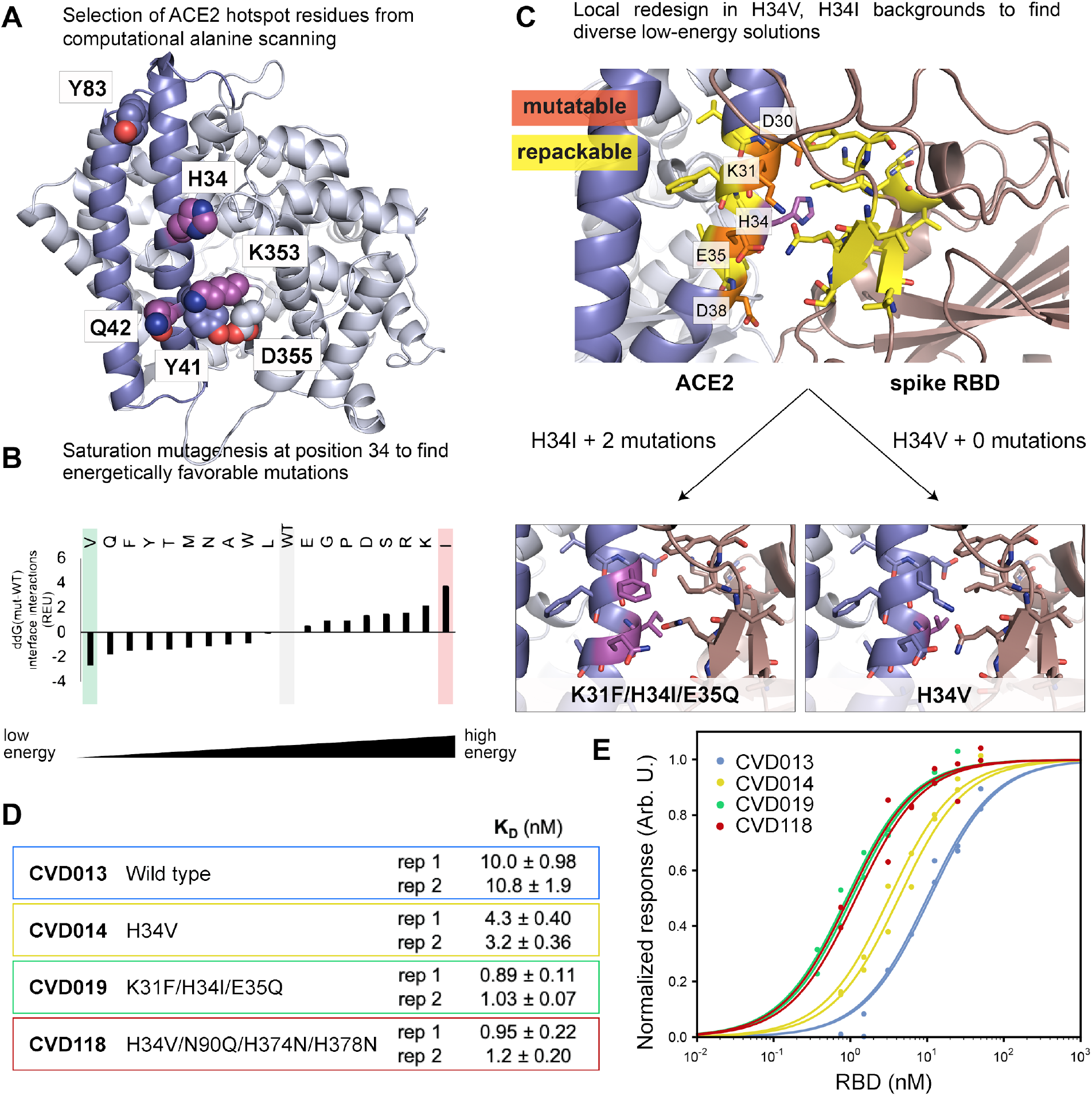
Computational design of ACE2 for improved binding affinity to the spike RBD. **(A)** Computational alanine scanning identified ACE2 hotspot residues (shown as spheres) that contribute strongly to binding the spike RBD. Residues 18-90 are shown in blue and residues 91-614 are shown in light blue. H34, Q42 and K353, shown as magenta spheres, were selected for computational saturation mutagenesis. **(B)** Computational saturation mutagenesis predicted several stabilizing mutations to H34. H34V and H34I were selected for further design to generate diverse ACE2 receptor trap models. **(C)** Flexible backbone design was performed around V34 ACE2 and I34 ACE2 (WT residue H34 shown in magenta). ACE2 residues that were permitted to change amino acid identity (“mutable” residues) are labeled and shown in orange; ACE2 and RBD residues that were allowed to change rotameric conformations and/or backbone atom positions (“repackable” residues) are shown in yellow. Design around H34I resulted in two additional mutations, shown in magenta. The N90Q glycan knockout mutation from DMS (28) was added to V34 ACE2. **(D-E)** *In vitro* BLI measurements show that designed ACE2(614)-Fc binding affinities to the RBD are improved 2- to 11-fold over WT ACE2(614)-Fc. To generate CVD118, the N90Q glycan knockout mutation from DMS (28) was added to V34 ACE2, as well as two histidine mutations to inactivate ACE2 peptidase activity (41). Each “rep” is a separate biological replicate. The table in **(D)** lists the K_D_ values for the designed ACE2 variants with errors of the fits for titration curves shown in **(E)**.

Computational alanine scanning suggested that the binding affinity of the ACE2-RBD interaction depends most crucially on 6 amino acid sidechains (H34, Y41, Q42, Y83, K353, and D355) on ACE2 as assessed by values of the predicted change in binding energy upon mutation to alanine (DDG(complex)) greater than 1 Rosetta Energy Unit (REU) (Table S1, Figure 2A). To determine which of these “hotspots” to target for computational design, we evaluated each hotspot residue by two metrics: the per-residue energy and the contribution of the residue to the interface energy (see Methods). Higher energies indicate lower stability. Hotspot residues H34, Q42, and K353 were targeted for further design.

To determine whether point mutations at these positions could improve the ACE2-RBD binding affinity, we performed computational saturation mutagenesis at these positions (excluding mutations to cysteine to avoid potential disulfide bond formation) and recalculated the interface energy for each model (Table S2). While we found no amino acid substitutions at positions 42 and 353 that were predicted to be stabilizing, several substitutions at position 34 were predicted to improve the interaction energy between ACE2 and the RBD (Figure 2B). Histidine 34 was mutated to a valine in the lowest-energy model because we anticipated favorable hydrophobic interactions with leucine 455 in the RBD. In the highest-energy model, histidine 34 was mutated to an overly bulky isoleucine.

We reasoned that both H34V and H34I, as the “best” and “worst” point mutants, were predicted to substantially affect the interface energy in the context of their chemical environment, and that additional local mutations might improve binding affinity in both models to yield different viable solutions. We applied the Rosetta “Coupled Moves” flexible backbone design protocol (26) to redesign the local environment of V34 and I34 in each model (Figure 2C). ACE2 sidechains within 4 Å of residue 34 were allowed to mutate, while other ACE2 and RBD sidechains within 8 Å of ACE2 residue 34 could change rotamer and/or backbone conformations (“repacking”, see Methods). This approach did not identify additional favorable mutations to H34V ACE2 in any of the models, but there were one to four additional mutations in the H34I ACE2 models. The lowest-energy designed protein based on H34I ACE2 had two additional mutations: K31F and E35Q (Figures 2C, S1). In this solution, ACE2 Q35 made a hydrogen bond with a repositioned RBD Q493, and ACE2 I34 packed against a repositioned RBD L455. On the ACE2 side of the interface, F31 made a favorable hydrophobic interaction with the methylene in Q35. For both lowest-energy redesigned interfaces, the root-mean-square deviations (RMSD) of the mutable and repackable positions (atoms corresponding to ACE2 residues 29-39 and RBD residues 416-418, 452-456, and 492-494) in the model vs. the WT structure were less than 1 Å, and the total summed predicted pairwise interface energies for both design solutions were comparable (Figure S1).

Next, we characterized the binding affinities of the computationally designed ACE2 variants as Fc fusions for spike RBD in BLI assays using purified proteins. We transiently expressed ACE2(614) with a C-terminal human IgG Fc domain fusion for improved affinity to spike, as shown in previous studies (11, 27), in Expi293 cells (see Methods). The BLI-measured K_D_ values of computationally-designed H34V ACE2(614)-Fc and K31F/H34I/E35Q ACE2(614)-Fc for the RBD were measured to be 3 and 11 times lower than the WT ACE2(614)-Fc, respectively (Figure 2D, E). We also tested binding of the RBD to H34V/N90Q/H374N/H378N ACE2(614)-Fc to determine the impact of removing a glycan that is adjacent to the interface at N90, as well as the protein’s native peptidase activity. A recent deep mutational scanning (DMS) study reported enrichment for ACE2 variants with mutations at the N90 glycosylation site (28); and histidines in position 374 and 378 together coordinate a Zn^2+^ ion necessary for enzymatic activity (Figure S2) (11). In the BLI assay, H34V/N90Q/H374N/H378N ACE2(614)-Fc showed 10-fold improved affinity over WT ACE2(614)-Fc (Figure 2D, E).

To further improve the binding affinity of the designed proteins for the spike RBD, we expressed a mutagenized library of ACE2(614) variants as Aga2p fusions for surface display in yeast cells, without the Fc domain to avoid avidity effects that might dominate affinity maturation. We chose four ACE2(614) variants as starting templates for a randomized yeast-displayed library, with the following mutations: H34V, N90Q, H34V/N90Q, and K31F/H34I/E35Q. Each of these variants, in addition to WT ACE2(614), were cloned as fusions to a Myc tag, Aga2p, and C-terminal enhanced GFP for a simple readout of induction and surface display level (Figure 3A) (29). The expression of WT ACE2(614) was first induced on EBY100 cells in SGCAA media at 20 °C and 30 °C, and we confirmed binding using biotinylated spike RBD-Fc and streptavidin Alexa Fluor 647 (Figure S3A, B). Binding of the RBD was well-correlated with GFP expression, precluding the need for myc-tag (expression) staining (Figure S3A, B).

**Figure 3.**
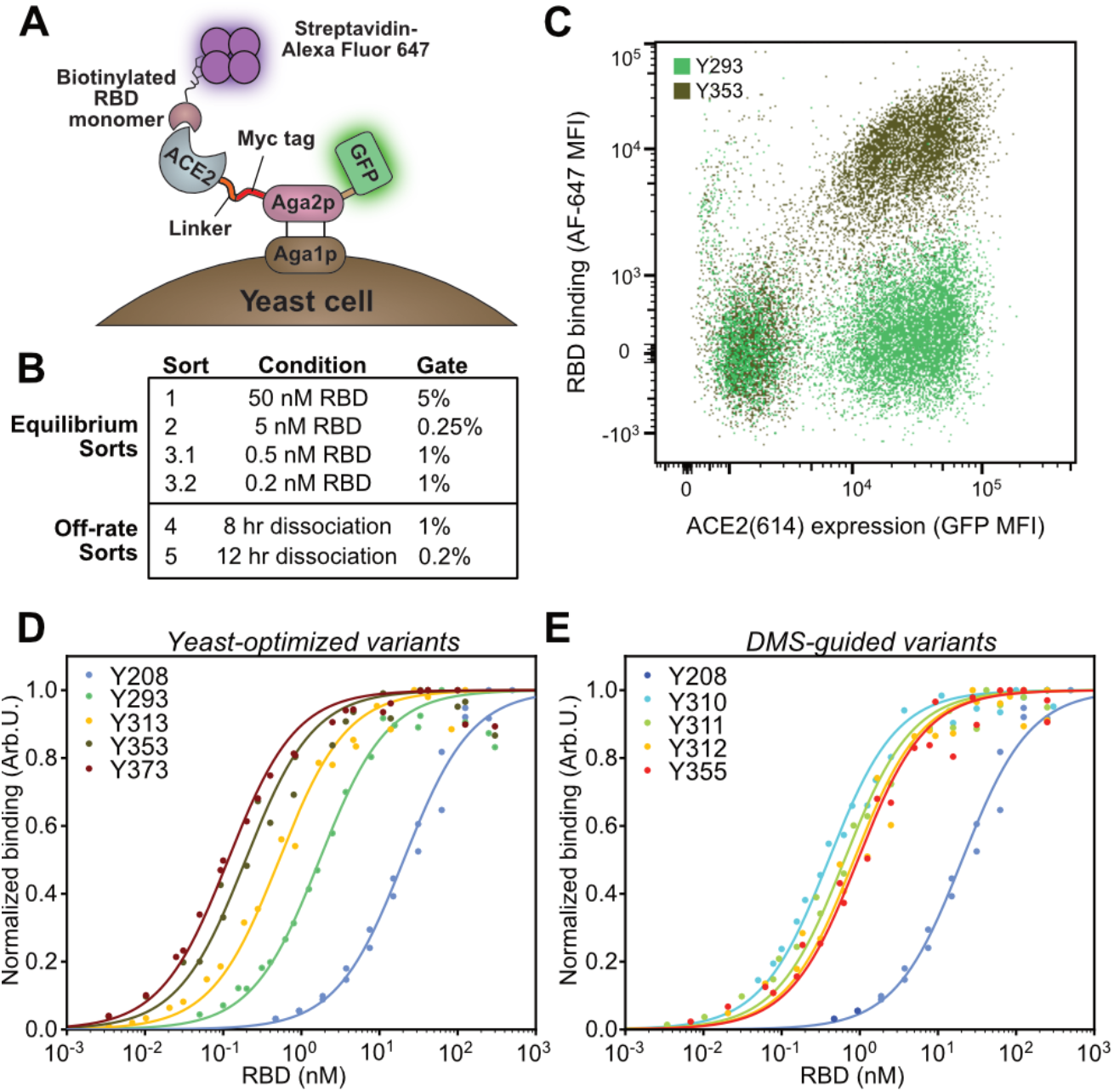
Affinity maturation by yeast surface display of pooled error-prone PCR libraries on designed ACE2 variants and DMS-guided design. **(A)** Monomeric ACE2(614) variants were expressed as Aga2p-GFP fusions on the surface of yeast cells. Binding of ACE2(614) to the spike RBD was quantified by incubation of the cells in a solution of biotinylated RBD, followed by staining the cells with streptavidin Alexa Fluor 647, washing the cells, and measuring fluorescent populations by flow cytometry. **(B)** Stringency was increased in each round of yeast display to enrich the cell population for ACE2(614) variants that bound the RBD tightly. Equilibrium sorts with decreasing RBD concentrations were used for sorts 1-3, and off-rate sorts with increasing dissociation times and decreasing gate size were used for the last two sorts. **(C)** Representative cell populations by flow cytometry for yeast expressing a tight-binding ACE2(614) variant from sort 4 compared to yeast expressing the computationally designed parent, K31F/H34I/E35Q ACE2(614), incubated with 0.1 nM RBD monomer. The additional mutations in the sort 4 variant shift the cell population higher on the *y*-axis due to tighter RBD binding. **(D)** On-yeast titration curves for cells expressing ACE2(614) variants chosen from sorts 3 (Y313), 4 (Y353) and 5 (Y373) that bind the RBD most tightly, compared to WT ACE2(614) (Y208) and K31F/H34I/E35Q ACE2(614) (Y293). **(E)** On-yeast titration curves for cells expressing the ACE2(614) variants generated by DMS-guided design, compared to WT ACE2(614) (Y208). Titration curves in **(D)** and **(E)** are fit to biological duplicates, shown as points. For variant names, mutations, and K_D,app_ values, see Table S3.

Sixteen sublibraries of ACE2 residues 18-103 were made by homologous recombination into ACE2(614) using the four input templates, each mutagenized at four different rates using dNTP analogues (30). After transformation into EBY100 cells for a total library size of 2.7 × 10^7^ members, sequencing of 24 pre-sort clones showed an even distribution of mutations across residues 18-103 with representation from all four input sequences (Figure S3C, D). We carried out sorts of increasing stringency using different concentrations of RBD monomer as outlined in Figures 3B and S4, and analyzed individual clones along the way. Sort 3 was performed with multiple binding stringencies and expression temperatures. For example, sorts of ACE2 induced at 30 °C did not show increased expression in subsequent rounds, but high affinity clones were observed from both 500 pM (for sort 3.1) and 200 pM (for sort 3.2) equilibrium sorts. The sequences of 21 clones from an 8 hour off-rate sort 4 did not show clonal convergence, but were enriched in favorable mutations in agreement with published DMS data (28). A very stringent fifth sort, in which surface-displayed ACE2 were allowed to dissociate from RBD for 12 hours at room temperature and only 0.2% of the cell population were collected, still did not result in sequence convergence, but showed enrichment of one clone derived from the computationally designed K31F/H34I/E35Q ACE2(614) variant. The enriched ACE2(614) variant had the following seven mutations: Q18R, K31F, N33D, H34S, E35Q, W69R, and Q76R. Interestingly, a majority of sequences from sorts 4 and 5 were derived from the K31F/H34I/E35Q ACE2(614) parent, but many of these had mutations at I34 to serine or alanine (Figure S5). This observation suggested that while I34 led to an alternate design variant, the isoleucine was not always the ideal amino acid at this position. A small number of additional mutations appeared in variants originating from different parents, including F40L/S, N49D/S, M62T/I/V, and Q101R, while others appeared only in the K31F/H34I/E35Q background, such as L79P/F and L91P.

Following sorts 3 through 5, 18-24 individual yeast clones were picked from each sort for further characterization. After growth and induction, we analyzed each population for high-affinity mutants by staining with decreasing concentrations of the RBD monomer. The best mutants from each sort were sequenced (mutations listed in Table S3) and their K_D,app_ values for the monomeric spike RBD were measured by on-yeast titrations (Figure 3D, Table S3). The best-characterized mutants from sorts 3.1, 3.2, 4, and 5 had affinities of 0.52 nM, 0.45 nM, 0.19 nM, and 0.12 nM, respectively (between 39- and 170-fold higher affinity than WT ACE2(614)). Though each sort contained a variety of mutants, the highest-affinity clones contained N33D and H34S mutations and were derived from the K31F/H34I/E35Q ACE2(614) variant. The low likelihood of multiple base mutations in a single codon in error-prone PCR mutagenesis likely favored the I34S mutation from this background; interestingly, ACE2 receptors from pangolin species that are hypothesized to be SARS-CoV-2 reservoirs also include a serine at position 34 (31, 32). ACE2 N33 does not directly contact the RBD in the crystal structure, but the enrichment of the N33D mutation in our affinity maturation was consistent with the enrichment of this point mutant in the DMS study (28).

As an orthogonal approach to generate affinity-enhanced ACE2 variants, we leveraged the results from the DMS experiment by Procko (28) to perform an additional round of DMS-guided computational design. Our original computational design strategy (Figure 2A-C) targeted alanine scan hotspots, as described. Procko’s DMS experiment identified beneficial ACE2 point mutations in the ACE2-RBD interface at non-hotspot positions, which could improve binding affinity by direct interactions with the RBD, as well as outside the binding interface, which might serve a stabilizing role. These two classes of mutations would not have been predicted by our computational design strategy. Thus, we performed another round of computational saturation mutagenesis at non-hotspot ACE2 positions in the ACE2-RBD interface (A25, T27, K31), as well as the non-interface residue W69, to predict additional mutations. We generated a set of DMS-guided, computationally designed ACE2 variants with 3-4 mutations each that included at least two mutations outside the interface chosen directly from the DMS dataset, combined with 1-2 mutations from the computational saturation mutagenesis at non-hotspot positions that were also enriched in DMS (Figure S6, Table S3). These designed proteins were displayed on the surface of yeast as ACE2(614)-Aga2p fusions. We measured the K_D,app_ of the DMS-guided ACE2(614) variants for the monomeric RBD to be between 0.4 and 1 nM by on-yeast titrations, which is between 21 and 51-fold higher than WT ACE2(614) (Figure 3E, Table S3). The ACE2(614) variant from DMS-guided computational design with the lowest K_D,app_ had the following mutations: A25V, T27Y, H34A, and F40D.

We next characterized binding of WT, computationally designed, and affinity-matured ACE2 variants in different Fc fusion formats. Using BLI, we tested whether inclusion of the natural C-terminal ACE2 collectrin domain (residues 615-740) could improve the protein’s binding affinity for spike (Figure 4A). A recent cryo-EM structure shows ACE2 as a dimer, with the collectrin domain connecting the extracellular peptidase domain of ACE2 to its transmembrane helix (33). The structure also reveals additional inter-collectrin domain contacts and C-terminal contacts between peptidase domains (Figure S7). Fc fusions of WT ACE2 containing the collectrin domain were also recently shown to be more effective in blocking viral infection (27), perhaps by repositioning the ACE2 monomers for improved binding to spike. Furthermore, the three RBDs in the spike trimer can independently bind ACE2 (34, 35) (Figure 4A). We hypothesized that the inclusion of the ACE2 collectrin domain and the additional two spike RBDs would increase the strength of the ACE2-spike interaction through stabilization and avidity effects. We tested binding of ACE2-Fc with the collectrin domain (ACE2(740)-Fc) to spike RBD and full-length spike (FL spike) using BLI. Compared to the ACE2(614)-Fc interaction with the spike RBD, we indeed saw a 3.7-fold decrease in the monovalent K_D_ of WT ACE2(740)-Fc for the spike RBD, and a dramatic decrease in the K_D_ of ACE2-Fc with or without the collectrin domain for FL spike (Figure 4B, Figure S8A-E). Both WT and computationally designed ACE2(740)-Fc binding interactions with the FL spike were too tight to be accurately measured by BLI due to the massively decreased off-rates (Figure 4B, Figure S8A-E). ACE2(740)-Fc variants from DMS-guided design and affinity maturation in yeast also had greatly reduced off-rates for monovalent interactions with the RBD (Figure S8F, G). The highest-affinity mutants from our yeast display campaign were poorly expressed as Fc fusions, indicating that despite numerous reports of stabilizing mutations from yeast display, ACE2 variant expression on yeast did not translate to well-folded soluble protein (36, 37).

**Figure 4.**
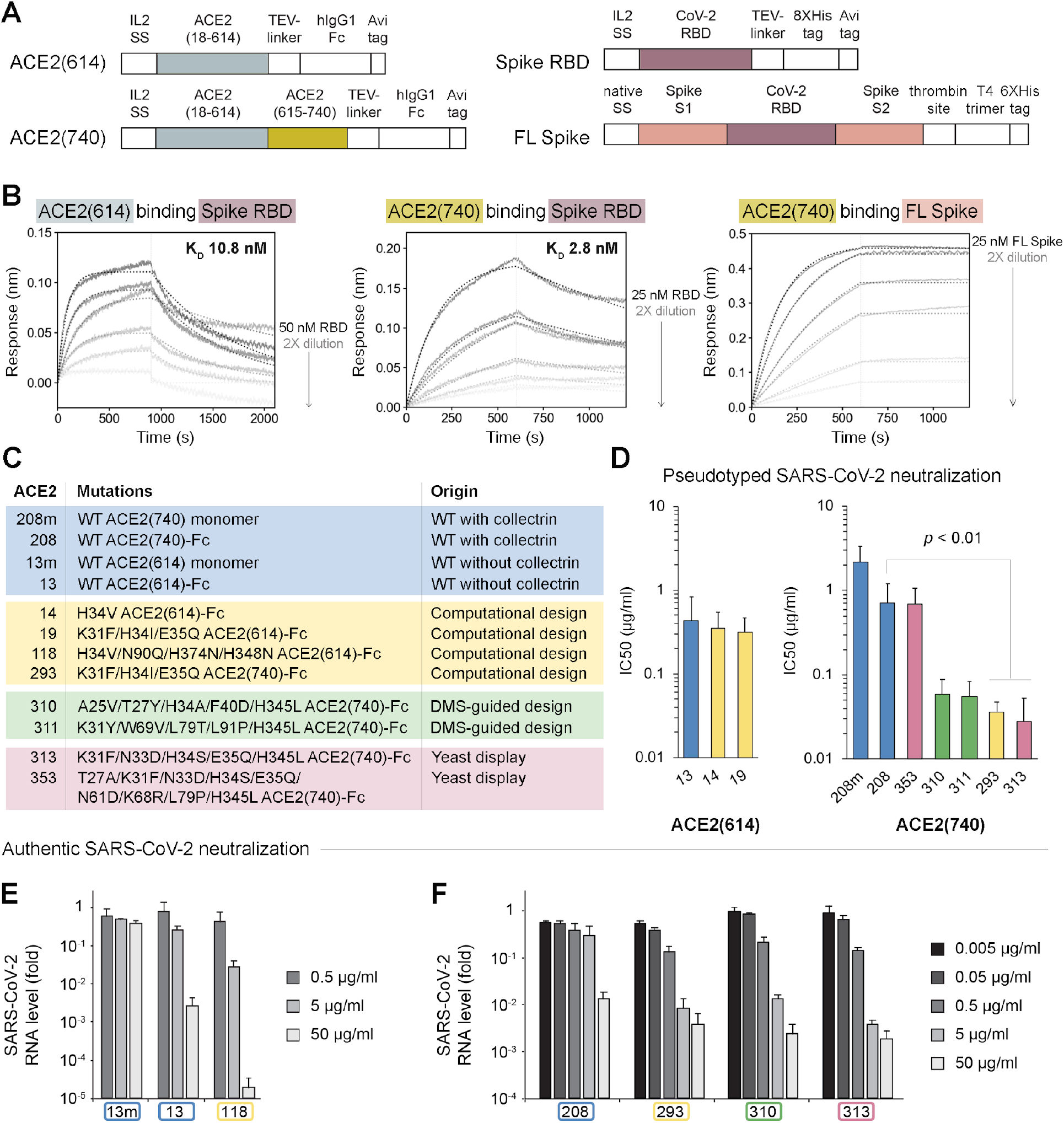
Increased stability, affinity, and avidity effects result in tighter binding between ACE2(740)-Fc and full-length spike and potent viral neutralization. **(A)** Plasmid constructs for expression of ACE2(614) and ACE2(740) *(left)* and spike, in the monomeric RBD and full-length (FL) forms *(right)*. Abbreviations: IL2 SS, IL2 signal sequence (cleaved); TEV-linker, TEV protease cleavage sequence and glycine-serine linker; hIgG1-Fc, human IgG1 Fc domain; Avi tag, target sequence for intracellular biotinylation by BirA; 8X- or 6XHis tag, polyhistidine tag; native SS, native spike signal sequence; thrombin site, thrombin cleavage site; T4 trimer, T4 bacteriophage fibritin trimerization motif. **(B)** BLI measurements show decreased K_D_ for ACE2(740)-Fc as compared to ACE2(614)-Fc for the interaction with the spike RBD. Binding between ACE2-Fc and the full-length spike protein results in a K_D_ less than 100 pM due to very low off-rates. Solid lines show response curves for two-fold dilution titration spanning 0.37-50 nM RBD. Dotted lines show calculated fits. **(C)** Table of all ACE2 variants with scaffolds and mutations tested in pseudotyped and authentic SARS-CoV-2 viral neutralization assays, listed with their origin. **(D)** Fc-fusion, inclusion of the collectrin domain, and affinity-enhancing mutations improve neutralization of ACE2 constructs against pseudotyped SARS-CoV-2 virus, except for misfolded variant 353. Error bars represent standard deviations over all technical replicates from 2-4 biological replicates. Biological replicates were separate experiments using different preparations of the ACE2 variant and pseudovirus, each with two or four technical replicates. Statistical significance with *p* < 0.01 was determined using a homoscedastic two-tailed t-test. Authentic SARS-CoV-2 viral neutralization experiments with ACE2 variants in VeroE6 cells showed that **(E)** Fc fusion (CVD013) and addition of computationally-predicted mutations (CVD118) enhance neutralization by greater than 50,000-fold over a control anti-GFP IgG antibody sample, and **(F)** inclusion of the collectrin domain, computationally predicted mutations (CVD293), DMS-guided mutations (CVD310), and mutations from affinity maturation in yeast (CVD313) enhance neutralization potency over a control anti-GFP IgG antibody sample to IC50 values of 73-136 ng/ml. **(E)** and **(F)** show results from different experiments in biological duplicate. Error bars represent the standard error of the mean.

The soluble domain of ACE2 converts angiotensin II to angiotensin(1-7), a vasodilator, and was shown to be safe in clinical trials (12, 13). The RBD binds outside the enzyme active site. We inactivated the peptidase activity of ACE2 to avoid off-target vasodilation effects without affecting the binding affinity for the RBD (Figures 2D, E, S2) (11). Our original inactivation mutations, H374N and H378N, ablated Zn^2+^ binding. However, protein stability is also an important factor to consider in engineering an optimal ACE2-based therapeutic scaffold. Incorporation of the ACE2 collectrin domain in our Fc-fused constructs improved the apparent melting temperature of the ACE2-Fc variants as measured by circular dichroism spectroscopy, but the H374N/H378N enzymatic inactivation mutations were destabilizing (Figure S9). Therefore we adapted the ACE2(740)-Fc scaffold to include the inactivation mutation H345L instead, which is important for substrate binding and is not destabilizing (Figure S2, Figure S9) (38). H345L ACE2(740)-Fc does not have detectable peptidase activity in an activity assay and does not impact binding to the spike RBD in a BLI assay (Figure S10), but maintains the thermal stability of WT ACE2.

We found that the binding affinity improvements to ACE2 were robust to the method of measurement (BLI vs. binding on yeast) and well-correlated with neutralization efficacy. To evaluate their efficacy in neutralizing SARS-CoV-2 infections, several affinity-improved ACE2 variants from computational design, DMS-guided design, and yeast display were expressed in the H345L ACE2(740)-Fc format, purified, and assayed for viral neutralization against pseudotyped lentivirus and authentic SARS-CoV-2 (Figure 4C-F). In the pseudotyped viral neutralization assay in ACE2-expressing HEK cells, different ACE2(740)-Fc molecules with mutations derived from computational design, DMS-guided design, and affinity maturation using yeast surface display neutralized SARS-CoV-2 with IC50 values of 58 ng/ml, 55 ng/ml, 36 ng/ml and 28 ng/ml, demonstrating multiple paths to significant improvements in efficacy (Table S4, Figure 4D, see variants 310, 311, 293, and 313, respectively). We also confirmed that the affinity-enhanced ACE2 variant 353 (T27A/K31F/N33D/H34S/E35Q/N61D/K68R/L79P/H345L ACE2(740)-Fc) does not effectively neutralize SARS-CoV-2, despite this molecule’s enrichment in the yeast display campaign and low K_D,app_, possibly because the variant is unstable or otherwise misfolded (Figure 4D, Figure S11). Taken together, the neutralization data revealed that the mutations to ACE2 from computational design and affinity maturation, addition of the collectrin domain, and fusion to the Fc domain significantly improve neutralization over unmodified ACE2(740). The unmodified ACE2(740) (Figure 4C, D, variant 208m) is similar to the molecule APN01 that is currently in clinical trials for treating SARS-CoV-2 infections (Figure 4D, Figure S11, Table S4) (*16, 17*).

Data from viral neutralization assays in which *bona fide* SARS-CoV-2 was used to infect VeroE6 cells in a biosafety level 3 facility closely reflected the results from the pseudotyped viral neutralization assays. We determined that fusion of the Fc domain to ACE2(614) improved neutralization by 370-fold over monomeric ACE2(614), and additional inclusion of computationally predicted mutations H34V and N90Q improved neutralization by more than 50,000-fold over an anti-GFP IgG control, at the highest concentration tested (Figure 4E, 50 µg/ml). The IC50 for intermediate this affinity-enhanced ACE2(614)-Fc variant, CVD118, was less than 0.5 µg/ml.

Addition of the ACE2 collectrin domain further improved neutralization potency, with ACE2(740)-Fc variants originating from computational design, DMS-guided design and affinity maturation in yeast demonstrating efficient neutralization in the neutralization assay using *bona fide* SARS-CoV-2 (Figure 4F). WT ACE2(740)-Fc (variant 208), computationally-designed variant 293, DMS-guided design 310, and yeast affinity-matured variant 313 were tested at concentrations starting at 0.05 µg/ml. Variants 293, 310 and 313 each considerably diminished viral RNA levels at concentrations between 0.05 and 50 µg/ml, while WT ACE2(740)-Fc only had neutralization efficacy at 5 µg/ml. Variants 310 and 313 displayed the most neutralization potency, with IC50s of approximately 90 ng/ml and 73 ng/ml, respectively (Figure 4F, Table S4). This neutralization potency is comparable to recently reported antibodies isolated from convalescent COVID-19 patients (6, 39). None of the ACE2 variants induced cytotoxicity in uninfected cells at the concentrations used in the neutralization assay (Figure S12). Additional live SARS-CoV-2 neutralization experiments with shorter incubation times (16 hours rather than 26 hours) and a different SARS-CoV-2 strain were conducted in a different laboratory to ensure reproducibility and measure the effect of the affinity-enhanced ACE2(740)-Fc molecules on viral entry more directly, and yielded similar results: IC50 values were in the range of 0.1-1 µg/ml for variant 293, and lower for variants 310 and 313 (Figure S13).

The inclusion of the ACE2 collectrin domain and the human IgG Fc domain dramatically increased the neutralization potency of the ACE2 variants through improved affinity, stability, and avidity (Figures 4, S2, S7). The Fc fusion results in ACE2 dimerization, but the collectrin domain may serve to position the ACE2 molecules closer together than would be achieved with the Fc alone (33). We also observed that the interaction of dimeric ACE2-Fc with the full-length trimeric spike protein is stronger than with the monomeric RBD due to avidity effects. As a combined result of these effects, the use of the H345L ACE2(740)-Fc scaffold is central to the neutralization potency of the affinity-enhanced variants.

ACE2-based therapeutics could be used to treat other respiratory infections with ACE2-dependent cell entry mechanisms, such as those caused by SARS-CoV-1 and HCoV-NL63 coronaviruses (40, 41). We tested whether WT ACE2(740)-Fc and our most robustly expressed ACE2(740)-Fc variants could bind the SARS-CoV-1 and NL63 spike RBDs (Figure 5). Indeed, WT ACE2(740)-Fc, a receptor trap from affinity maturation in yeast (variant 313, with K31F, N33D, H34S, E35Q, and enzymatic inactivation mutation H345L), and its computationally designed parent (variant 293, K31F, H34I, E35Q) bound with nanomolar K_D_ to the SARS-CoV-1 RBD (Figure 5D-F) and tens of nanomolar K_D_ to the NL63 RBD (Figure 5G-I), which is close to previous observations for the NL63 RBD-WT ACE2 interaction (40). The weaker binding affinity for the NL63 RBD interaction with ACE2-Fc is likely due to its low structure/sequence similarity to the SARS-CoV-2 RBD, while the SARS-CoV-1 and SARS-CoV-2 RBDs have similar structures and 73% sequence identity (2) (Figure S14). In contrast, the ACE2 variants did not bind appreciably to the Middle East respiratory syndrome (MERS) RBD up to 150 nM (Figure S15); MERS-CoV particles enter cells not via ACE2 but through interactions with the dipeptidyl pepdidase IV (DPP4, also known as CD26) membrane protein (42).

**Figure 5.**
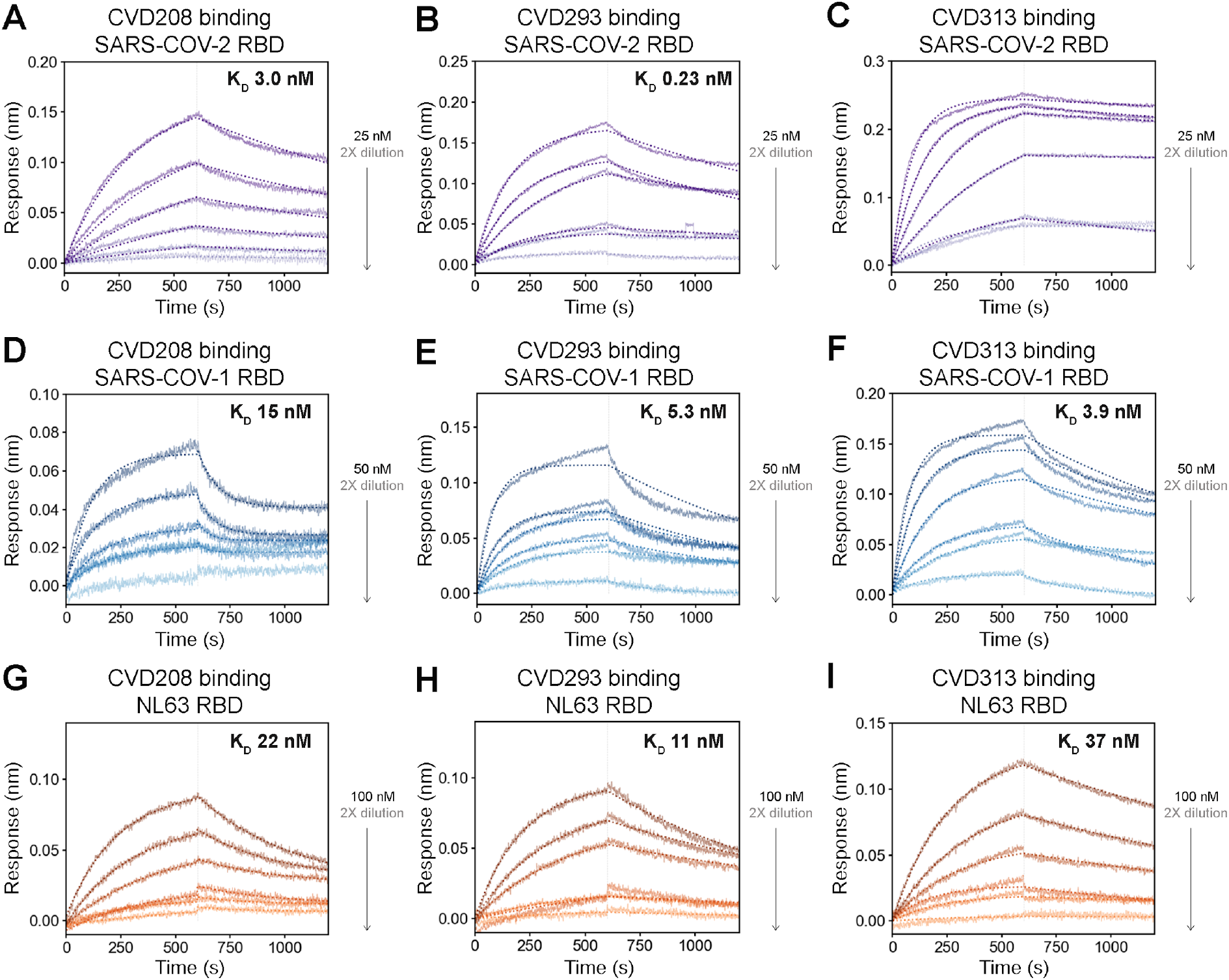
WT and engineered ACE2(740)-Fc binds the SARS-CoV-1 RBD and the HCoV-NL63. Representative BLI measurements for **(A, D, G)** WT ACE2(740)-Fc, **(B, E, H)** K31F/H34I/E35Q ACE2(740)-Fc, and **(C, F, I)** K31F/N33D/H34S/E35Q/H345L ACE2(740)-Fc interactions with the SARS-CoV-2 RBD **(A-C)**, the SARS-CoV-1 RBD **(D-F)**, and the HCoV-NL63 RBD **(G-I)** at concentrations of 0.375 nM to 100 nM RBD, with the highest RBD concentration tested and K_D_ values as indicated. Dotted lines show calculated fits. The extremely slow off-rate observed in **(C)** precluded K_D_ determination.

## Discussion

Engineered receptor traps have been used extensively as biotherapeutics for binding vascular endothelial growth factor, tumor necrosis factor alpha, and other cytokines and produced at therapeutic scale (30, 43–45). Affinity is often sufficiently improved by simply presenting these proteins in dimeric form fused to human engineered Fc to achieve avidity and afford long half-lives. There are no FDA-approved examples of receptor traps as antiviral biologics (46, 47), though some have entered clinical trials for HIV treatment (48). Here we show it is possible using both computational design and selection methods to dramatically improve the binding for dimeric ACE2 for the SARS-CoV-2 spike RBD to the range of high affinity antibodies.

We used several computational design methods in Rosetta to predict mutations that could enhance the affinity of ACE2 for the SARS-CoV-2 RBD, with one innovation: following computational saturation mutagenesis, we proceeded to the local redesign step with the lowest-energy point mutant (H34V) as well as the highest-energy point mutant (H34I). Our hypothesis was that mutating and optimizing the positions of surrounding residues would allow us to identify alternative low-energy solutions for the ACE2-RBD interaction, which proved to be the case. We experimentally validated that the variant designed from H34I ACE2 improved binding affinity, pseudotyped viral neutralization, and authentic viral neutralization in two independent laboratories as compared to other computationally predicted mutations, despite the fact that no single mutation in this design conferred a fitness advantage in DMS (Figures 2, 4, Table S1) (28). Furthermore, the same ACE2 variant was the parent of the most effective ACE2 molecules to emerge from affinity maturation (Table S3, Figure S5). This demonstrates the potential benefits of developing computational design methods that include steps to diversify solutions.

Engineered ACE2 receptor traps present key advantages in treating SARS-CoV-2 infections. Several groups have isolated antibodies from convalescent patients, confirmed their neutralization potencies, characterized their affinities to the RBD, and in some cases, determined their structures. A subset of the neutralizing antibodies block ACE2 while the remainder bind spike epitopes outside the ACE2-binding interface (3, 4, 6–10, 18–20, 39). By contrast, ACE2 receptor traps directly compete with the essential viral entry mechanism. Recent reports indicate that RBD-binding antibodies are also susceptible to diminished affinity at lower pH, which could lead to lower viral neutralization and potentially re-infection (35). Finally, viral escape mutations can render antibody therapeutics ineffective, but escape mutations that reduce the efficacy of an ACE2 receptor trap binding are also likely to reduce viral invasion. Many of the affinity-enhancing mutations that we described are in the receptor binding site and so it is conceivable that they could be selectively targeted by a mutant virus, but we show that even the NL63 RBD can still bind our ACE2 variants despite its significant sequence and structure divergence from the SARS-CoV-2 RBD. Receptor traps based on wild-type ACE2 (11, 49), antibody fusions (50), or affinity enhanced mutants (27), which include naturally occurring mutations that were also found in our engineering efforts (31), have also been reported to neutralize pseudovirus and spike-based cell fusion. Our work harnesses the power of protein engineering approaches to build on these lines of research by engineering orders of magnitude higher affinity and demonstrating potent neutralization of authentic SARS-CoV-2 virus. We also anticipate that engineered ACE2 receptor traps could synergize in a cocktail with neutralizing antibodies that bind the RBD outside the ACE2 binding site to treat viral infections (20).

The systematic two-pronged affinity optimization approach for engineering ACE2 receptor traps was achieved by a small team in several months, which is comparable to antibody isolation from convalescent patients or selection by *in vitro* methods (10). Thus, it represents a rapid and orthogonal approach to generating therapeutic candidates for treating future viral pandemics, without any prerequisite for an infected human population. Engineered receptor traps can be stockpiled as potentially useful drug candidates for multiple viruses that use the same port of entry as we show for NL63 and pandemic viruses SARS-CoV-1 and CoV-2 and decrease the likelihood of viral resistance. Our study on ACE2 provides a systematic road map to redesigning an entry receptor as a therapeutic, and we believe the same strategy could be applied to other entry receptors such as DPP4 for treating MERS and ANPEP for treating upper respiratory tract infections by HCoV-229E (42, 51).

## Acknowledgements

We wish to thank Jamie Byrnes, Susanna Elledge, the entire COVID team in the Wells lab, and members of the Kortemme lab for helpful discussions and support. We gratefully acknowledge the Chan Zuckerberg Biohub scientists for plasmids to express SARS-CoV-1 and HCoV-NL63 RBDs and the Florian Krammer lab for plasmids to express the full-length SARS-CoV-2 spike protein. Cell sorting was performed at the UCSF Laboratory for Cell Analysis which is supported by a National Cancer Institute Cancer Center Support Grant (P30CA082103). We thank Darryl Falzarano of the Vaccine and Infectious Disease Organization-International Vaccine Centre, University of Saskatchewan, for providing the SARS-CoV-2 strain for this work. We also valued helpful discussions with Dr. Diane Barber at UCSF and Drs. Ho Cho, Hari Hariharan, and Henry Chan at Bristol Myers Squibb. AG was supported by NIH 1K99GM135529. IL and NJK were supported by NSF Graduate Research Fellowships. XXZ was supported by a Merck Fellowship from the Damon Runyon Cancer Research Foundation (DRG-2297-17). SAL is a Merck Fellow of the Helen Hay Whitney Foundation. Research in the JVR lab was supported by Rockefeller University and NIH U19AI111825. APW was supported by NIH DP2 OD022552. Research in the TCH lab was supported by the Canadian Institutes of Health Research and the Li Ka Shing Institute of Virology. TK was supported by funding from the Chan Zuckerberg Biohub Investigator Program and a COVID-19 grant from the UCSF Program for Breakthrough Biomedical Research (PBBR). JAW was supported by generous funding from the Chan Zuckerberg Biohub Investigator Program, the Harry and Dianna Hind Professorship and the Harrington Foundation.

## Methods

### Structural modeling and computational protein design

See “Supplemental computational methods” section in the Supplemental Information file for command lines and input files for each section. We used the 2019.38 release of Rosetta 3.12 (Git SHA1 hash number: 2019.38.post.dev+231.master.04d3e581085 04d3e581085629b0f0c46f1e1aef9e61978e0eeb).

#### Preparation of ACE2-spike structure for modeling

To model ACE2-spike interactions, we used the 2.50 Å resolution X-ray structure of the spike receptor binding domain complexed with the soluble extracellular domain of ACE2 as determined by Wang *et al*. (PDB 6LZG) (23). This structure was downloaded from the PDB, relaxed with coordinate constraints on backbone and sidechain heavy atoms, and minimized in Rosetta without constraints using default options using the beta_nov16 score function (see command lines in the “Supplemental computational methods” section).

#### Identification of ACE2 residues that contribute to binding in the ACE2-spike interface and are chosen for design

To determine which residues contribute most strongly to the ACE2-spike interaction, we used the Robetta Computational Interface Alanine Scanning Server (24, 25) (publicly available at http://robetta.bakerlab.org/alascansubmit.jsp) to perform a computational alanine scan on the relaxed and minimized input structure of the complex. The alanine scan identified 18 ACE2 residues in the interface, six of which had DDG(complex) values greater than 1. These “hotspots” represent amino acid side chains that are predicted to significantly destabilize the interface when mutated to alanine. See Table S1 for the full results of computational alanine scanning.

We used two metrics to determine which hotspots could most likely be mutated to improve the ACE2-spike binding affinity: (1) the total per-residue energy as evaluated by the Rosetta score function to be the sum of all one-body and half the sum of all two-body energies for that residue, and (2) the total contribution of the residue to the interface energy, which is the sum of pairwise residue energies over all residue pairs (R1, R2) where R1 belongs to ACE2 and R2 belongs to the spike RBD. We classified hotspot residues that had total residue energies in the top 30% of all residues in ACE2 as well as total cross-interface interaction energies greater than 0.5 REU as residues in the ACE2-spike interface to be targeted for design. These residues were H34, Q42, and K353.

#### Computational saturation mutagenesis at targeted ACE2 interface residue positions

We systematically computationally mutated H34, Q42 and K353 in ACE2 to every other amino acid except cysteine, allowing all residues with sidechain heavy atoms in ACE2 or spike within 6 Å of the mutated position to repack (change rotameric conformation), minimized the entire complex, and recalculated all of the pairwise interaction energies across the ACE2-spike interface and various interface metrics. These interface metrics were: the solvent-accessible surface area buried at the interface; the change in energy when ACE2 and spike RBD are separated vs. when they are complexed; the energy of separated chains per unit interface area; the number of buried and unsatisfied hydrogen bonds at the interface; a packing statistic score for the interface; the binding energy of the interface calculated with cross-interface energy terms; a binding energy calculated using Rosetta’s centroid mode and score3 score function; the number of residues at the interface; the average energy of each residue at the interface; the energy of each side of the interface; the average per-residue energy for each side of the interface; the average energy of a residue in the complex; the total number of cross-interface hydrogen bonds; and the interface energy from cross-interface hydrogen bonds. Each point mutation was modeled five times using this protocol, and the lowest of the summed cross-interface pairwise interaction energies from the five trials was used for comparison to the wild-type interface value.

#### Redesign of ACE2 interface residues incorporating H34V or H34I mutations

We generated two additional sets of models to select constructs for experimental testing that incorporated the most and least energetically favorable point mutations to H34 from the computational saturation mutagenesis. In these simulations, we first mutated H34 to either valine or isoleucine. Residues within 6 Å of the interface were repacked, and minimization was applied to the interface backbone and sidechain torsion angles. A flexible backbone design algorithm (Coupled Moves) (26) was applied to allow neighboring ACE2 residues 30, 31, 35, and 38 to change amino acid identities while allowing ACE2 residues 29, 32, 33, 34, 36, and 37 and RBD residues 416, 417, 418, 452, 453, 455, 456, 492, 493, 494 to repack. Changes in the positions of backbone atoms were allowed for ACE2 residues 30-38 and RBD residues 417, 453-455, and 493. The whole complex in the lowest-energy solution for the redesigned interface was again repacked and minimized, and the final structure was scored. The lowest of the summed cross-interface pairwise interaction energies from 20 trials was used.

### Generation of plasmids, strains, and proteins

#### Cloning

SARS-CoV-2 spike RBD and ACE2 variants were cloned into a pFUSE-based vector for mammalian expression using the Gibson method, transformed into XL10-Gold cells, and grown on low-salt LB + 25 µg/ml Zeocin. The genes encoding the SARS-CoV-2 RBD (328-533), ACE2(18-614) or ACE2(18-740) were inserted between two SpeI sites with a 5’ mutated IL-2 signal sequence for secretion and 3’ sequences for Gly-Ser linker, TEV protease cut site, human IgG1 hinge and Fc, and AviTag. The RBD monomer was also cloned into a similar construct where the Fc domain was replaced with an 8XHis tag. ACE2 point mutations were made by stitching 5’ and 3’ fragments encoding the desired mutation by PCR and inserting the resulting gene into ACE2 vectors digested with KasI and BsiWI with the Gibson method. Wild-type ACE2(18-614) was inserted into NheI/BamHI-double digested pCL2 as follows: fragments encoding amino acids 18-105 and 106-614 were PCR-amplified with 25-bp Gibson overlap regions introducing a silent mutation at S105 to introduce a new BamHI site, and a 3’ silent mutation to eliminate the BamHI site in pCL2. These fragments were stitched together by PCR and inserted into pCL2, transformed into XL10-Gold, and grown on LB + 50 µg/ml carbenicillin. ACE2 variants in pCL2 were exchanged by digesting this vector with NheI/BamHI. ACE2 variants from yeast display were amplified and cloned into the ACE2(18-740)-Fc fusion vector in between the BlpI restriction sites on the IL-2 signal peptide and in the ACE2 gene.

#### Transfections

A cell line derived from Expi293 cells expressing an ER-localized biotin ligase (BirA) gene was grown in Expi293 media (Thermofisher Scientific) supplemented with 100 µM biotin (GoldBio) under 8% CO2 at 37 °C and used for all protein expressions. Transfections were carried out in 30 ml media as described in the Expi293 manual: 75 million cells at >98% viability were suspended in 25.5 ml media with 100 µM biotin. Expifectamine (81 µl) and plasmid DNA (30 µg total) were mixed separately with 1.5 ml OptiMEM each, incubated at room temperature for 5 minutes, combined, and incubated for 20 minutes. The mixture was added to the cells, and after 20 hours of growth Enhancers 1 and 2 (150 µl and 1.5 ml, respectively) were added. Proteins were allowed to express for 5 days.

#### Protein purification

Expi293 cells were spun down at 3000 × *g* for 20 minutes, and the supernatant from each culture containing protein of interest was collected, filtered through a 0.22 µm syringe filter and neutralized with 10X phosphate buffered saline (PBS, 0.01 M phosphate buffer, 0.0027 M KCl and 0.137 M NaCl, Millipore Sigma P4417-100TAB), pH 7.4. The supernatants were purified using a peristaltic pump and a HiTrap Protein A column (GE Healthcare). Fc-fused proteins were acid-eluted into 1X PBS, pH 7.4, from protein A columns, and then buffer-exchanged into 1X PBS, pH 7.4, using spin concentration columns (Millipore Sigma). Later ACE2(740)-Fc constructs were eluted with 50 mM Tris pH 7.2, 4 M MgCl_2_ and similarly buffer exchanged. Biotinylation was quantified by denaturing proteins at 0.1 mg/ml in Lamelli buffer with 5 mM DTT for 5 minutes at 95 °C, followed by addition of a molar excess of avidin and SDS-PAGE. Proteins were >95% pure and >95% biotinylated as determined by ImageJ analysis.

### Biophysical characterization of ACE2 mutants

#### Determination of binding affinity to spike using bio-layer interferometry (BLI)

In our BLI experiments, the biotinylated ACE2 variant is tethered to an optically transparent biosensor tip by a biotin-streptavidin interaction, and spike is present as the analyte in solution in the microplate. ACE2 gene sequences and mutations are listed in Appendix 1. Affinity measurements were carried out at room temperature using an Octet RED96 system and streptavidin (SA)-coated biosensor tips (Pall ForteBio). Biotinylated ACE2 variants were diluted to 10 nM in PBS with 0.2% BSA and 0.05% Tween-20 (PBS-T), pH 7.4 to be used as the antigen. Antigen-bound SA-tips were washed in in PBS-T pH 7.4, separately exposed to the spike solutions at concentrations ranging from 0 to 50 nM spike in the same buffer during an association period, and then returned to the washing well during a dissociation period. The binding protocol was as follows: rinse tips in PBS-T, 60 seconds; load tips with antigen, 180 seconds; establish baseline by rinsing tips in PBS-T buffer, 180 seconds; association with analyte, 600 seconds; dissociation in baseline wells, 900 seconds. Raw data was fit to 1:1 binding curves in Octet Data Analysis HT software version 10.0 using curve fitting kinetic analysis with global fitting. The theoretical equilibrium binding signal response data (R equilibrium) were normalized by the steady-state group maximum response (R_max_) values, and the steady state affinity was determined using the Hill equation. Non-cooperative binding kinetics were assumed. All fits to BLI data had R^2^ (goodness of fit) > 0.90.

#### Determination of stability by circular dichroism (CD) spectroscopy with thermal denaturation

CD data were collected on a Jasco J-710 spectrometer using purified ACE2 variant solutions in 1 mm quartz cuvettes. ACE2 variants were diluted in 300 µl PBS, pH 7.4 to concentrations ranging from 2 to 3 µM. Melting curves at 225 nm were measured by increasing the temperature from 25 °C to 80 °C using a rate of 1 °C per minute. CD spectra from 200 to 280 nm were measured at 25 °C and 80 °C. Melting curve data were normalized using an average of the before-melt baseline as 0% and an average of the after-melt baseline as 100%, and the apparent melting temperature, T_m,app_ was determined to be the temperature at which 50% of the protein was denatured between these points. Melting was irreversible for WT ACE2.

#### ACE2 proteolytic activity assay

Hydrolysis of (7-methoxycoumarin-4-yl)acetyl-Ala-Pro-Lys(2,4-dinitrophenyl)-OH (Mca-APK-DNP, Enzo Life Sciences) was used to quantify ACE2 peptidase activity. 50 µl each of solutions of ACE2 variants (diluted to 0.3 nM) and 100 µM Mca-APK-DNP in 50 mM MES buffer, 1 M NaCl, and 10 µM ZnCl_2_ were mixed in a 96-well plate. Fluorescence increase over time was monitored (320 ex./405 em.).

#### Library preparation

ACE2 H34V, N90Q, H34V/N90Q, and K31F/H34I/E35Q were cloned into pCL2 as N-terminal fusions to Aga2p followed by eGFP. A silent BamHI site was incorporated at G104/S105, and the BamHI site at the C-terminal linker in pCL2 was deleted for downstream library generation. 1 ng was used as template for initial error-prone PCR using 8-oxo-GTP and dPTP (TriLink Biotechnologies). Four 50 µl PCR reactions were performed on each template with increasing concentrations of mutagenic nucleotides (5 µM, 10 µM, 50 µM, and 100 µM). The 5 and 10 µM reactions were carried out using Taq polymerase in standard buffer under the following conditions: Initial denaturation at 95 °C for 30 seconds, 20 cycles of PCR (95 °C for 20 seconds, 55 °C for 20 sec, 68 °C for 45 sec), and final elongation at 68 °C for 300 sec. These reactions were loaded onto a 2% agarose gel, and the resulting bands at ∼330 bp were excised and purified. The 50 and 100 µM mutagenic PCRs were carried out similarly except with 5 cycles of PCR followed by DpnI digestion of the template for several hours at 37 °C. After heat inactivation of the DpnI, 5 µl of crude PCR reaction mixture were used as template for a standard Phusion PCR and gel purified as above. To generate enough DNA for yeast transformation 150 ng from each of these 16 PCRs were used as templates for Phusion PCRs (2 × 50 µl reactions each). After spot-checking several reactions for the expected products by agarose gel, these reactions were pooled into 4 tubes and ethanol precipitated with 0.1 volumes of 3 M sodium acetate pH 5.2, 0.1 µg glycogen, and 3 volumes of ethanol. These were incubated several hours at room temperature followed by overnight at −80 °C and centrifuged for 20 minutes at 16,000 × g. Pellets were washed with cold 70% ethanol and centrifuged again. After removing the supernatant, the pellets were air-dried and suspended in 20 µl of sterile water and centrifuged again.

#### Yeast transformations

Electrocompetent EBY100 were prepared by the method of Benatuil *et al*. (52). Briefly, a stationary phase culture of EBY100 was subcultured to an OD600 of 0.3 in 200 ml YPD media and grown at 30 °C with shaking at 250 rpm for 4.5 hours. Upon reaching OD600 = 1.6, cells were centrifuged at 3000 × g for 3 minutes, washed twice with 100 ml ice-cold MilliQ water and once with 100 ml ice-cold electroporation buffer (1 M sorbitol, 1 mM CaCl2). Cells were resuspended in 50 ml 0.1 M lithium acetate/0.01 M DTT and incubated at 30 °C for 30 minutes with shaking at 250 rpm. Cells were pelleted and washed once more with 100 ml electroporation buffer and resuspended in a minimal volume of electroporation buffer. Sublibraries prepared above were pooled into four separate electroporation cuvettes by parental ACE2 variant. A total of 30 µg of each pool was mixed with 10 µg pCL2-ACE2(18-614)BamHI-Aga2-sfGFP previously digested with NheI-HF and BamHI-HF and mixed with 400 µl electrocompetent EBY100 and incubated on ice for 5 minutes. Cells were electroporated using a Biorad Gene Pulser Xcell with an exponential pulse (2.5 kV and 25 μF), pooled, and recovered in 40 ml of 1:1 YPD:electroporation buffer at 30 °C with shaking at 250 rpm. After 2 hours the cells were centrifuged and resuspended in SDCAA and diluted up to 500 ml in SDCAA. Serial dilutions starting at 1/100 were plated on SDCAA agar plates. After 4 days at 30 °C, 56 colonies were counted on a 1/100 dilution plate for a total library size of 2.8 × 10^7^.

#### Flow cytometry

Flow cytometry analysis of individual ACE2-expressing yeast clones was carried out using a Beckman Coulter Cytoflex flow cytometer. Approximately 50,000 cells from an overnight SGCAA culture were pelleted by centrifugation at 3,000 × g for 3 minutes and washed in HyClone PBS + 3% BSA (PBSA). Cells were resuspended in 100 µl PBSA, and appropriate volumes of biotinylated RBD monomer were added to avoid ligand depletion (estimating 50,000 copies of ACE2 per cell). These were incubated 2-4 hours at room temperature with rotation to reach equilibrium (53), washed three times with PBSA, and stained for 20 minutes on ice with 1 µg/ml Alexa Fluor 647-conjugated streptavidin (Thermofisher Scientific S21374). After two more washes with HyClone PBS (without BSA) cells were suspended in 200 µl and analyzed. To fit yeast binding data to KDs, Alexa Fluor 647 mean fluorescence intensities were extracted from the GFP-positive population, background subtracted using secondary only controls, normalized to the highest fluorescent population, and fit to the Hill equation without cooperativity in Python.

#### Library screening

Cells were sorted using a BD FACS Aria II. For sort 1, approximately 10^8^ induced library cells were washed twice with PBSA and stained with 50 nM biotinylated RBD monomer (10 ml) for 1 hour at room temperature. After washing 3x with 5 ml ice-cold PBSA, cells were incubated on ice with a 1/1000 dilution of streptavidin Alexa Fluor 647 conjugate. Cells were washed twice with 10 ml PBSA and immediately used to sort binding clones. For sort 1, the top 5% of RBD binders from 5 × 10^7^ cells were sorted into 3 ml SDCAA. These were pelleted at 3000 × g for 3 minutes and used to inoculate 50 ml in SDCAA. After overnight growth at 30 °C, cells were spun down and used to start a 50 ml, OD = 1 culture in SGCAA. For sort 2, 10^7^ cells were washed and stained with 10 ml of 5 nM biotinylated RBD monomer and processed similarly to round 1. The top 0.25% of RBD binders were sorted into 3 ml SDCAA, diluted to a 5 ml culture in SDCAA, and grown at 30 °C overnight. For sorts 3.1 and 3.2, cells from sort 2 were subcultured (starting OD = 1 in SGCAA) and induced at 20 °C overnight. 10^7^ cells were washed with PBSA stained with 500 pM or 200 pM RBD monomer (80 ml) for 4 hours at RT followed by washing and secondary staining as above. Approximately 6 × 10^6^ cells were analyzed and the top 1% were collected and cultured in 20 ml SDCAA. For sort 4, cells from sort 3 were subcultured as above. 5 × 10^6^ cells were washed in PBSA and stained with 5 ml 5 nM RBD monomer for 30 minutes at RT, followed by 3 washes with PBSA. Cells were resuspended in 10 ml PBSA with 20 nM H34V-ACE2(614)-Fc and incubated at RT for 8 hours with rotation. Cells were washed once and stained with 2 ml 1/1000 streptavidin Alexa Fluor 647 for 20 minutes, washed twice with PBSA and resuspended in 1 ml PBSA. The top 1% of cells were sorted and cultured in 20 ml SDCAA. 100 µl of a 1/100 dilution of the culture was plated on SDCAA agar to isolated individual clones for analysis. Sort 5 was performed similarly to sort 4 but with a 12-hour dissociation with 20 nM soluble H34V-ACE2(614)-Fc as a competitor. The top 0.2% of cells were collected and cultured as above.

#### Analysis of yeast library

To analyze sequences, plasmids were isolated from yeast using a modified version of Singh and Weil (54). 5-10 ml of saturated SDCAA culture were pelleted and resuspended in 200 µl buffer P1. 10 µl of Zymolyase (Zymo Research E1004) or 50 µl Lyticase from *Arthrobacter luteus* (Sigma L4025-25KU) were added and the cells were incubated at 37 °C for 1-2 hours. An equal volume of buffer P2 was added and cells were incubated 10 minutes at RT with gentle mixing. 350 µl buffer N3 was added and the lysate was centrifuged at 16,200 × g for 10 minutes. The supernatant was applied to an EconoSpin miniprep column, washed with 500 µl of buffer PB followed by 750 µl buffer PE, and eluted in 50 µl buffer EB. Library pools were transformed into XL10-Gold competent cells and plated for individual colonies on LB + 50 µg/ml carbenicillin agar plates, while individual clones were amplified directly by PCR.

#### SARS-CoV-2 pseudotyped virus neutralization assay

Pseudotyped reporter virus assays were conducted as previously described (55). Pseudovirus plasmids were a gift from Peter Kim’s lab at Stanford University. HEK-ACE2 cells were cultured in D10 media (DMEM + 1% Pen/Strep + 10% heat-inactivated FBS). Briefly, spike-pseudovirus with a luciferase reporter gene was prepared by transfecting plasmids into HEK-293T cells. After 24 hours, the transfection solution was replaced with D10 media and the virus was propagated for 48 hours before harvest and filtration of supernatants. To titer each virus batch, HEK-ACE2 were seeded at 10,000 cells and infected with two-fold dilution series of stock virus for 60 hours. Cellular expression of luciferase reporter indicating viral infection was determined using Bright-Glo™ Luciferase Assay System (Promega). For neutralization assays, virus stock was diluted to 3-5 × 10^5^ luminescence units.

Pseudovirus neutralization assays were performed on HEK-ACE2 cells seeded at 10,000 cells/well in 40 μL of D10. To determine IC50, blocker dose series were prepared at 3× concentration in D10 media. In 96-well format, 50 μL of 3× blocker and 50 μL of virus were mixed in each well, and the virus and blocker solution was incubated for 1 hour at 37°C. After pre-incubation, 80 μL of the virus and blocker inoculum were transferred to HEK-ACE2 cells. Infection was carried out for 60 hours at 37 °C, at which point the intracellular luciferase signal was measured using the Bright-Glo™ Luciferase Assay (Promega). Neutralization was determined by normalizing the luminescent signal to the average value of the no blocker control. IC50 average values and standard deviations were calculated using 4-8 technical replicates (repeated experiments run at the same time) from 2-4 biological replicates (using different virus stocks and different ACE2 variant preparations).

#### SARS-CoV-2 neutralization assay at biosafety level 3

VeroE6 cells were plated in a 96-well plate at 1.2 × 10^4^ cells per well and incubated overnight. At biosafety level 3, blocking or control (anti-GFP antibody) proteins and the Canadian clinical isolate of SARS-CoV-2 (V-2/Canada/VIDO-01/2027 Feb 2020) were mixed in fresh media supplemented with 3% fetal bovine serum (Gibco) and preincubated for 1 hour at 37°C. The cells were washed once with PBS and infected at the MOI of 0.1 with the proteins for 1 hour at 37 °C and 5% CO2. Next, the mix was removed, and the cells were washed twice with PBS. Complete culture medium was added to each well, and cells were incubated at 37 °C and 5% CO2 for 24 hours followed by cell lysates collection for viral quantitation by qPCR. Mock cells were incubated with the culture supernatant from uninfected VeroE6.

#### SARS-CoV-2 quantitative reverse-transcription PCR assay

For RNA analysis, total RNA was extracted using NucleoSpin RNA kit (Macherey-Nagel) following the manufacturer’s protocol. Total RNA was reverse transcribed using 0.5–1 µg of total RNA and ImProm-II Reverse Transcriptase (Promega) according to the manufacturer’s protocol. qRT-PCR was performed with PerfeCTa SYBR Green SuperMix (Quanta BioSciences) in the CFX96 Touch Real-Time PCR Detection System (Bio-Rad). The cycling conditions were 45 cycles of 94 °C for 30 s, 55 °C for 60 s, and 68 °C for 20 s. Gene expression (fold change) was calculated using the 2^(−ΔΔCT)^ method using human β-actin mRNA transcript as the internal control. The following forward and reverse primer pairs were used for PCR: β-actin 5’-TGGATCAGCAAGCAGGAGTATG-3’ and 5’-GCATTTGCGGTGGACGAT-3’, SARS-CoV-2 spike 5’-CAATGGTTTAACAGGCACAGG-3’ and 5’-CTCAAGTGTCTGTGGATCACG -3’ (2).

#### Additional SARS-CoV-2 neutralization assays at biosafety level 3

Results in Figure S13 were generated from live SARS-CoV-2 neutralization assays performed as previously described (15) at the University of California, San Francisco. Of note, the clinical strain used in the assay was SARS-CoV-2 virus clinical isolate 2019-nCoV/USA-WA1/2020 (BEI resources) and the infection duration was 16 hours instead of 24 hours.

#### Cytotoxicity assay

The CellTiter-Glo Luminescent Cell Viability Assay (Promega) was used for quantitation of ATP in cultured cells. Cells lysates were assayed after mixing 100 µl of complete media with 100 µl of reconstituted CellTiter-Glo Reagent (buffer plus substrate) following the manufacturer’s instructions. Samples were mixed by shaking the plates after which luminescence was recorded with a GloMax Explorer Model GM3510 (Promega) 10 min after adding the reagent.

