## Supplemental Information for "Engineered ACE2 receptor traps potently neutralize SARS-CoV-2"

### Supplemental computational methods: command lines and input files

*Preparation of ACE2-spike structure for modeling.*

Command line for relax:

```
~/Rosetta/source/bin/relax.macosclangrelease -in:file:s 6lzg.pdb -database
~/Rosetta/database -relax:constrain_relax_to_start_coords -
relax:coord_constrain_sidechains -out:suffix _relaxed -beta_nov16 -
corrections::beta_nov16
```

Command line for minimize:

```
~/Rosetta/source/bin/minimize.macosclangrelease -in:file:s
6lzg_relaxed_0001.pdb -database ~/Rosetta/database -out:suffix _min -
beta_nov16 -corrections::beta_nov16
```

*Identification of ACE2 residues that contribute to binding in the ACE2-RBD interface and are chosen for design.*

Command line for determining the energy of each pairwise interaction across the ACE2-Spike interface:

```
~/Rosetta/source/bin/interface_energy.macosclangrelease -in:file:s
6lzg_rm.pdb -face1 face1_ -face2 face2_ -beta_nov16 -corrections::beta_nov16
```

*Computational saturation mutagenesis at selected ACE2 interface residue positions.*

Example command line for running saturation mutagenesis protocol using RosettaScript XML (1):

```
~/Rosetta/Rosetta/main/source/bin/rosetta_scripts.linuxgccrelease -
parser:protocol H34A.xml -in:file:s ../6lzg_rm.pdb @../flags.txt -database
/home/anum/Rosetta/Rosetta/main/database -out:suffix _H34A
```

Example XML:

```
<ROSETTASCRIPTS>

    <SCOREFXNS>
      <ScoreFunction
            name="beta"
            weights="beta_nov16"/>
    </SCOREFXNS>

    <TASKOPERATIONS>
    </TASKOPERATIONS>

    <MOVERS>
      <InterfaceAnalyzerMover
            name="int_ddG"
            scorefxn="beta"
            fixedchains="A_B"/>
```

```
    <MutateResidue
          name="mutate_residue_1"
          target="34A"
          new_res="ALA"/>
    <MinMover
          name="minimize"
          scorefxn="beta"
          chi="1"
          bb="1"
          tolerance="0.005"/>
    <RepackMinimize
          name="repack_interface"
          scorefxn_repack="beta"
          scorefxn_minimize="beta"
          repack_partner1="1"
          repack_partner2="1"
          design_partner1="0"
          design_partner2="0"
          interface_cutoff_distance="6.0"
          repack_non_ala="1"
          minimize_bb="1"
          minimize_rb="1"
          minimize_sc="1"
          optimize_fold_tree="1"/>
  </MOVERS>

  <PROTOCOLS>
    <Add mover_name="mutate_residue_1"/>
    <Add mover_name="repack_interface"/>
    <Add mover_name="minimize"/>
    <Add mover_name="int_ddG"/>
  </PROTOCOLS>

</ROSETTASCRIPTS>
```

Example flags file:

```
-packing
    -ex1
    -ex1aro
    -extrachi_cutoff 0
    -ex2
-nstruct 5
-overwrite
-mute core.util.prof
-mute core.io.database
-corrections::beta_nov16
```

*Redesign of ACE2 interface residues incorporating H34V or H34I mutations.*

Example command line for running Coupled Moves using RosettaScript XML (1, 2):

```
~/Rosetta/Rosetta/main/source/bin/rosetta_scripts.linuxgccrelease -
parser:protocol H34V.xml @H34_flags.txt -database
/home/anum/Rosetta/Rosetta/main/database -out:suffix _CM
```

Example XML:

```
<ROSETTASCRIPTS>

    <SCOREFXNS>
      <ScoreFunction
            name="beta"
            weights="beta_nov16"/>
    </SCOREFXNS>

    <TASKOPERATIONS>
      <ReadResfile
            name="resfile"
            filename="H34.res"/>
    </TASKOPERATIONS>

    <MOVERS>
      <InterfaceAnalyzerMover
            name="int_ddG"
            scorefxn="beta"
            fixedchains="A_B"/>
      <MutateResidue
            name="mutate_residue_1"
            target="34A"
            new_res="VAL"/>
      <MinMover
            name="minimize"
            scorefxn="beta"
            chi="1"
            bb="1"
            tolerance="0.005"/>
      <RepackMinimize
            name="repack_interface"
            scorefxn_repack="beta"
            scorefxn_minimize="beta"
            repack_partner1="1"
            repack_partner2="1"
            design_partner1="0"
            design_partner2="0"
            interface_cutoff_distance="6.0"
            repack_non_ala="1"
            minimize_bb="1"
            minimize_rb="1"
            minimize_sc="1"
            optimize_fold_tree="1"/>
      <CoupledMovesProtocol
            name="coupled_moves"
            task_operations="resfile"/>
    </MOVERS>

    <PROTOCOLS>
```

```
        <Add mover_name="mutate_residue_1"/>
        <Add mover_name="repack_interface"/>
        <Add mover_name="coupled_moves"/>
        <Add mover_name="repack_interface"/>
        <Add mover_name="minimize"/>
        <Add mover_name="int_ddG"/>
    </PROTOCOLS>

</ROSETTASCRIPTS>
```

Example flags file:

```
-in
    -file
        -s 6lzg_rm.pdb

-packing
    -ex1
    -ex1aro
    -extrachi_cutoff 0
    -ex2
-number_ligands 0
-coupled_moves
    -initial_repack false
    -ligand_mode false
    -ligand_weight 0.0
-resfile H34.res
-nstruct 20
-min_pack true
-beta_nov16
-overwrite
-mute core.util.prof
-mute core.io.database
```

Example resfile:

```
NATRO
START

29 A NATAA
30 - 31 A ALLAAxc
32 - 34 A NATAA
35 A ALLAAxc
36 - 37 A NATAA
38 A ALLAAxc
39 A NATAA
416 - 418 B NATAA
452 - 456 B NATAA
492 - 494 B NATAA
```

Supplemental tables

**Table S1. Computational alanine scanning results using established protocols** (3, 4)**.** Columns contain: pdb#: PDB residue number; chain: PDB chain ID (chain A is ACE2, chain B is the spike RBD); int_id: equal to 1 if at least one atom in the residue is within 4 Å of an atom on the other chain, and 0 otherwise; aa: amino acid type; DDG(complex): predicted change in binding energy upon alanine mutation; DG(partner): predicted change in stability of the mutated complex partner upon alanine mutation; DMS beneficial mutations: for ACE2, beneficial point mutations predicted in Procko (5).

| pdb# | chain | int_id | aa | DDG(complex) | DG(partner) | DMS beneficial mutations |
|---|---|---|---|---|---|---|
| 417 | B | 1 | LYS | 0.21 | 0.78 | |
| 449 | B | 1 | TYR | 1.44 | 1.25 | |
| 453 | B | 1 | TYR | 1.37 | 3.11 | |
| 455 | B | 1 | LEU | 1 | 1.6 | |
| 456 | B | 1 | PHE | 1.87 | 1.61 | |
| 486 | B | 1 | PHE | 2.31 | -0.6 | |
| 487 | B | 1 | ASN | 1.71 | 0.64 | |
| 489 | B | 1 | TYR | 1.86 | 1.56 | |
| 493 | B | 1 | GLN | 1.63 | -0.29 | |
| 494 | B | 0 | SER | -0.02 | 0.21 | |
| 498 | B | 1 | GLN | 1.31 | 1.4 | |
| 500 | B | 1 | THR | -0.03 | 0.3 | |
| 501 | B | 1 | ASN | 0.22 | 1.61 | |
| 503 | B | 0 | VAL | 0.03 | -0.1 | |
| 505 | B | 1 | TYR | 2.09 | 0.57 | |
| 19 | A | 1 | SER | 0.71 | -0.42 | V, W, Y, F, P |
| 24 | A | 1 | GLN | 0.59 | 1.01 | T |
| 27 | A | 1 | THR | 0.73 | -0.06 | M, L, A, D, K H, W, Y, F, C |
| 28 | A | 1 | PHE | 0.17 | 3.88 | none |
| 30 | A | 1 | ASP | 0.14 | -0.98 | I, V, E |
| 31 | A | 1 | LYS | 0.61 | -0.47 | W, Y |
| 34 | A | 1 | HIS | 1.86 | -0.59 | V, A, S, P |
| 35 | A | 1 | GLU | 0.66 | -0.24 | M, V, D, C |
| 38 | A | 1 | ASP | 0.35 | -0.67 | none |
| 41 | A | 1 | TYR | 2.39 | 3.13 | R |
| 42 | A | 1 | GLN | 2.71 | -0.59 | M, L, I, V, K, R, H, C |
| 45 | A | 0 | LEU | 0.26 | 0.93 | none |
| 79 | A | 1 | LEU | 0.6 | 0.93 | M, I, V, T, R, W, Y, F, P |
| 82 | A | 1 | MET | 0.5 | 0.4 | R, G, C |
| 83 | A | 1 | TYR | 2.21 | 3.79 | none |
| 351 | A | 0 | LEU | 0.02 | 3.19 | F |
| 353 | A | 1 | LYS | 1.23 | 1.01 | none |
| 355 | A | 1 | ASP | 3.23 | 2.22 | none |

**Table S2. Computational saturation mutagenesis at ACE2 positions 34, 42 and 353 scored using Rosetta total score and interface energies.** Columns contain the following information: Mutation: single-letter amino acid identifier to which original sidechain was mutated: Total energy: Rosetta score for the complex (in REU); Interface: the lowest calculated interface energy of five trials (in REU); delta: the difference between the interface energy of the wild-type complex and the interface energy of the point mutant (in REU). Interface energies lower than those determined for the WT complex are highlighted in yellow (see "Computational saturation mutagenesis at targeted ACE2 interface residue positions," Methods).

| H34 | | | | Q42 | | | | K353 | | | |
|---|---|---|---|---|---|---|---|---|---|---|---|
| Mutation | Total energy | Interface | delta | Mutation | Total energy | Interface | delta | Mutation | Total energy | Interface | delta |
| A | -2079.41 | -58.1059 | -0.974 | A | -2075.896 | -53.869 | 3.263 | A | -2074.771 | -54.8121 | 2.320 |
| R | -2075.486 | -55.5158 | 1.617 | R | -2079.445 | -55.8825 | 1.250 | R | -2073.901 | -52.8174 | 4.315 |
| N | -2077.512 | -58.2728 | -1.141 | N | -2077.357 | -55.039 | 2.093 | N | -2074.215 | -57.2794 | -0.147 |
| D | -2072.993 | -55.7567 | 1.376 | D | -2075.277 | -54.0632 | 3.069 | D | -2067.808 | -54.8079 | 2.324 |
| Q | -2077.21 | -58.9142 | -1.782 | Q | | | | Q | -2073.781 | -54.8442 | 2.288 |
| E | -2076.387 | -56.5619 | 0.570 | E | -2078.102 | -53.8338 | 3.299 | E | -2068.393 | -50.321 | 6.811 |
| G | -2075.524 | -56.1769 | 0.955 | G | -2074.099 | -53.8018 | 3.331 | G | -2070.87 | -51.5659 | 5.566 |
| H | | | | H | -2077.636 | -54.026 | 3.106 | H | -2071.823 | -56.1115 | 1.021 |
| I | -2068.827 | -53.3366 | 3.796 | I | -2078.778 | -54.1272 | 3.005 | I | -2064.506 | -52.8856 | 4.247 |
| L | -2072.173 | -57.2587 | -0.126 | L | -2079.427 | -55.135 | 1.997 | L | -2075.772 | -54.933 | 2.199 |
| K | -2075.855 | -54.9393 | 2.193 | K | -2079.221 | -56.2299 | 0.902 | K | | | |
| M | -2076.286 | -58.3653 | -1.233 | M | -2078.496 | -53.8281 | 3.304 | M | -2073.167 | -57.2724 | -0.140 |
| F | -2077.452 | -58.6328 | -1.501 | F | -2076.563 | -53.973 | 3.159 | F | -2075.907 | -54.8568 | 2.276 |
| P | -2062.583 | -56.1616 | 0.971 | P | -2065.709 | -52.2274 | 4.905 | P | -1984.256 | -53.6565 | 3.476 |
| S | -2076.239 | -55.6296 | 1.503 | S | -2076.745 | -53.8748 | 3.258 | S | -2069.483 | -56.5399 | 0.592 |
| T | -2075.284 | -58.5258 | -1.394 | T | -2076.591 | -54.0055 | 3.127 | T | -2067.071 | -55.8814 | 1.251 |
| W | -2076.567 | -58.0243 | -0.892 | W | -2077.569 | -54.8777 | 2.255 | W | -2068.537 | -55.7901 | 1.342 |
| Y | -2077.384 | -58.5473 | -1.415 | Y | -2075.443 | -54.2037 | 2.929 | Y | -2073.076 | -56.874 | 0.258 |
| V | -2078.676 | -59.8262 | -2.694 | V | -2077.3 | -53.9666 | 3.166 | V | -2066.138 | -55.3603 | 1.772 |
| WT | -2048.31 | -57.1323 | 0.000 | WT | -2048.31 | -57.1323 | 0.000 | WT | -2048.31 | -57.1323 | 0.000 |

**Table S3. Apparent binding affinities of ACE2(614) variants measured on the surface of yeast with monomeric SARS-CoV-2 spike RBD.** $K_{D,app}$ values reported as the average from the fit to all data in duplicate experiments, as shown in Figure 3D-E, with the errors of the fit. Aga2p-GFP constructs were used in all yeast surface display experiments (6). These are listed alongside the ACE2(614)-Fc and ACE2(740)-Fc constructs that include the same affinity-enhancing mutations for convenience.

| **Mutations** | **Origin** | **$K_{D,app}$ (nM) (on yeast)** | **Aga2p-GFP construct** | **ACE2(614)-Fc construct** | **ACE2(740)-Fc construct** |
|---|---|---|---|---|---|
| - | WT | 20.4 ± 1.8 | Y208 | CVD013 | CVD208 |
| H34V | Computational design | 9.29 ± 0.78 | Y295 | CVD014, CVD127* | CVD295 |
| N90Q | (5) | 4.54 ± 0.55 | Y117 | CVD117 | |
| K31F, H34I, E35Q | Computational design | 1.71 ± 0.02 | Y293 | CVD019 | CVD293 |
| H34V, N90Q | Computational design | 5.47 ± 0.68 | Y292 | CVD118*, CVD278† | CVD292 |
| A25V, T27Y, H34A, F40D | DMS-guided design | 0.40 ± 0.03 | Y310 | | CVD310† |
| K31Y, W69V, L79T, L91P | DMS-guided design | 0.64 ± 0.05 | Y311 | | CVD311† |
| T27Y, H34A, N90Q | DMS-guided design | 0.84 ± 0.09 | Y312 | | CVD312† |
| S19P, Q42L, L79T, N90Q | DMS-guided design | 0.94 ± 0.09 | Y355 | | CVD355† |
| K31F, N33D, H34S, E35Q | Yeast display, round 3.1 | 0.52 ± 0.04 | Y313 | | CVD313† |
| K31F, N33D, H34A, E35Q, N49D, N51S, N53S, E57G, N64D | Yeast display, round 3.2 | 0.45 ± 0.18 | Y354 | | CVD354† |
| T27A, K31F, N33D, H34S, E35Q, N61D, K68R, L79P | Yeast display, round 4 | 0.19 ± 0.02 | Y353 | | CVD353† |
| S19P, N33S, H34V, F40L, N49D, L100P | Yeast display, round 4 | 0.61 ± 0.04 | Y375 | | CVD375† |
| Q18R, K31F, N33D, H34S, E35Q, W69R, Q76R | Yeast display, round 5 | 0.12 ± 0.01 | Y373 | | CVD373† |

* includes H374N/H378N inactivation mutations
† includes H345L inactivation mutation

**Table S4. Half-maximal inhibitory concentrations (IC50s) of ACE2 variants.** IC50 values were calculated from titration experiments in pseudotyped lentiviral SARS-CoV-2 neutralization assays and authentic SARS-CoV-2 neutralization assays in a biosafety level 3 facility with VeroE6 cells. IC50 values reported as means. Errors reported for pseudovirus experiments are standard deviations between 4 and 12 technical replicates. Errors reported for authentic SARS-CoV-2 experiments represent the error of the fit for biological replicates.

| ID | Mutations | Scaffold | IC50 (μg/ml), pseudovirus | IC50 (μg/ml), VeroE6 |
|---|---|---|---|---|
| CVD013 | - | ACE2(614)-Fc | 0.43 ± 0.39 | |
| CVD208 | - | ACE2(740)-Fc | 0.71 ± 0.51 | |
| CVD208m | - | ACE2(740) | 2.19 ± 1.15 | |
| CVD014 | H34V | ACE2(614)-Fc | 0.35 ± 0.19 | |
| CVD019 | K31F, H34I, E35Q | ACE2(614)-Fc | 0.31 ± 0.16 | |
| CVD118 | H34V, N90Q, H374N†, H378N† | ACE2(614)-Fc | | < 0.5 |
| CVD293 | K31F, H34I, E35Q | ACE2(740)-Fc | 0.036 ± 0.01 | 0.136 ± 0.08 |
| CVD310 | A25V, T27Y, H34A, F40D, H345L† | ACE2(740)-Fc | 0.058 ± 0.03 | 0.089 ± 0.01 |
| CVD311 | K31Y, W69V, L79T, L91P, H345L† | ACE2(740)-Fc | 0.055 ± 0.03 | |
| CVD313 | K31F, N33D, H34S,E35Q, H345L† | ACE2(740)-Fc | 0.028 ± 0.02 | 0.073 ± 0.02 |
| CVD353 | T27A, K31F, N33D, H34S,E35Q, N61D, K68R, L79P, H345L† | ACE2(740)-Fc | 0.69 ± 0.38 | |

† mutation inactivating the ACE2 enzymatic function

Supplemental figures

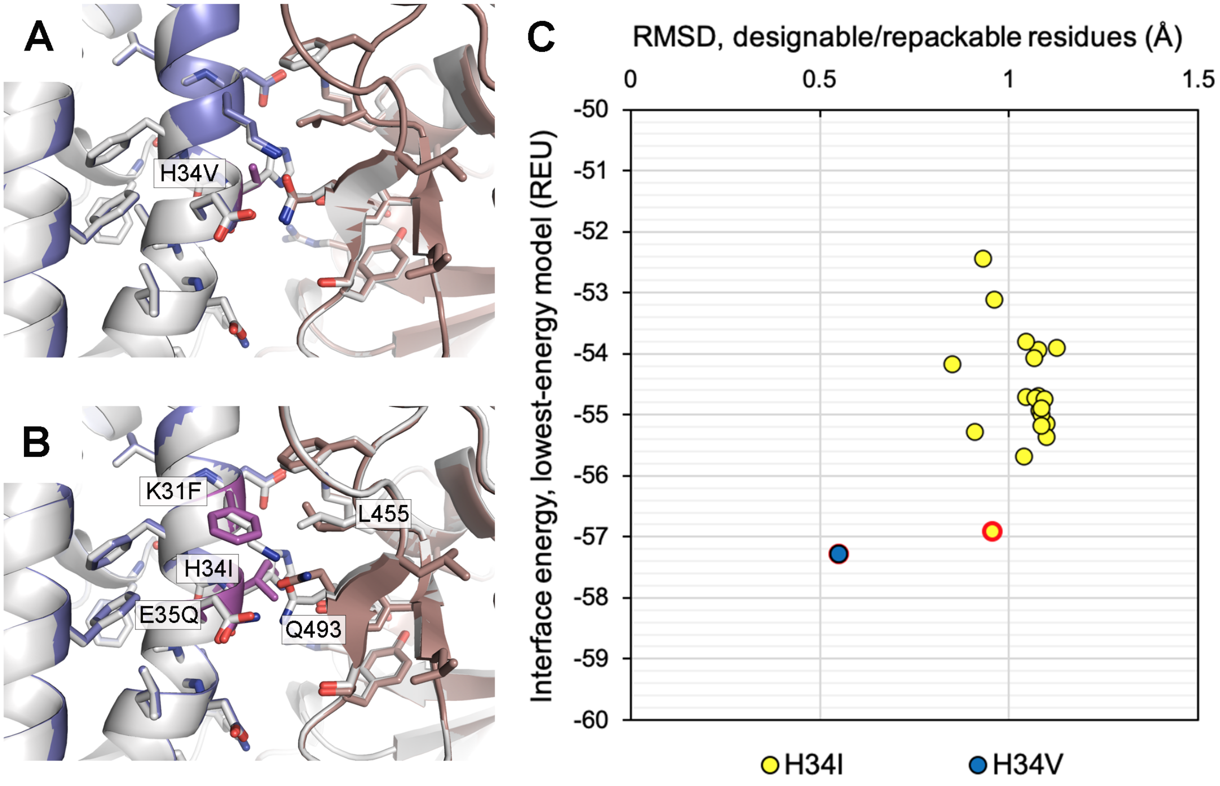

**Figure S1. Comparison between models of redesigned ACE2-SARS-CoV2 spike RBD interfaces in ACE2 H34V and H34I backgrounds.** Lowest-energy models for redesigned ACE2 in the **(A)** H34V and **(B)** H34I mutation backgrounds. The wild-type ACE2-RBD interface is shown in white. Redesigned ACE2 is shown in blue. Repacked RBD is shown in dark salmon. Mutated residues in the redesigned ACE2 are shown in magenta. Sidechains with any atoms within 6 Å of position 34 are shown as sticks. Mutated ACE2 residues and RBD residues that adopted different rotameric conformations are labeled. **(C)** Root-mean-square deviation (RMSD) from WT ACE2 for all backbone and sidechain heavy atoms belonging to mutable and redesignable residues vs. summed pairwise interface energies for lowest energy solutions for all trials of ACE2-spike RBD interface design in H34V and H34I backgrounds. All the solutions for the H34V models have very similar energies and RMSD, but solutions in the H34I background are diverse. Red-circled points are the lowest-energy solutions depicted in **(A)** and **(B)**. Models based on PDB 6LZG (7).

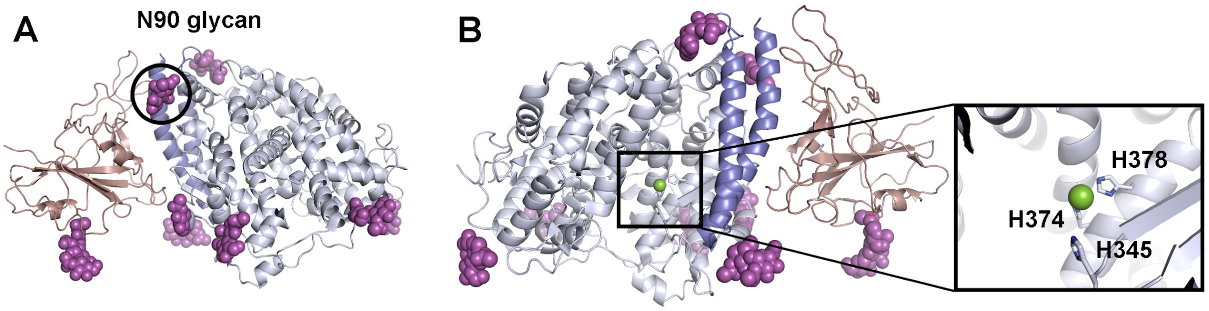

**Figure S2. Additional ACE2 mutations for enzyme inactivation and improved affinity to the spike RBD. (A)** Procko (5) showed that mutants in which the N90 glycan (circled) adjacent to the ACE2-RBD interface was knocked out were enriched in DMS selection experiments. **(B)** The active site of ACE2 (square region) binds a $Zn^{2+}$ ion, which is coordinated by H378 and H374 (8). H345, also shown, is important for substrate binding in catalysis (9). The spike RBD is shown in dark salmon, ACE2 is shown in light blue with N-terminal helices (residues 18-90) shown in dark blue, glycans are shown in magenta, and the $Zn^{2+}$ ion is shown in green. Structures are from PDB 6M17 (10).

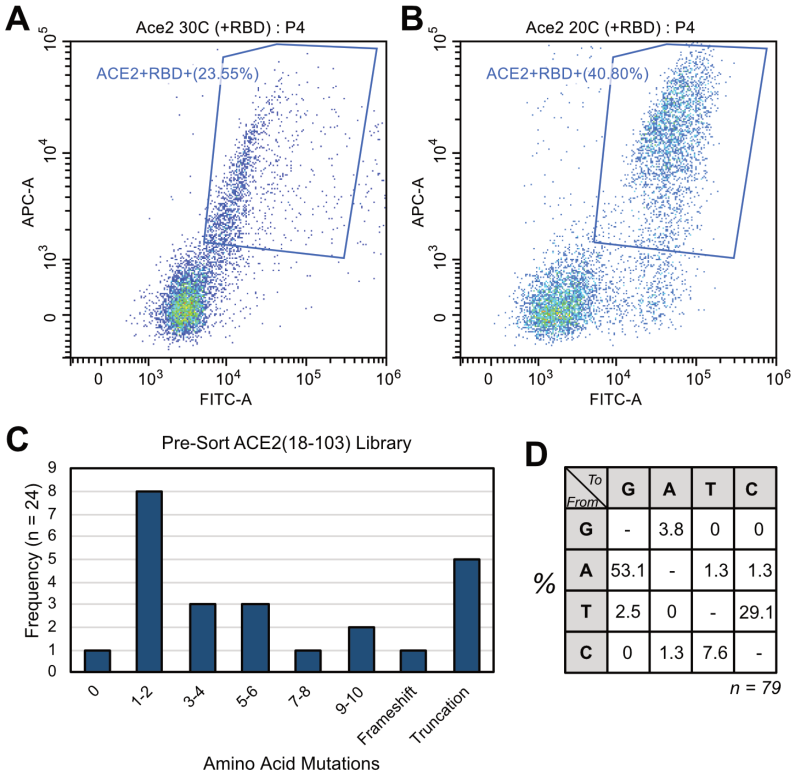

**Figure S3. Yeast surface display of ACE2(614) and library analysis.** WT ACE2(614) was expressed as an Aga2p fusion at **(A)** 30 °C and **(B)** 20 °C and bound to 100 nM biotinylated Spike-RBD-Fc followed by streptavidin Alexa Fluor 647. In **(A)** and **(B)**, the FITC-A axis represents protein expression, and the APC-A axis represents binding to the RBD. **(C)** Analysis of 24 clones of ACE2(614) mutagenized at amino acids 18-103 by error-prone PCR showed a broad distribution of mutations per clone. **(D)** 79 total DNA mutations from these 24 clones had mutational bias expected from dNTP analog epPCR.

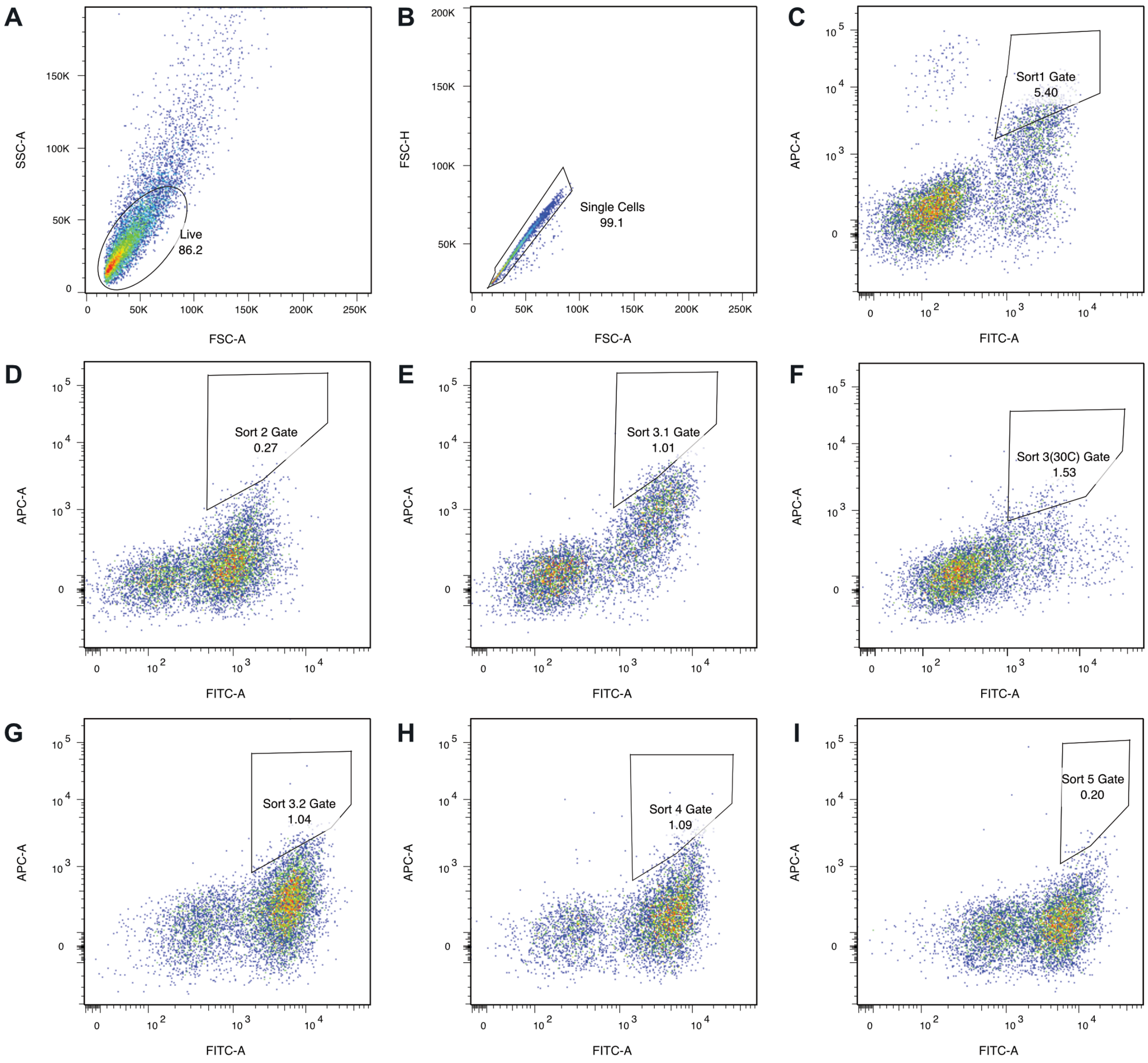

**Figure S4. ACE2(614) affinity maturation sort gates.** Each sample was gated on **(A)** live cells and **(B)** singlets. Approximate sort gates are shown for **(C)** Sort 1 (50 nM RBD monomer), **(D)** Sort 2 (5 nM RBD monomer), **(E)** Sort 3.1 (0.5 nM RBD monomer), **(F)** Sort 3 (0.5 nM RBD monomer, 30 °C expression), **(G)** Sort 3.2 (0.2 nM RBD monomer), **(H)** Sort 4 (8 hour dissociation with 20 nM soluble H34V-ACE2(614)-Fc competitor), and **(I)** Sort 5 (12 hour dissociation with 20 nM soluble H34V-ACE2(614)-Fc competitor).

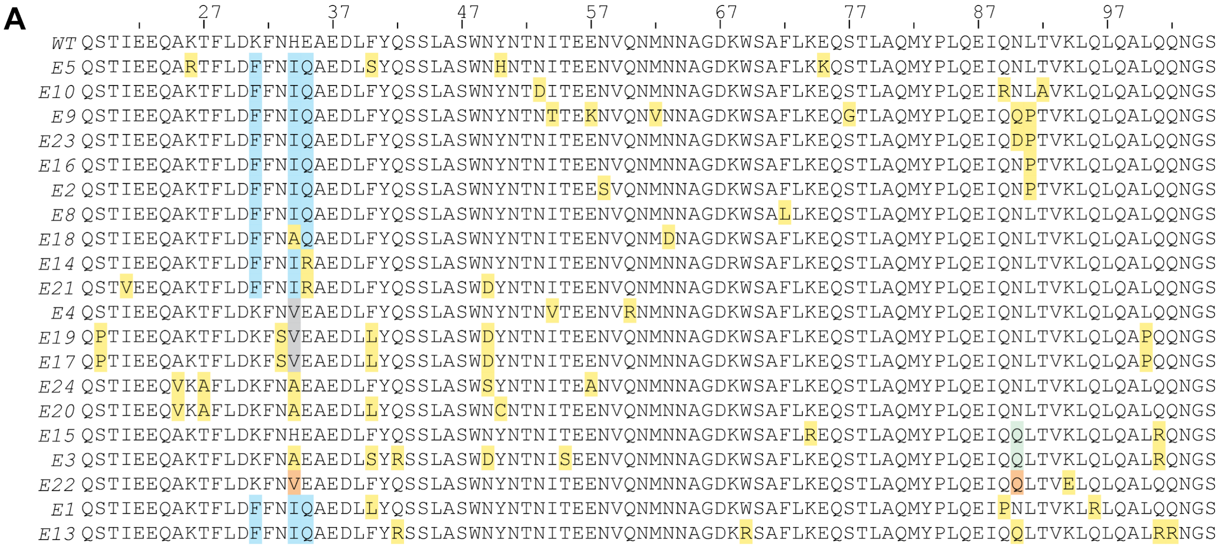

**Figure S5. Sorts 4-5 ACE2 sequence alignments. (A)** Sanger sequencing of individual clones from Sort 4 showed no convergence. **(B)** Sort 5 was enriched for Q18R/K31F/N33D/H34S/E35Q/W69R/Q76R ACE2(614). The colors are as follows: yellow, mutations from error-prone PCR; blue, mutation most likely from K31F/H34I/E35Q parent; gray, mutation most likely from H34V parent; green, mutation most likely from N90Q parent; brown, mutation most likely from H34V/N90Q parent.

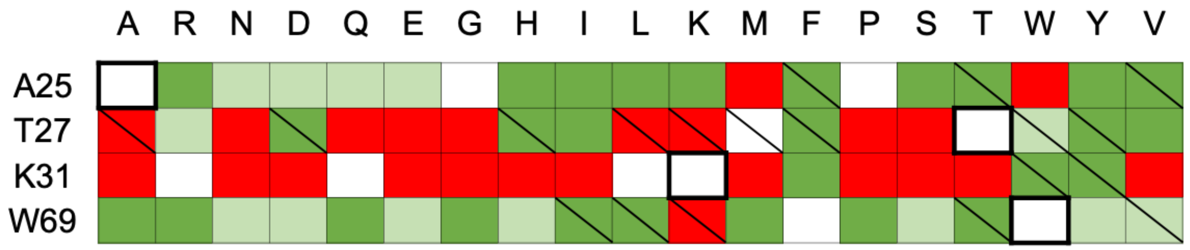

**Figure S6. Computational saturation mutagenesis on additional positions in the ACE2-spike RBD interface.** Interface energies for the whole ACE2-spike RBD interface were calculated on the lowest total energy models from computational saturation mutagenesis at several positions that were not alanine-scanning hotspots. Boxes are colored according to the interface energy difference of the point mutant model with the WT ACE2-RBD: red, DDG > 0.5 REU; white, DDG ≤ 0.5 REU and ≥ -0.6 REU; light green, DDG < -0.6 REU and ≥ -0.9 REU; dark green, DDG < -0.9 REU. Thick lines around boxes indicate the WT amino acid. Boxes with diagonal lines indicate amino acid substitutions identified as beneficial by DMS (5).

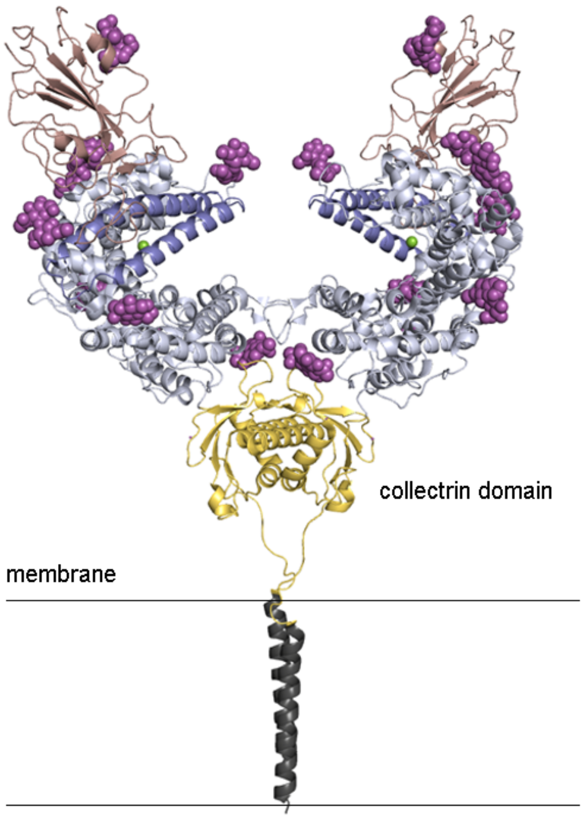

**Figure S7. ACE2 residues 615-740 form a collectrin domain.** In ACE2, the collectrin domain (yellow) connects the transmembrane helices (dark gray) to the soluble extracellular peptidase domain (residues 18-90 in blue, residues 91-614 in light blue). The SARS-CoV-2 spike RBD is shown in dark salmon. The $Zn^{2+}$ ion is shown in green. All glycans are shown in magenta. PDB 6M17 (10).

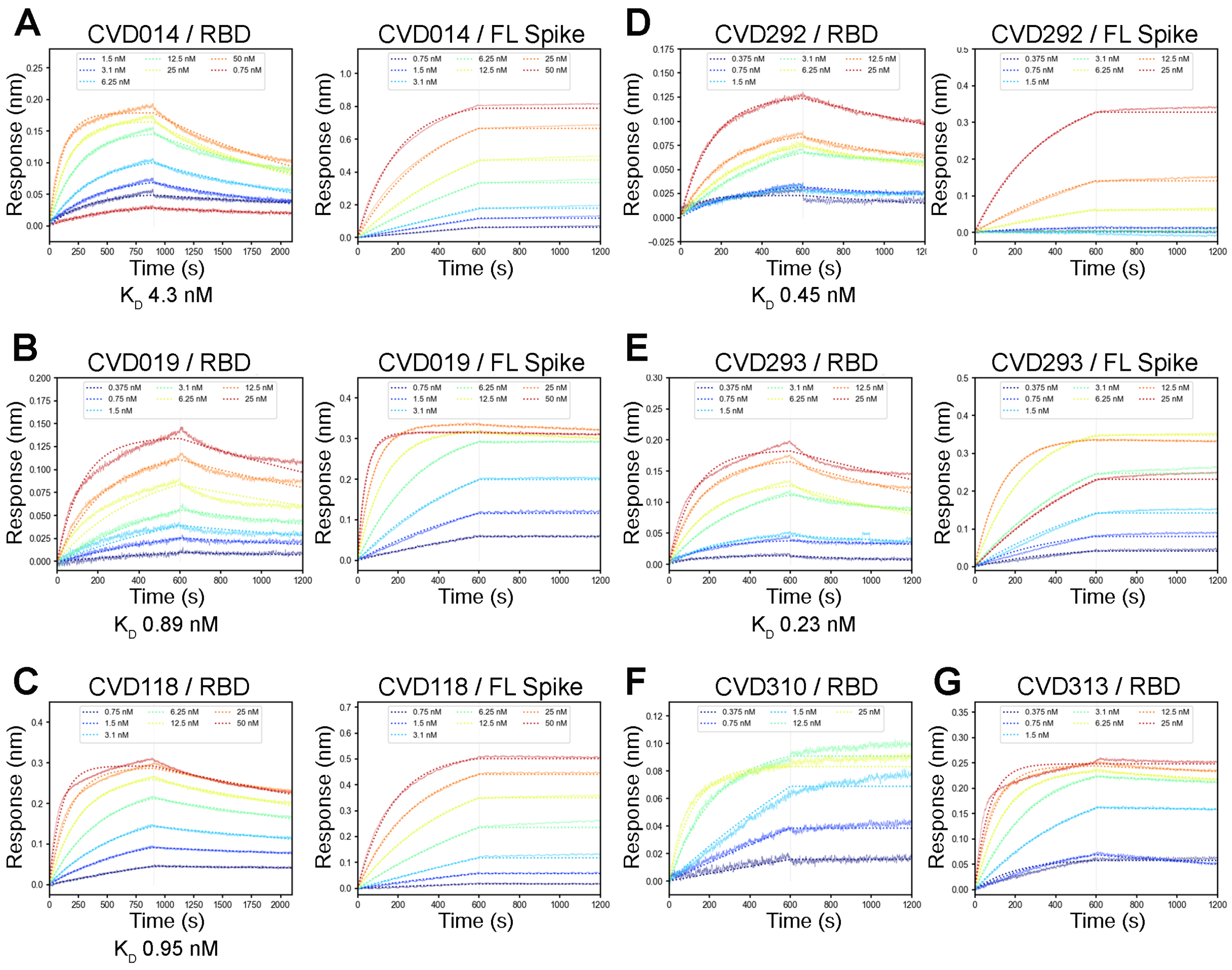

**Figure S8. ACE2-Fc variants with and without collectrin domain bind full-length (FL) spike more tightly than spike RBD.** Representative BLI measurements show that designed **(A-C)** ACE2(614)-Fc variants and **(D-E)** ACE2(740)-Fc variants have a higher binding affinity for FL spike (right) as compared to the monomeric spike RBD (left). $K_D$ for all ACE2 variants binding to RBD are reported. Due to decreased off-rates, $K_D$ for FL spike could not be calculated. **(F-G)** ACE2(740)-Fc variants from DMS-guided design and affinity maturation in yeast bind the spike RBD with decreased off-rates, precluding accurate estimation of binding affinity from BLI experiments.

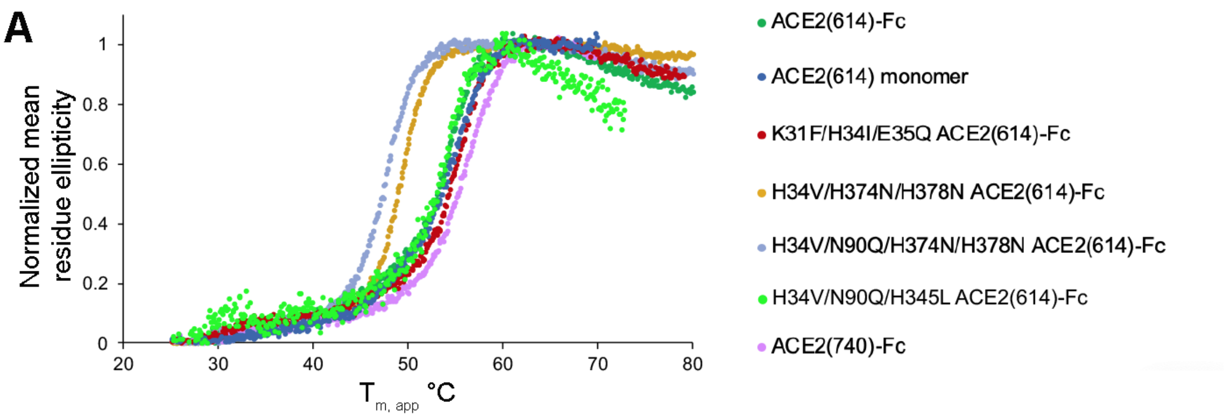

| CVD118 | CVD127 | CVD013 | CVD278 | CVD013-m | CVD019 | CVD208 |
|---|---|---|---|---|---|---|
| ACE2(18-614)-Fc H34V, N90Q, H374N, H378N | ACE2(18-614)-Fc H34V, H374N, H378N | ACE2(18-614)-Fc | ACE2(18-614)-Fc H34V, N90Q, H345L | ACE2(18-614) monomer | ACE2(18-614) K31F, H34I, E35Q | ACE2(18-740)-Fc |
| 47.3 °C | 48.9 °C | 51.8 °C | 52.7 °C | 53.1 °C | 54.1 °C | 55.3 °C |

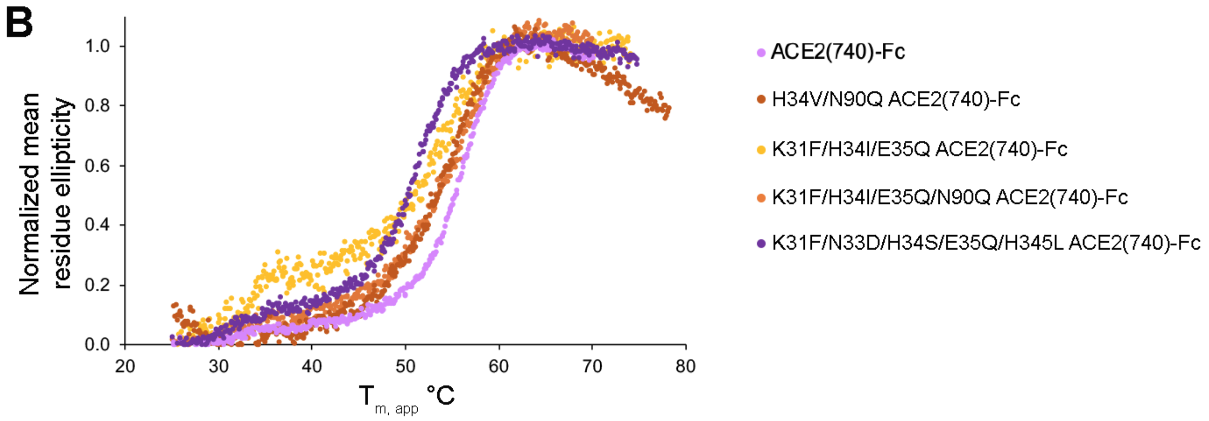

| CVD293 | CVD313 | CVD294 | CVD292 | CVD208 |
|---|---|---|---|---|
| ACE2(18-740)-Fc K31F, H34I, E35Q | ACE2(18-740)-Fc K31F, N33D, H34S, E35Q, H345L | ACE2(18-740)-Fc K31F, H34I, E35Q, N90Q | ACE2(18-740)-Fc H34V, N90Q | ACE2(18-740)-Fc |
| 50.0 °C | 50.2 °C | 52.5 °C | 52.6 °C | 55.3 °C |

**Figure S9. Thermostability of ACE2-Fc constructs measured by circular dichroism spectroscopy (CD). (A)** CD melt curves show that affinity-enhancing mutations do not greatly reduce protein stability, but that mutations to residues coordinating the $Zn^{2+}$ ion (H374N, H378N) do destabilize the ACE2-Fc scaffold. Catalytic inactivation by the H345L mutation is not destabilizing and has similar affinity to the WT ACE2-Fc (Figure S10). Inclusion of the collectrin domain, ACE2 residues 615-740, further enhances stability. **(B)** CD melt curves show slight destabilization of ACE2(740)-Fc variants with computationally-designed (CVD292, 293, 294 variants) and yeast-selected (CVD313) affinity-enhancing mutations. These variants have apparent melting temperatures ($T_{m,app}$) in the 50-53 °C range.

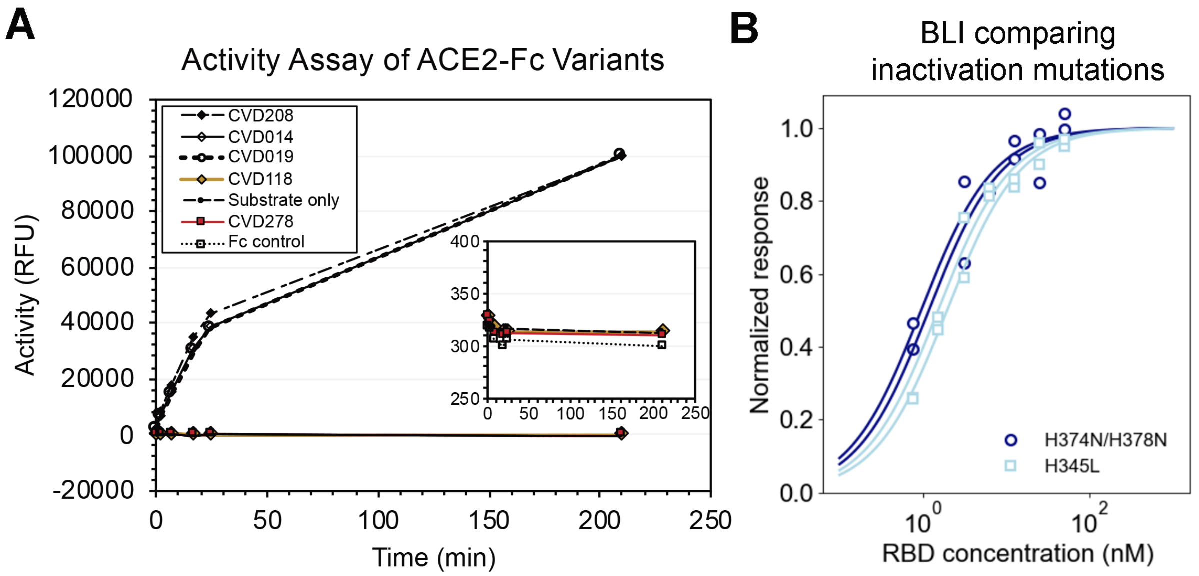

**Figure S10. *In vitro* binding and activity characterization of inactivation mutations in ACE2-Fc. (A)** Enzyme activity was assayed by monitoring the increase in fluorescence resulting from hydrolysis of Mca-APK-DNP. CVD014 and 019 had similar activity to wild-type CVD208 but the H374N/H378N mutations (in CVD118) and the H345L mutation (in CVD278) have no detectable catalytic activity. **(B)** BLI data comparing the effect on binding the spike RBD for the different inactivation mutations, H374N/H378N and H345L. Both ACE2-Fc constructs also contain affinity-enhancing mutations H34V and N90Q.

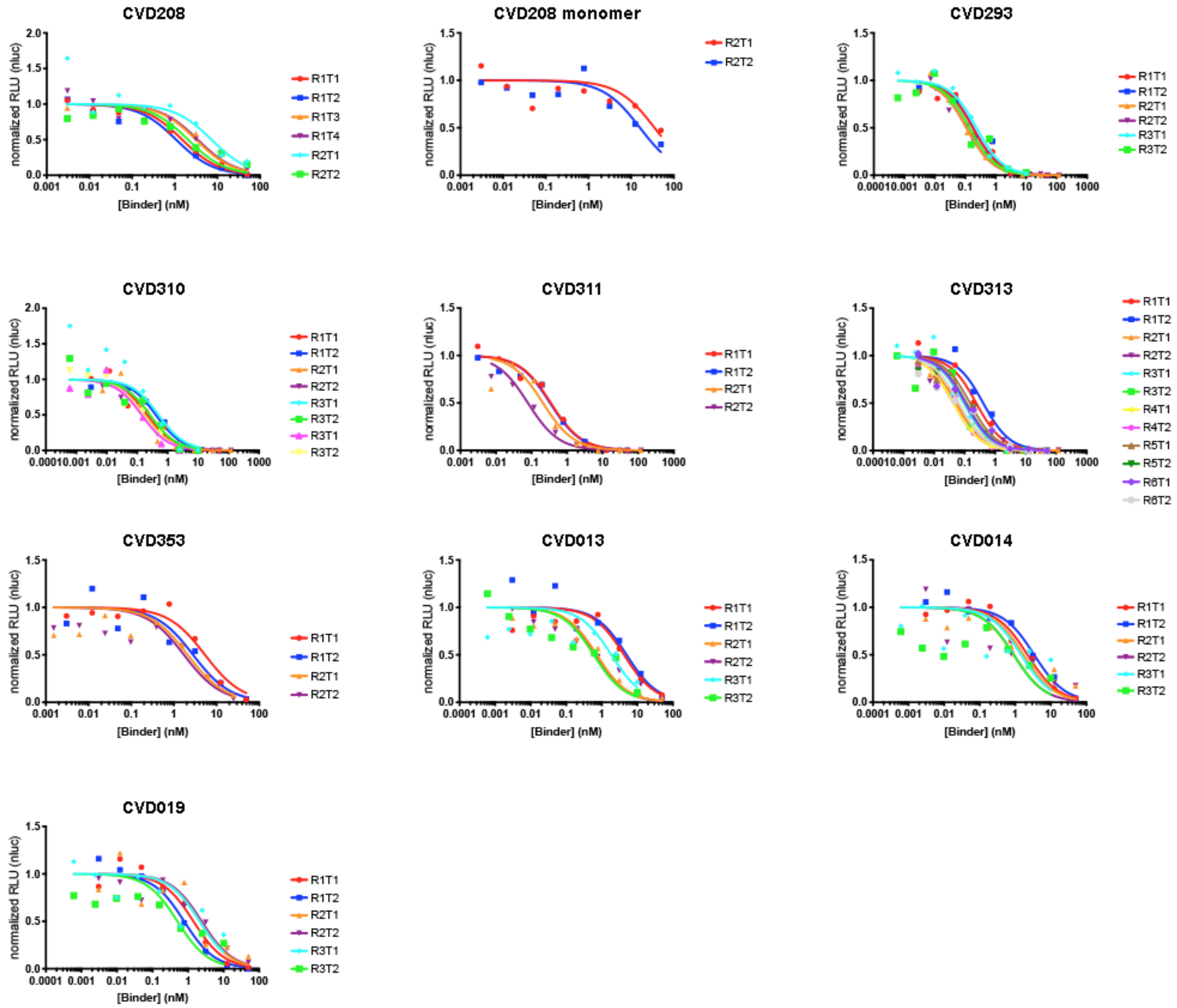

**Figure S11. Pseudotyped SARS-CoV-2 neutralization IC50 curves for WT and engineered ACE2(614)-Fc and ACE2(740)-Fc molecules.** Normalized luminescent response for each technical replicate is shown as a separate line. Labels indicate biological replicate (R) and technical replicate (T) for each experiment (e.g. R1T1, biological replicate 1 and technical replicate 1).

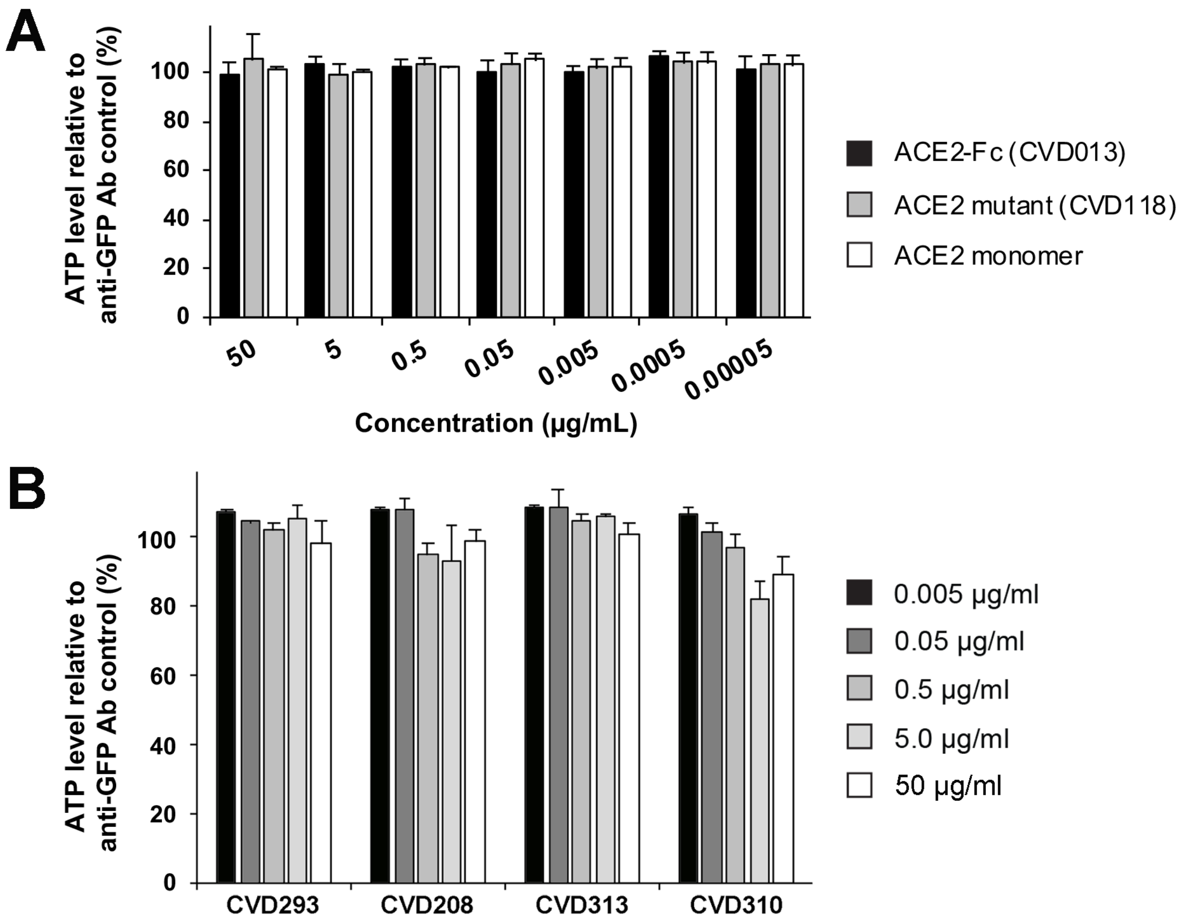

**Figure S12. ACE2 variants are not strongly cytotoxic in uninfected VeroE6 cells after 24 hours of treatment.** ATP release was measured by luminescence signal using the CellTiter-Glo assay (Promega), divided by the signal from the anti-GFP IgG control experiment, and multiplied by 100 for separate experiments. Error bars represent standard error of the mean for biological duplicates.

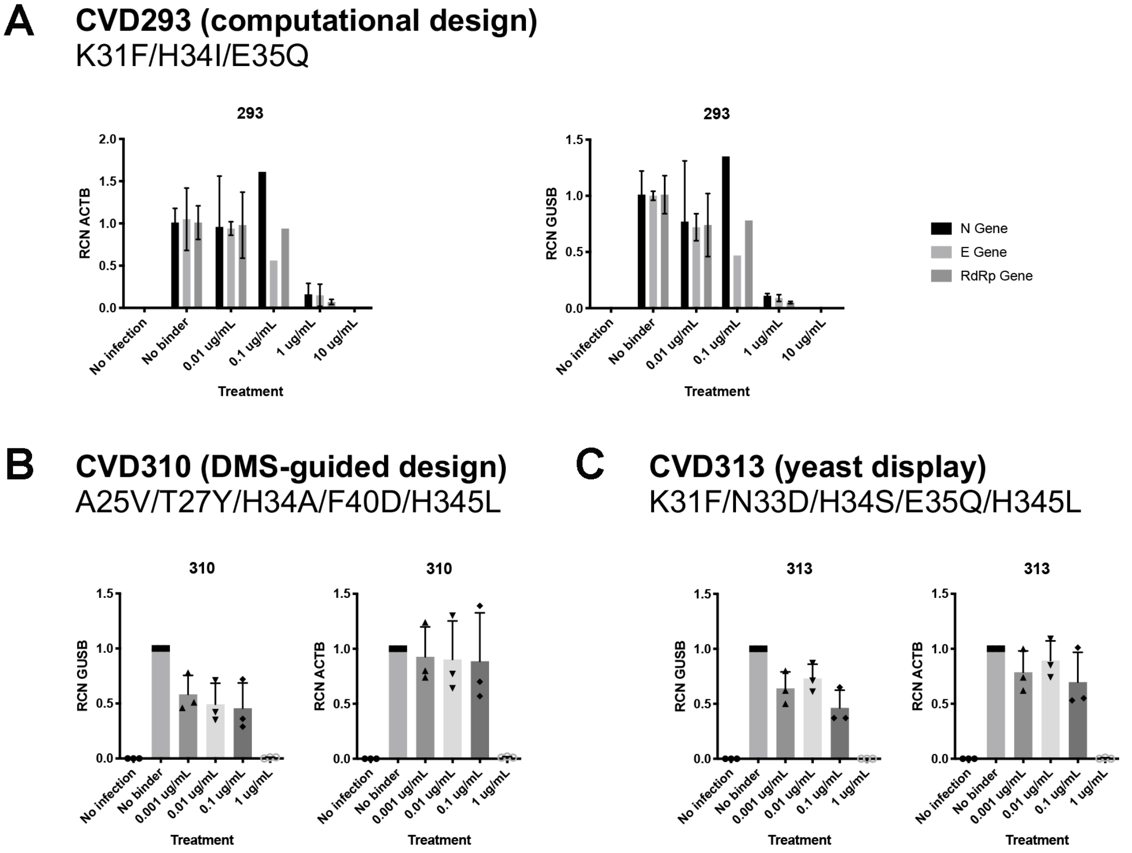

**Figure S13. Computationally designed and affinity-matured ACE2(740)-Fc variants effectively block infection of VeroE6 cells by SARS-CoV-2.** Live virus qPCR assays of three viral genes (N, E, RdRp) were run 16 hours post-infection using host genes BGUS or ACTB as normalization controls. **(A)** K31F/H34I/E35Q ACE2(740)-Fc (variant 293) has an IC50 between 0.1 and 1 µg/ml. Error bars represent technical duplicates. **(B)** A25V/T27Y/H34A/F40D/H345L ACE2(740)-Fc (variant 310) and **(C)** K31F/N33D/H34S/E35Q/H345L ACE2(740)-Fc (variant 313) neutralize completely at 1 µg/ml. Symbols in **(B)** and **(C)** represent the signal for the individual viral genes.

```
A CoV-2 328 RFPNITNLCPFGEVFNATRFASVYAWNRKRISNCVADYSVLYNSASFSTFKCYGVSPTKLNDLCFTNVYA 397
  CoV-1 318 ---NITNLCPFGEVFNATKFPSVYAWERKKISNCVADYSVLYNSTFFSTFKCYGVSATKLNDLCFSNVYA 384
  NL63       ----------------------------------------------------------------------

  CoV-2 398 DSFVIRGDEVRQIAPGQTGKIADYNYKLPDDFTGCVIAWNSNNLDSKVGGN---------YN--YLYR-- 454
  CoV-1 385 DSFVVKGDDVRQIAPGQTGVIADYNYKLPDDFMGCVLAWNTRNIDATSTGN---------YN--YKYR-- 441
  NL63  481 -------------------QHTDINFTATASFGGSCYVCKPHQVNISLNGNTSVCVRTSHFSIRYIYNRV 531

  CoV-2 455 -----------LFRKSNLKPFERD-------ISTEIYQAGSTPCNGVEGFNCYFPLQSYGFQP---TNGV 503
  CoV-1 442 -----------YLRHGKLRPFERD-------ISNVPFSPDGKPCTP-PALNCYWPLNDYGFYT---TTGI 489
  NL63  532 KSGSPGDSSWHIYLKSGTCPFSFSKLNNFQKFKTICFSTVEV------PGSCNFPLEATWHYTSYTIVGA 595

  CoV-2 504 GYQPYRVVVLSFELLHAPATVCGPKKSTNL                                         533
  CoV-1 490 GYQPYRVVVLSFELLNAPATV---------                                         510
  NL63  596 LYVTWS---EGNSITGVPYPVSGI------                                         616
```

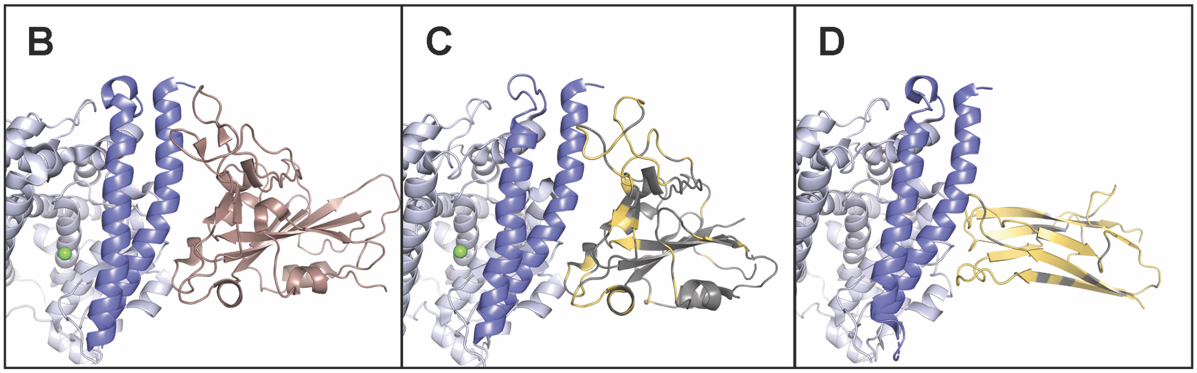

**Figure S14. Sequences and structures of receptor binding domains from SARS-CoV-2, SARS-CoV-1, and HCoV-NL63 spike proteins. (A)** Sequence alignment for RBDs from all three spike proteins. Yellow residues in the SARS-CoV-1 and HCoV-NL63 RBD sequences (bottom rows) are different from the residue at that position in the SARS-CoV-2 RBD (top row). Numbers flanking each row indicate residue positions for each RBD sequence. **(B)** Structure of the ACE2-SARS-CoV-2 RBD interface (PDB 6LZG) (7). **(C)** Structure of the ACE2-SARS-CoV-1 RBD interface (PDB 2AJF) (11). **(D)** Structure of the ACE2-NL63 RBD interface (PDB 3KBH) (12). In **(B-D)**, ACE2 residues 18-90 are colored dark blue, ACE2 residues 91-614 are colored light blue, and the active site $Zn^{2+}$ ion is colored green. In **(B)**, the RBD is colored dark salmon. In **(C)** and **(D)**, the RBD residues are colored by sequence alignment: residues that are white in **(A)** are colored gray in the structures, and residues that are yellow in **(A)** are colored yellow in the structures.

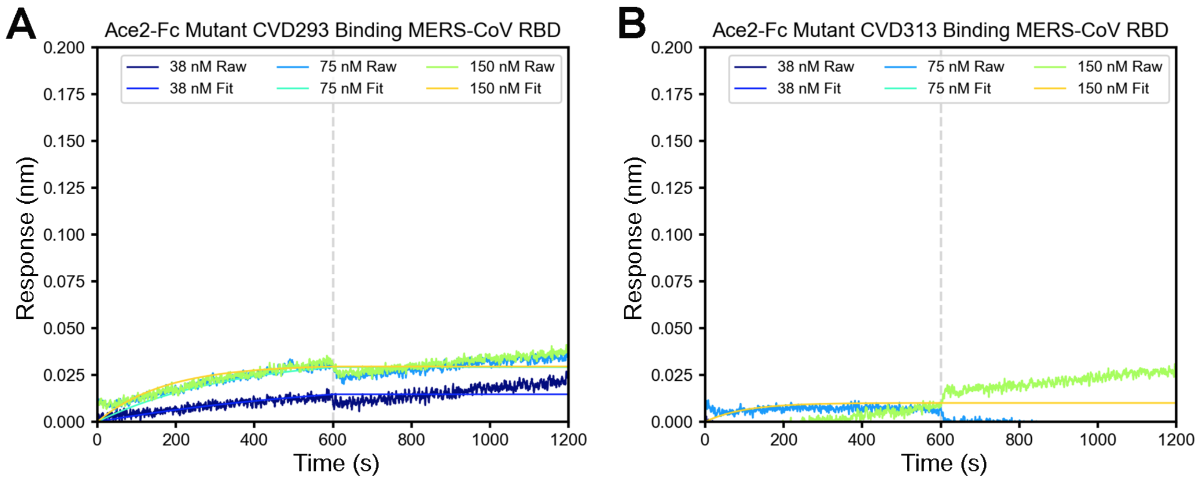

**Figure S15. Computationally designed and affinity-matured ACE2(740)-Fc variants do not bind MERS RBD.** BLI measurements for **(A)** K31F/H34I/E35Q ACE2(740)-Fc binding MERS-CoV RBD and **(B)** K31F/N33D/H34S/E35Q/H345L ACE2(740)-Fc binding the MERS-CoV RBD.
