## Appendix 1 for "Engineered ACE2 receptor traps potently neutralize SARS-CoV-2"

Appendix 1: Sequences of ACE2 mutants in this work

| | |
|---|---|
| 13 | CAATCTACCATCGAAGAGCAGGCCAAAACATTCCTCGACAAGTTTAATCACGAGGCTGAAGACC<br>TTTTCTACCAATCAAGTCTGGCTAGCTGGAATTACAATACAAACATTACAGAGGAGAACGTACA<br>AAACATGAATAACGCAGGGGACAAGTGGAGCGCATTCCTTAAGGAACAAAGTACCCTTGCGCAA<br>ATGTATCCGCTGCAAGAGATTCAAAACCTGACGGTTAAGCTGCAACTTCAGGCCCTCCAACAAA<br>ATGGAAGTTCAGTCTTGTCAGAAGACAAAAGCAAGCGACTGAACACCATCCTTAACACCATGTC<br>AACCATATATTCAACAGGTAAAGTTTGCAATCCGGATAACCCCCAAGAATGTTTGCTTCTTGAA<br>CCCGGTCTCAACGAAATTATGGCCAACAGTCTTGATTACAACGAGCGATTGTGGGCATGGGAAA<br>GTTGGAGGAGTGAGGTAGGCAAACAGTTGAGACCTCTTTATGAAGAGTACGTTGTCCTTAAAAA<br>TGAAATGGCTCGCGCGAATCATTATGAAGACTATGGTGACTACTGGAGGGGGGATTATGAGGTG<br>AACGGGGTGGACGGATACGATTACTCTAGGGGCCAGCTGATAGAGGATGTCGAGCACACCTTTG<br>AGGAGATTAAGCCGTTGTACGAACATTTGCACGCCTATGTCAGGGCTAAGCTCATGAACGCTTA<br>TCCGAGTTATATCTCCCCGATAGGATGCTTGCCTGCTCACTTGTTGGGCGATATGTGGGGACGC<br>TTTTGGACCAACTTGTATTCCCTTACGGTACCGTTCGGCCAGAAACCAAATATCGACGTGACAG<br>ACGCAATGGTGGATCAAGCATGGGATGCGCAACGAATCTTCAAGGAGGCAGAAAAATTTTTCGT<br>TTCAGTTGGACTCCCAAACATGACGCAGGGTTTCTGGGAGAACTCAATGTTGACAGATCCAGGT<br>AATGTGCAGAAAGCGGTTTGCCACCCTACTGCATGGGATCTTGGTAAAGGGGACTTCCGCATAC<br>TCATGTGTACGAAAGTAACTATGGACGACTTTCTTACTGCGCACCACGAGATGGGGCACATACA<br>ATACGATATGGCGTACGCAGCTCAACCTTTCCTTCTGCGGAACGGGGCGAATGAAGGATTTCAC<br>GAGGCAGTGGGTGAGATTATGTCCCTGTCAGCTGCCACTCCGAAACATCTGAAAAGCATCGGCC<br>TGTTGAGCCCAGACTTCCAAGAAGATAATGAGACCGAAATAAACTTCCTTCTGAAGCAAGCACT<br>GACTATTGTAGGTACCTTGCCCTTTACCTACATGCTGGAGAAGTGGAGGTGGATGGTATTTAAG<br>GGGGAGATACCGAAAGATCAATGGATGAAAAAGTGGTGGGAAATGAAAAGGGAGATCGTTGGCG<br>TAGTTGAACCAGTACCGCATGATGAGACGTACTGCGATCCGGCTAGTCTGTTCCATGTCTCTAA<br>TGATTACTCTTTCATCCGCTACTACACCCGCACGCTGTATCAATTCCAGTTCCAAGAAGCTCTC<br>TGTCAGGCTGCCAAGCACGAAGGACCGCTGCACAAATGCGACATTAGCAATTCTACAGAGGCGG<br>GTCAGAAGTTGTTCAATATGCTTAGACTGGGGAAGAGCGAACCGTGGACGCTCGCTTTGGAGAA<br>CGTTGTTGGAGCTAAGAATATGAACGTCAGGCCCTTGCTGAATTACTTTGAACCTCTGTTTACG<br>TGGTTGAAAGACCAAAATAAAAACTCCTTTGTTGGGTGGAGTACTGACTGGTCCCCCTATGCG |
| 208 | CAATCTACCATCGAAGAGCAGGCCAAAACATTCCTCGACAAGTTTAATCACGAGGCTGAAGACC<br>TTTTCTACCAATCAAGTCTGGCTAGCTGGAATTACAATACAAACATTACAGAGGAGAACGTACA<br>AAACATGAATAACGCAGGGGACAAGTGGAGCGCATTCCTTAAGGAACAAAGTACCCTTGCGCAA<br>ATGTATCCGCTGCAAGAGATTCAAAACCTGACGGTTAAGCTGCAACTTCAGGCCCTCCAACAAA<br>ATGGAAGTTCAGTCTTGTCAGAAGACAAAAGCAAGCGACTGAACACCATCCTTAACACCATGTC<br>AACCATATATTCAACAGGTAAAGTTTGCAATCCGGATAACCCCCAAGAATGTTTGCTTCTTGAA<br>CCCGGTCTCAACGAAATTATGGCCAACAGTCTTGATTACAACGAGCGATTGTGGGCATGGGAAA<br>GTTGGAGGAGTGAGGTAGGCAAACAGTTGAGACCTCTTTATGAAGAGTACGTTGTCCTTAAAAA<br>TGAAATGGCTCGCGCGAATCATTATGAAGACTATGGTGACTACTGGAGGGGGGATTATGAGGTG<br>AACGGGGTGGACGGATACGATTACTCTAGGGGCCAGCTGATAGAGGATGTCGAGCACACCTTTG<br>AGGAGATTAAGCCGTTGTACGAACATTTGCACGCCTATGTCAGGGCTAAGCTCATGAACGCTTA<br>TCCGAGTTATATCTCCCCGATAGGATGCTTGCCTGCTCACTTGTTGGGCGATATGTGGGGACGC<br>TTTTGGACCAACTTGTATTCCCTTACGGTACCGTTCGGCCAGAAACCAAATATCGACGTGACAG<br>ACGCAATGGTGGATCAAGCATGGGATGCGCAACGAATCTTCAAGGAGGCAGAAAAATTTTTCGT<br>TTCAGTTGGACTCCCAAACATGACGCAGGGTTTCTGGGAGAACTCAATGTTGACAGATCCAGGT<br>AATGTGCAGAAAGCGGTTTGCCACCCTACTGCATGGGATCTTGGTAAAGGGGACTTCCGCATAC<br>TCATGTGTACGAAAGTAACTATGGACGACTTTCTTACTGCGCACCACGAGATGGGGCACATACA<br>ATACGATATGGCGTACGCAGCTCAACCTTTCCTTCTGCGGAACGGGGCGAATGAAGGATTTCAC |

| | |
|---|---|
| | GAGGCAGTGGGTGAGATTATGTCCCTGTCAGCTGCCACTCCGAAACATCTGAAAAGCATCGGCC<br>TGTTGAGCCCAGACTTCCAAGAAGATAATGAGACCGAAATAAACTTCCTTCTGAAGCAAGCACT<br>GACTATTGTAGGTACCTTGCCCTTTACATACATGCTGGAGAAGTGGAGGTGGATGGTATTTAAG<br>GGGGAGATACCGAAAGATCAATGGATGAAAAAGTGGTGGGAAATGAAAAGGGAGATCGTTGGCG<br>TAGTTGAACCAGTACCGCATGATGAGACGTACTGCGATCCGGCTAGTCTGTTCCATGTCTCTAA<br>TGATTACTCTTTCATCCGCTACTACACCCGCACGCTGTATCAATTCCAGTTCCAAGAAGCTCTC<br>TGTCAGGCTGCCAAGCACGAAGGACCGCTGCACAAATGCGACATTAGCAATTCTACAGAGGCGG<br>GTCAGAAGTTGTTCAATATGCTTAGACTGGGGAAGAGCGAACCGTGGACGCTCGCTTTGGAGAA<br>CGTTGTTGGAGCTAAGAATATGAACGTCAGGCCCTTGCTGAATTACTTTGAACCTCTGTTTACG<br>TGGTTGAAAGACCAAAATAAAAACTCCTTTGTTGGGTGGAGTACTGACTGGTCCCCCTATGCGG<br>ACCAAAGCATCAAAGTGAGGATAAGCCTAAAATCAGCTCTTGGAGATAAAGCATATGAATGGAA<br>CGACAATGAAATGTACCTGTTCCGATCATCTGTTGCATATGCTATGAGGCAGTACTTTTTAAAA<br>GTAAAAAATCAGATGATTCTTTTTGGGGAGGAGGATGTGCGAGTGGCTAATTTGAAACCAAGAA<br>TCTCCTTTAATTTCTTTGTCACTGCACCTAAAAATGTGTCTGATATCATTCCTAGAACTGAAGT<br>TGAAAAGGCCATCAGGATGTCCCGGAGCCGTATCAATGATGCTTTCCGTCTGAATGACAACAGC<br>CTAGAGTTTCTGGGGATACAGCCAACACTTGGACCTCCTAACCAGCCCCCTGTTTCC |
| Y208 | CAATCTACCATCGAAGAGCAGGCCAAAACATTCCTCGACAAGTTTAATCACGAGGCTGAAGACC<br>TTTTCTACCAATCAAGTCTGGCTAGCTGGAATTACAATACAAACATTACAGAGGAGAACGTACA<br>AAACATGAATAACGCAGGGGACAAGTGGAGCGCATTCCTTAAGGAACAAAGTACCCTTGCGCAA<br>ATGTATCCGCTGCAAGAGATTCAAAACCTGACGGTTAAGCTGCAACTTCAGGCCCTCCAACAAA<br>ATGGATCCTCAGTCTTGTCAGAAGACAAAAGCAAGCGACTGAACACCATCCTTAACACCATGTC<br>AACCATATATTCAACAGGTAAAGTTTGCAATCCGGATAACCCCCAAGAATGTTTGCTTCTTGAA<br>CCCGGTCTCAACGAAATTATGGCCAACAGTCTTGATTACAACGAGCGATTGTGGGCATGGGAAA<br>GTTGGAGGAGTGAGGTAGGCAAACAGTTGAGACCTCTTTATGAAGAGTACGTTGTCCTTAAAAA<br>TGAAATGGCTCGCGCGAATCATTATGAAGACTATGGTGACTACTGGAGGGGGGATTATGAGGTG<br>AACGGGGTGGACGGATACGATTACTCTAGGGGCCAGCTGATAGAGGATGTCGAGCACACCTTTG<br>AGGAGATTAAGCCGTTGTACGAACATTTGCACGCCTATGTCAGGGCTAAGCTCATGAACGCTTA<br>TCCGAGTTATATCTCCCCGATAGGATGCTTGCCTGCTCACTTGTTGGGCGATATGTGGGGACGC<br>TTTTGGACCAACTTGTATTCCCTTACGGTACCGTTCGGCCAGAAACCAAATATCGACGTGACAG<br>ACGCAATGGTGGATCAAGCATGGGATGCGCAACGAATCTTCAAGGAGGCAGAAAAATTTTTCGT<br>TTCAGTTGGACTCCCAAACATGACGCAGGGTTTCTGGGAGAACTCAATGTTGACAGATCCAGGT<br>AATGTGCAGAAAGCGGTTTGCCACCCTACTGCATGGGATCTTGGTAAAGGGGACTTCCGCATAC<br>TCATGTGTACGAAAGTAACTATGGACGACTTTCTTACTGCGCACCACGAGATGGGGCACATACA<br>ATACGATATGGCGTACGCAGCTCAACCTTTCCTTCTGCGGAACGGGGCGAATGAAGGATTTCAC<br>GAGGCAGTGGGTGAGATTATGTCCCTGTCAGCTGCCACTCCGAAACATCTGAAAAGCATCGGCC<br>TGTTGAGCCCAGACTTCCAAGAAGATAATGAGACCGAAATAAACTTCCTTCTGAAGCAAGCACT<br>GACTATTGTAGGTACCTTGCCCTTTACATACATGCTGGAGAAGTGGAGGTGGATGGTATTTAAG<br>GGGGAGATACCGAAAGATCAATGGATGAAAAAGTGGTGGGAAATGAAAAGGGAGATCGTTGGCG<br>TAGTTGAACCAGTACCGCATGATGAGACGTACTGCGATCCGGCTAGTCTGTTCCATGTCTCTAA<br>TGATTACTCTTTCATCCGCTACTACACCCGCACGCTGTATCAATTCCAGTTCCAAGAAGCTCTC<br>TGTCAGGCTGCCAAGCACGAAGGACCGCTGCACAAATGCGACATTAGCAATTCTACAGAGGCGG<br>GTCAGAAGTTGTTCAATATGCTTAGACTGGGGAAGAGCGAACCGTGGACGCTCGCTTTGGAGAA<br>CGTTGTTGGAGCTAAGAATATGAACGTCAGGCCCTTGCTGAATTACTTTGAACCTCTGTTTACG<br>TGGTTGAAAGACCAAAATAAAAACTCCTTTGTTGGGTGGAGTACTGACTGGTCCCCCTATGCG |
| 14 | CAATCTACCATCGAAGAGCAGGCCAAAACATTCCTCGACAAGTTTAATGTCGAGGCTGAAGACC<br>TTTTCTACCAATCAAGTCTGGCTAGCTGGAATTACAATACAAACATTACAGAGGAGAACGTACA<br>AAACATGAATAACGCAGGGGACAAGTGGAGCGCATTCCTTAAGGAACAAAGTACCCTTGCGCAA |

| | |
|---|---|
| | ATGTATCCGCTGCAAGAGATTCAAAACCTGACGGTTAAGCTGCAACTTCAGGCCCTCCAACAAA<br>ATGGAAGTTCAGTCTTGTCAGAAGACAAAAGCAAGCGACTGAACACCATCCTTAACACCATGTC<br>AACCATATATTCAACAGGTAAAGTTTGCAATCCGGATAACCCCCAAGAATGTTTGCTTCTTGAA<br>CCCGGTCTCAACGAAATTATGGCCAACAGTCTTGATTACAACGAGCGATTGTGGGCATGGGAAA<br>GTTGGAGGAGTGAGGTAGGCAAACAGTTGAGACCTCTTTATGAAGAGTACGTTGTCCTTAAAAA<br>TGAAATGGCTCGCGCGAATCATTATGAAGACTATGGTGACTACTGGAGGGGGGATTATGAGGTG<br>AACGGGGTGGACGGATACGATTACTCTAGGGGCCAGCTGATAGAGGATGTCGAGCACACCTTTG<br>AGGAGATTAAGCCGTTGTACGAACATTTGCACGCCTATGTCAGGGCTAAGCTCATGAACGCTTA<br>TCCGAGTTATATCTCCCCGATAGGATGCTTGCCTGCTCACTTGTTGGGCGATATGTGGGGACGC<br>TTTTGGACCAACTTGTATTCCCTTACGGTACCGTTCGGCCAGAAACCAAATATCGACGTGACAG<br>ACGCAATGGTGGATCAAGCATGGGATGCGCAACGAATCTTCAAGGAGGCAGAAAAATTTTTCGT<br>TTCAGTTGGACTCCCAAACATGACGCAGGGTTTCTGGGAGAACTCAATGTTGACAGATCCAGGT<br>AATGTGCAGAAAGCGGTTTGCCACCCTACTGCATGGGATCTTGGTAAAGGGGACTTCCGCATAC<br>TCATGTGTACGAAAGTAACTATGGACGACTTTCTTACTGCGCACCACGAGATGGGGCACATACA<br>ATACGATATGGCGTACGCAGCTCAACCTTTCCTTCTGCGGAACGGGGCGAATGAAGGATTTCAC<br>GAGGCAGTGGGTGAGATTATGTCCCTGTCAGCTGCCACTCCGAAACATCTGAAAAGCATCGGCC<br>TGTTGAGCCCAGACTTCCAAGAAGATAATGAGACCGAAATAAACTTCCTTCTGAAGCAAGCACT<br>GACTATTGTAGGTACCTTGCCCTTTACCTACATGCTGGAGAAGTGGAGGTGGATGGTATTTAAG<br>GGGGAGATACCGAAAGATCAATGGATGAAAAAGTGGTGGGAAATGAAAAGGGAGATCGTTGGCG<br>TAGTTGAACCAGTACCGCATGATGAGACGTACTGCGATCCGGCTAGTCTGTTCCATGTCTCTAA<br>TGATTACTCTTTCATCCGCTACTACACCCGCACGCTGTATCAATTCCAGTTCCAAGAAGCTCTC<br>TGTCAGGCTGCCAAGCACGAAGGACCGCTGCACAAATGCGACATTAGCAATTCTACAGAGGCGG<br>GTCAGAAGTTGTTCAATATGCTTAGACTGGGGAAGAGCGAACCGTGGACGCTCGCTTTGGAGAA<br>CGTTGTTGGAGCTAAGAATATGAACGTCAGGCCCTTGCTGAATTACTTTGAACCTCTGTTTACG<br>TGGTTGAAAGACCAAAATAAAAACTCCTTTGTTGGGTGGAGTACTGACTGGTCCCCCTATGCG |
| 295 | CAATCTACCATCGAAGAGCAGGCCAAAACATTCCTCGACAAGTTTAATGTCGAGGCTGAAGACC<br>TTTTCTACCAATCAAGTCTGGCTAGCTGGAATTACAATACAAACATTACAGAGGAGAACGTACA<br>AAACATGAATAACGCAGGGGACAAGTGGAGCGCATTCCTTAAGGAACAAAGTACCCTTGCGCAA<br>ATGTATCCGCTGCAAGAGATTCAAAACCTGACGGTTAAGCTGCAACTTCAGGCCCTCCAACAAA<br>ATGGATCCTCAGTCTTGTCAGAAGACAAAAGCAAGCGACTGAACACCATCCTTAACACCATGTC<br>AACCATATATTCAACAGGTAAAGTTTGCAATCCGGATAACCCCCAAGAATGTTTGCTTCTTGAA<br>CCCGGTCTCAACGAAATTATGGCCAACAGTCTTGATTACAACGAGCGATTGTGGGCATGGGAAA<br>GTTGGAGGAGTGAGGTAGGCAAACAGTTGAGACCTCTTTATGAAGAGTACGTTGTCCTTAAAAA<br>TGAAATGGCTCGCGCGAATCATTATGAAGACTATGGTGACTACTGGAGGGGGGATTATGAGGTG<br>AACGGGGTGGACGGATACGATTACTCTAGGGGCCAGCTGATAGAGGATGTCGAGCACACCTTTG<br>AGGAGATTAAGCCGTTGTACGAACATTTGCACGCCTATGTCAGGGCTAAGCTCATGAACGCTTA<br>TCCGAGTTATATCTCCCCGATAGGATGCTTGCCTGCTCACTTGTTGGGCGATATGTGGGGACGC<br>TTTTGGACCAACTTGTATTCCCTTACGGTACCGTTCGGCCAGAAACCAAATATCGACGTGACAG<br>ACGCAATGGTGGATCAAGCATGGGATGCGCAACGAATCTTCAAGGAGGCAGAAAAATTTTTCGT<br>TTCAGTTGGACTCCCAAACATGACGCAGGGTTTCTGGGAGAACTCAATGTTGACAGATCCAGGT<br>AATGTGCAGAAAGCGGTTTGCCACCCTACTGCATGGGATCTTGGTAAAGGGGACTTCCGCATAC<br>TCATGTGTACGAAAGTAACTATGGACGACTTTCTTACTGCGCACCACGAGATGGGGCACATACA<br>ATACGATATGGCGTACGCAGCTCAACCTTTCCTTCTGCGGAACGGGGCGAATGAAGGATTTCAC<br>GAGGCAGTGGGTGAGATTATGTCCCTGTCAGCTGCCACTCCGAAACATCTGAAAAGCATCGGCC<br>TGTTGAGCCCAGACTTCCAAGAAGATAATGAGACCGAAATAAACTTCCTTCTGAAGCAAGCACT<br>GACTATTGTAGGTACCTTGCCCTTTACATACATGCTGGAGAAGTGGAGGTGGATGGTATTTAAG<br>GGGGAGATACCGAAAGATCAATGGATGAAAAAGTGGTGGGAAATGAAAAGGGAGATCGTTGGCG |

| | |
|---|---|
| | TAGTTGAACCAGTACCGCATGATGAGACGTACTGCGATCCGGCTAGTCTGTTCCATGTCTCTAA<br>TGATTACTCTTTCATCCGCTACTACACCCGCACGCTGTATCAATTCCAGTTCCAAGAAGCTCTC<br>TGTCAGGCTGCCAAGCACGAAGGACCGCTGCACAAATGCGACATTAGCAATTCTACAGAGGCGG<br>GTCAGAAGTTGTTCAATATGCTTAGACTGGGGAAGAGCGAACCGTGGACGCTCGCTTTGGAGAA<br>CGTTGTTGGAGCTAAGAATATGAACGTCAGGCCCTTGCTGAATTACTTTGAACCTCTGTTTACG<br>TGGTTGAAAGACCAAAATAAAAACTCCTTTGTTGGGTGGAGTACTGACTGGTCCCCCTATGCGG<br>ACCAAAGCATCAAAGTGAGGATAAGCCTAAAATCAGCTCTTGGAGATAAAGCATATGAATGGAA<br>CGACAATGAAATGTACCTGTTCCGATCATCTGTTGCATATGCTATGAGGCAGTACTTTTTAAAA<br>GTAAAAAATCAGATGATTCTTTTTGGGGAGGAGGATGTGCGAGTGGCTAATTTGAAACCAAGAA<br>TCTCCTTTAATTTCTTTGTCACTGCACCTAAAAATGTGTCTGATATCATTCCTAGAACTGAAGT<br>TGAAAAGGCCATCAGGATGTCCCGGAGCCGTATCAATGATGCTTTCCGTCTGAATGACAACAGC<br>CTAGAGTTTCTGGGGATACAGCCAACACTTGGACCTCCTAACCAGCCCCCTGTTTCC |
| 19 | CAATCTACCATCGAAGAGCAGGCCAAAACATTCCTCGACTTCTTTAATATCCAGGCTGAAGACC<br>TTTTCTACCAATCAAGTCTGGCTAGCTGGAATTACAATACAAACATTACAGAGGAGAACGTACA<br>AAACATGAATAACGCAGGGGACAAGTGGAGCGCATTCCTTAAGGAACAAAGTACCCTTGCGCAA<br>ATGTATCCGCTGCAAGAGATTCAAAACCTGACGGTTAAGCTGCAACTTCAGGCCCTCCAACAAA<br>ATGGAAGTTCAGTCTTGTCAGAAGACAAAAGCAAGCGACTGAACACCATCCTTAACACCATGTC<br>AACCATATATTCAACAGGTAAAGTTTGCAATCCGGATAACCCCCAAGAATGTTTGCTTCTTGAA<br>CCCGGTCTCAACGAAATTATGGCCAACAGTCTTGATTACAACGAGCGATTGTGGGCATGGGAAA<br>GTTGGAGGAGTGAGGTAGGCAAACAGTTGAGACCTCTTTATGAAGAGTACGTTGTCCTTAAAAA<br>TGAAATGGCTCGCGCGAATCATTATGAAGACTATGGTGACTACTGGAGGGGGGATTATGAGGTG<br>AACGGGGTGGACGGATACGATTACTCTAGGGGCCAGCTGATAGAGGATGTCGAGCACACCTTTG<br>AGGAGATTAAGCCGTTGTACGAACATTTGCACGCCTATGTCAGGGCTAAGCTCATGAACGCTTA<br>TCCGAGTTATATCTCCCCGATAGGATGCTTGCCTGCTCACTTGTTGGGCGATATGTGGGGACGC<br>TTTTGGACCAACTTGTATTCCCTTACGGTACCGTTCGGCCAGAAACCAAATATCGACGTGACAG<br>ACGCAATGGTGGATCAAGCATGGGATGCGCAACGAATCTTCAAGGAGGCAGAAAAATTTTTCGT<br>TTCAGTTGGACTCCCAAACATGACGCAGGGTTTCTGGGAGAACTCAATGTTGACAGATCCAGGT<br>AATGTGCAGAAAGCGGTTTGCCACCCTACTGCATGGGATCTTGGTAAAGGGGACTTCCGCATAC<br>TCATGTGTACGAAAGTAACTATGGACGACTTTCTTACTGCGCACCACGAGATGGGGCACATACA<br>ATACGATATGGCGTACGCAGCTCAACCTTTCCTTCTGCGGAACGGGGCGAATGAAGGATTTCAC<br>GAGGCAGTGGGTGAGATTATGTCCCTGTCAGCTGCCACTCCGAAACATCTGAAAAGCATCGGCC<br>TGTTGAGCCCAGACTTCCAAGAAGATAATGAGACCGAAATAAACTTCCTTCTGAAGCAAGCACT<br>GACTATTGTAGGTACCTTGCCCTTTACCTACATGCTGGAGAAGTGGAGGTGGATGGTATTTAAG<br>GGGGAGATACCGAAAGATCAATGGATGAAAAAGTGGTGGGAAATGAAAAGGGAGATCGTTGGCG<br>TAGTTGAACCAGTACCGCATGATGAGACGTACTGCGATCCGGCTAGTCTGTTCCATGTCTCTAA<br>TGATTACTCTTTCATCCGCTACTACACCCGCACGCTGTATCAATTCCAGTTCCAAGAAGCTCTC<br>TGTCAGGCTGCCAAGCACGAAGGACCGCTGCACAAATGCGACATTAGCAATTCTACAGAGGCGG<br>GTCAGAAGTTGTTCAATATGCTTAGACTGGGGAAGAGCGAACCGTGGACGCTCGCTTTGGAGAA<br>CGTTGTTGGAGCTAAGAATATGAACGTCAGGCCCTTGCTGAATTACTTTGAACCTCTGTTTACG<br>TGGTTGAAAGACCAAAATAAAAACTCCTTTGTTGGGTGGAGTACTGACTGGTCCCCCTATGCG |
| 293 | CAATCTACCATCGAAGAGCAGGCCAAAACATTCCTCGACTTCTTTAATATCCAGGCTGAAGACC<br>TTTTCTACCAATCAAGTCTGGCTAGCTGGAATTACAATACAAACATTACAGAGGAGAACGTACA<br>AAACATGAATAACGCAGGGGACAAGTGGAGCGCATTCCTTAAGGAACAAAGTACCCTTGCGCAA<br>ATGTATCCGCTGCAAGAGATTCAAAACCTGACGGTTAAGCTGCAACTTCAGGCCCTCCAACAAA<br>ATGGAAGTTCAGTCTTGTCAGAAGACAAAAGCAAGCGACTGAACACCATCCTTAACACCATGTC<br>AACCATATATTCAACAGGTAAAGTTTGCAATCCGGATAACCCCCAAGAATGTTTGCTTCTTGAA<br>CCCGGTCTCAACGAAATTATGGCCAACAGTCTTGATTACAACGAGCGATTGTGGGCATGGGAAA |

| | |
|---|---|
| | GTTGGAGGAGTGAGGTAGGCAAACAGTTGAGACCTCTTTATGAAGAGTACGTTGTCCTTAAAAA<br>TGAAATGGCTCGCGCGAATCATTATGAAGACTATGGTGACTACTGGAGGGGGGATTATGAGGTG<br>AACGGGGTGGACGGATACGATTACTCTAGGGGCCAGCTGATAGAGGATGTCGAGCACACCTTTG<br>AGGAGATTAAGCCGTTGTACGAACATTTGCACGCCTATGTCAGGGCTAAGCTCATGAACGCTTA<br>TCCGAGTTATATCTCCCCGATAGGATGCTTGCCTGCTCACTTGTTGGGCGATATGTGGGGACGC<br>TTTTGGACCAACTTGTATTCCCTTACGGTACCGTTCGGCCAGAAACCAAATATCGACGTGACAG<br>ACGCAATGGTGGATCAAGCATGGGATGCGCAACGAATCTTCAAGGAGGCAGAAAAATTTTTCGT<br>TTCAGTTGGACTCCCAAACATGACGCAGGGTTTCTGGGAGAACTCAATGTTGACAGATCCAGGT<br>AATGTGCAGAAAGCGGTTTGCCACCCTACTGCATGGGATCTTGGTAAAGGGGACTTCCGCATAC<br>TCATGTGTACGAAAGTAACTATGGACGACTTTCTTACTGCGCACCACGAGATGGGGCACATACA<br>ATACGATATGGCGTACGCAGCTCAACCTTTCCTTCTGCGGAACGGGGCGAATGAAGGATTTCAC<br>GAGGCAGTGGGTGAGATTATGTCCCTGTCAGCTGCCACTCCGAAACATCTGAAAAGCATCGGCC<br>TGTTGAGCCCAGACTTCCAAGAAGATAATGAGACCGAAATAAACTTCCTTCTGAAGCAAGCACT<br>GACTATTGTAGGTACCTTGCCCTTTACATACATGCTGGAGAAGTGGAGGTGGATGGTATTTAAG<br>GGGGAGATACCGAAAGATCAATGGATGAAAAAGTGGTGGGAAATGAAAAGGGAGATCGTTGGCG<br>TAGTTGAACCAGTACCGCATGATGAGACGTACTGCGATCCGGCTAGTCTGTTCCATGTCTCTAA<br>TGATTACTCTTTCATCCGCTACTACACCCGCACGCTGTATCAATTCCAGTTCCAAGAAGCTCTC<br>TGTCAGGCTGCCAAGCACGAAGGACCGCTGCACAAATGCGACATTAGCAATTCTACAGAGGCGG<br>GTCAGAAGTTGTTCAATATGCTTAGACTGGGGAAGAGCGAACCGTGGACGCTCGCTTTGGAGAA<br>CGTTGTTGGAGCTAAGAATATGAACGTCAGGCCCTTGCTGAATTACTTTGAACCTCTGTTTACG<br>TGGTTGAAAGACCAAAATAAAAACTCCTTTGTTGGGTGGAGTACTGACTGGTCCCCCTATGCGG<br>ACCAAAGCATCAAAGTGAGGATAAGCCTAAAATCAGCTCTTGGAGATAAAGCATATGAATGGAA<br>CGACAATGAAATGTACCTGTTCCGATCATCTGTTGCATATGCTATGAGGCAGTACTTTTTAAAA<br>GTAAAAAATCAGATGATTCTTTTTGGGGAGGAGGATGTGCGAGTGGCTAATTTGAAACCAAGAA<br>TCTCCTTTAATTTCTTTGTCACTGCACCTAAAAATGTGTCTGATATCATTCCTAGAACTGAAGT<br>TGAAAAGGCCATCAGGATGTCCCGGAGCCGTATCAATGATGCTTTCCGTCTGAATGACAACAGC<br>CTAGAGTTTCTGGGGATACAGCCAACACTTGGACCTCCTAACCAGCCCCCTGTTTCC |
| 117 | CAATCTACCATCGAAGAGCAGGCCAAAACATTCCTCGACAAGTTTAATCACGAGGCTGAAGACC<br>TTTTCTACCAATCAAGTCTGGCTAGCTGGAATTACAATACAAACATTACAGAGGAGAACGTACA<br>AAACATGAATAACGCAGGGGACAAGTGGAGCGCATTCCTTAAGGAACAAAGTACCCTTGCGCAA<br>ATGTATCCGCTGCAAGAGATTCAACAACTGACGGTTAAGCTGCAACTTCAGGCCCTCCAACAAA<br>ATGGATCCTCAGTCTTGTCAGAAGACAAAAGCAAGCGACTGAACACCATCCTTAACACCATGTC<br>AACCATATATTCAACAGGTAAAGTTTGCAATCCGGATAACCCCCAAGAATGTTTGCTTCTTGAA<br>CCCGGTCTCAACGAAATTATGGCCAACAGTCTTGATTACAACGAGCGATTGTGGGCATGGGAAA<br>GTTGGAGGAGTGAGGTAGGCAAACAGTTGAGACCTCTTTATGAAGAGTACGTTGTCCTTAAAAA<br>TGAAATGGCTCGCGCGAATCATTATGAAGACTATGGTGACTACTGGAGGGGGGATTATGAGGTG<br>AACGGGGTGGACGGATACGATTACTCTAGGGGCCAGCTGATAGAGGATGTCGAGCACACCTTTG<br>AGGAGATTAAGCCGTTGTACGAACATTTGCACGCCTATGTCAGGGCTAAGCTCATGAACGCTTA<br>TCCGAGTTATATCTCCCCGATAGGATGCTTGCCTGCTCACTTGTTGGGCGATATGTGGGGACGC<br>TTTTGGACCAACTTGTATTCCCTTACGGTACCGTTCGGCCAGAAACCAAATATCGACGTGACAG<br>ACGCAATGGTGGATCAAGCATGGGATGCGCAACGAATCTTCAAGGAGGCAGAAAAATTTTTCGT<br>TTCAGTTGGACTCCCAAACATGACGCAGGGTTTCTGGGAGAACTCAATGTTGACAGATCCAGGT<br>AATGTGCAGAAAGCGGTTTGCCACCCTACTGCATGGGATCTTGGTAAAGGGGACTTCCGCATAC<br>TCATGTGTACGAAAGTAACTATGGACGACTTTCTTACTGCGCACAACGAGATGGGGAACATACA<br>ATACGATATGGCGTACGCAGCTCAACCTTTCCTTCTGCGGAACGGGGCGAATGAAGGATTTCAC<br>GAGGCAGTGGGTGAGATTATGTCCCTGTCAGCTGCCACTCCGAAACATCTGAAAAGCATCGGCC<br>TGTTGAGCCCAGACTTCCAAGAAGATAATGAGACCGAAATAAACTTCCTTCTGAAGCAAGCACT |

| | |
|---|---|
| | GACTATTGTAGGTACCTTGCCCTTTACATACATGCTGGAGAAGTGGAGGTGGATGGTATTTAAG<br>GGGGAGATACCGAAAGATCAATGGATGAAAAAGTGGTGGGAAATGAAAAGGGAGATCGTTGGCG<br>TAGTTGAACCAGTACCGCATGATGAGACGTACTGCGATCCGGCTAGTCTGTTCCATGTCTCTAA<br>TGATTACTCTTTCATCCGCTACTACACCCGCACGCTGTATCAATTCCAGTTCCAAGAAGCTCTC<br>TGTCAGGCTGCCAAGCACGAAGGACCGCTGCACAAATGCGACATTAGCAATTCTACAGAGGCGG<br>GTCAGAAGTTGTTCAATATGCTTAGACTGGGGAAGAGCGAACCGTGGACGCTCGCTTTGGAGAA<br>CGTTGTTGGAGCTAAGAATATGAACGTCAGGCCCTTGCTGAATTACTTTGAACCTCTGTTTACG<br>TGGTTGAAAGACCAAAATAAAAACTCCTTTGTTGGGTGGAGTACTGACTGGTCCCCCTATGCG |
| Y117 | CAATCTACCATCGAAGAGCAGGCCAAAACATTCCTCGACAAGTTTAATCACGAGGCTGAAGACC<br>TTTTCTACCAATCAAGTCTGGCTAGCTGGAATTACAATACAAACATTACAGAGGAGAACGTACA<br>AAACATGAATAACGCAGGGGACAAGTGGAGCGCATTCCTTAAGGAACAAAGTACCCTTGCGCAA<br>ATGTATCCGCTGCAAGAGATTCAACAACTGACGGTTAAGCTGCAACTTCAGGCCCTCCAACAAA<br>ATGGATCCTCAGTCTTGTCAGAAGACAAAAGCAAGCGACTGAACACCATCCTTAACACCATGTC<br>AACCATATATTCAACAGGTAAAGTTTGCAATCCGGATAACCCCCAAGAATGTTTGCTTCTTGAA<br>CCCGGTCTCAACGAAATTATGGCCAACAGTCTTGATTACAACGAGCGATTGTGGGCATGGGAAA<br>GTTGGAGGAGTGAGGTAGGCAAACAGTTGAGACCTCTTTATGAAGAGTACGTTGTCCTTAAAAA<br>TGAAATGGCTCGCGCGAATCATTATGAAGACTATGGTGACTACTGGAGGGGGGATTATGAGGTG<br>AACGGGGTGGACGGATACGATTACTCTAGGGGCCAGCTGATAGAGGATGTCGAGCACACCTTTG<br>AGGAGATTAAGCCGTTGTACGAACATTTGCACGCCTATGTCAGGGCTAAGCTCATGAACGCTTA<br>TCCGAGTTATATCTCCCCGATAGGATGCTTGCCTGCTCACTTGTTGGGCGATATGTGGGGACGC<br>TTTTGGACCAACTTGTATTCCCTTACGGTACCGTTCGGCCAGAAACCAAATATCGACGTGACAG<br>ACGCAATGGTGGATCAAGCATGGGATGCGCAACGAATCTTCAAGGAGGCAGAAAAATTTTTCGT<br>TTCAGTTGGACTCCCAAACATGACGCAGGGTTTCTGGGAGAACTCAATGTTGACAGATCCAGGT<br>AATGTGCAGAAAGCGGTTTGCCACCCTACTGCATGGGATCTTGGTAAAGGGGACTTCCGCATAC<br>TCATGTGTACGAAAGTAACTATGGACGACTTTCTTACTGCGCACCACGAGATGGGGCACATACA<br>ATACGATATGGCGTACGCAGCTCAACCTTTCCTTCTGCGGAACGGGGCGAATGAAGGATTTCAC<br>GAGGCAGTGGGTGAGATTATGTCCCTGTCAGCTGCCACTCCGAAACATCTGAAAAGCATCGGCC<br>TGTTGAGCCCAGACTTCCAAGAAGATAATGAGACCGAAATAAACTTCCTTCTGAAGCAAGCACT<br>GACTATTGTAGGTACCTTGCCCTTTACATACATGCTGGAGAAGTGGAGGTGGATGGTATTTAAG<br>GGGGAGATACCGAAAGATCAATGGATGAAAAAGTGGTGGGAAATGAAAAGGGAGATCGTTGGCG<br>TAGTTGAACCAGTACCGCATGATGAGACGTACTGCGATCCGGCTAGTCTGTTCCATGTCTCTAA<br>TGATTACTCTTTCATCCGCTACTACACCCGCACGCTGTATCAATTCCAGTTCCAAGAAGCTCTC<br>TGTCAGGCTGCCAAGCACGAAGGACCGCTGCACAAATGCGACATTAGCAATTCTACAGAGGCGG<br>GTCAGAAGTTGTTCAATATGCTTAGACTGGGGAAGAGCGAACCGTGGACGCTCGCTTTGGAGAA<br>CGTTGTTGGAGCTAAGAATATGAACGTCAGGCCCTTGCTGAATTACTTTGAACCTCTGTTTACG<br>TGGTTGAAAGACCAAAATAAAAACTCCTTTGTTGGGTGGAGTACTGACTGGTCCCCCTATGCG |
| 118 | CAATCTACCATCGAAGAGCAGGCCAAAACATTCCTCGACAAGTTTAATGTCGAGGCTGAAGACC<br>TTTTCTACCAATCAAGTCTGGCTAGCTGGAATTACAATACAAACATTACAGAGGAGAACGTACA<br>AAACATGAATAACGCAGGGGACAAGTGGAGCGCATTCCTTAAGGAACAAAGTACCCTTGCGCAA<br>ATGTATCCGCTGCAAGAGATTCAACAACTGACGGTTAAGCTGCAACTTCAGGCCCTCCAACAAA<br>ATGGATCCTCAGTCTTGTCAGAAGACAAAAGCAAGCGACTGAACACCATCCTTAACACCATGTC<br>AACCATATATTCAACAGGTAAAGTTTGCAATCCGGATAACCCCCAAGAATGTTTGCTTCTTGAA<br>CCCGGTCTCAACGAAATTATGGCCAACAGTCTTGATTACAACGAGCGATTGTGGGCATGGGAAA<br>GTTGGAGGAGTGAGGTAGGCAAACAGTTGAGACCTCTTTATGAAGAGTACGTTGTCCTTAAAAA<br>TGAAATGGCTCGCGCGAATCATTATGAAGACTATGGTGACTACTGGAGGGGGGATTATGAGGTG<br>AACGGGGTGGACGGATACGATTACTCTAGGGGCCAGCTGATAGAGGATGTCGAGCACACCTTTG<br>AGGAGATTAAGCCGTTGTACGAACATTTGCACGCCTATGTCAGGGCTAAGCTCATGAACGCTTA |

| | |
|---|---|
| | TCCGAGTTATATCTCCCCGATAGGATGCTTGCCTGCTCACTTGTTGGGCGATATGTGGGGACGC<br>TTTTGGACCAACTTGTATTCCCTTACGGTACCGTTCGGCCAGAAACCAAATATCGACGTGACAG<br>ACGCAATGGTGGATCAAGCATGGGATGCGCAACGAATCTTCAAGGAGGCAGAAAAATTTTTCGT<br>TTCAGTTGGACTCCCAAACATGACGCAGGGTTTCTGGGAGAACTCAATGTTGACAGATCCAGGT<br>AATGTGCAGAAAGCGGTTTGCCACCCTACTGCATGGGATCTTGGTAAAGGGGACTTCCGCATAC<br>TCATGTGTACGAAAGTAACTATGGACGACTTTCTTACTGCGCACAACGAGATGGGGAACATACA<br>ATACGATATGGCGTACGCAGCTCAACCTTTCCTTCTGCGGAACGGGGCGAATGAAGGATTTCAC<br>GAGGCAGTGGGTGAGATTATGTCCCTGTCAGCTGCCACTCCGAAACATCTGAAAAGCATCGGCC<br>TGTTGAGCCCAGACTTCCAAGAAGATAATGAGACCGAAATAAACTTCCTTCTGAAGCAAGCACT<br>GACTATTGTAGGTACCTTGCCCTTTACATACATGCTGGAGAAGTGGAGGTGGATGGTATTTAAG<br>GGGGAGATACCGAAAGATCAATGGATGAAAAAGTGGTGGGAAATGAAAAGGGAGATCGTTGGCG<br>TAGTTGAACCAGTACCGCATGATGAGACGTACTGCGATCCGGCTAGTCTGTTCCATGTCTCTAA<br>TGATTACTCTTTCATCCGCTACTACACCCGCACGCTGTATCAATTCCAGTTCCAAGAAGCTCTC<br>TGTCAGGCTGCCAAGCACGAAGGACCGCTGCACAAATGCGACATTAGCAATTCTACAGAGGCGG<br>GTCAGAAGTTGTTCAATATGCTTAGACTGGGGAAGAGCGAACCGTGGACGCTCGCTTTGGAGAA<br>CGTTGTTGGAGCTAAGAATATGAACGTCAGGCCCTTGCTGAATTACTTTGAACCTCTGTTTACG<br>TGGTTGAAAGACCAAAATAAAAACTCCTTTGTTGGGTGGAGTACTGACTGGTCCCCCTATGCG |
| 278 | CAATCTACCATCGAAGAGCAGGCCAAAACATTCCTCGACAAGTTTAATGTCGAGGCTGAAGACC<br>TTTTCTACCAATCAAGTCTGGCTAGCTGGAATTACAATACAAACATTACAGAGGAGAACGTACA<br>AAACATGAATAACGCAGGGGACAAGTGGAGCGCATTCCTTAAGGAACAAAGTACCCTTGCGCAA<br>ATGTATCCGCTGCAAGAGATTCAACAACTGACGGTTAAGCTGCAACTTCAGGCCCTCCAACAAA<br>ATGGATCCTCAGTCTTGTCAGAAGACAAAAGCAAGCGACTGAACACCATCCTTAACACCATGTC<br>AACCATATATTCAACAGGTAAAGTTTGCAATCCGGATAACCCCCAAGAATGTTTGCTTCTTGAA<br>CCCGGTCTCAACGAAATTATGGCCAACAGTCTTGATTACAACGAGCGATTGTGGGCATGGGAAA<br>GTTGGAGGAGTGAGGTAGGCAAACAGTTGAGACCTCTTTATGAAGAGTACGTTGTCCTTAAAAA<br>TGAAATGGCTCGCGCGAATCATTATGAAGACTATGGTGACTACTGGAGGGGGGATTATGAGGTG<br>AACGGGGTGGACGGATACGATTACTCTAGGGGCCAGCTGATAGAGGATGTCGAGCACACCTTTG<br>AGGAGATTAAGCCGTTGTACGAACATTTGCACGCCTATGTCAGGGCTAAGCTCATGAACGCTTA<br>TCCGAGTTATATCTCCCCGATAGGATGCTTGCCTGCTCACTTGTTGGGCGATATGTGGGGACGC<br>TTTTGGACCAACTTGTATTCCCTTACGGTACCGTTCGGCCAGAAACCAAATATCGACGTGACAG<br>ACGCAATGGTGGATCAAGCATGGGATGCGCAACGAATCTTCAAGGAGGCAGAAAAATTTTTCGT<br>TTCAGTTGGACTCCCAAACATGACGCAGGGTTTCTGGGAGAACTCAATGTTGACAGATCCAGGT<br>AATGTGCAGAAAGCGGTTTGCCTCCCTACTGCATGGGATCTTGGTAAAGGGGACTTCCGCATAC<br>TCATGTGTACGAAAGTAACTATGGACGACTTTCTTACTGCGCACCACGAGATGGGGCACATACA<br>ATACGATATGGCGTACGCAGCTCAACCTTTCCTTCTGCGGAACGGGGCGAATGAAGGATTTCAC<br>GAGGCAGTGGGTGAGATTATGTCCCTGTCAGCTGCCACTCCGAAACATCTGAAAAGCATCGGCC<br>TGTTGAGCCCAGACTTCCAAGAAGATAATGAGACCGAAATAAACTTCCTTCTGAAGCAAGCACT<br>GACTATTGTAGGTACCTTGCCCTTTACATACATGCTGGAGAAGTGGAGGTGGATGGTATTTAAG<br>GGGGAGATACCGAAAGATCAATGGATGAAAAAGTGGTGGGAAATGAAAAGGGAGATCGTTGGCG<br>TAGTTGAACCAGTACCGCATGATGAGACGTACTGCGATCCGGCTAGTCTGTTCCATGTCTCTAA<br>TGATTACTCTTTCATCCGCTACTACACCCGCACGCTGTATCAATTCCAGTTCCAAGAAGCTCTC<br>TGTCAGGCTGCCAAGCACGAAGGACCGCTGCACAAATGCGACATTAGCAATTCTACAGAGGCGG<br>GTCAGAAGTTGTTCAATATGCTTAGACTGGGGAAGAGCGAACCGTGGACGCTCGCTTTGGAGAA<br>CGTTGTTGGAGCTAAGAATATGAACGTCAGGCCCTTGCTGAATTACTTTGAACCTCTGTTTACG<br>TGGTTGAAAGACCAAAATAAAAACTCCTTTGTTGGGTGGAGTACTGACTGGTCCCCCTATGCG |
| 292 | CAATCTACCATCGAAGAGCAGGCCAAAACATTCCTCGACAAGTTTAATGTCGAGGCTGAAGACC<br>TTTTCTACCAATCAAGTCTGGCTAGCTGGAATTACAATACAAACATTACAGAGGAGAACGTACA |

| | |
|---|---|
| | AAACATGAATAACGCAGGGGACAAGTGGAGCGCATTCCTTAAGGAACAAAGTACCCTTGCGCAA<br>ATGTATCCGCTGCAAGAGATTCAACAACTGACGGTTAAGCTGCAACTTCAGGCCCTCCAACAAA<br>ATGGATCCTCAGTCTTGTCAGAAGACAAAAGCAAGCGACTGAACACCATCCTTAACACCATGTC<br>AACCATATATTCAACAGGTAAAGTTTGCAATCCGGATAACCCCCAAGAATGTTTGCTTCTTGAA<br>CCCGGTCTCAACGAAATTATGGCCAACAGTCTTGATTACAACGAGCGATTGTGGGCATGGGAAA<br>GTTGGAGGAGTGAGGTAGGCAAACAGTTGAGACCTCTTTATGAAGAGTACGTTGTCCTTAAAAA<br>TGAAATGGCTCGCGCGAATCATTATGAAGACTATGGTGACTACTGGAGGGGGGATTATGAGGTG<br>AACGGGGTGGACGGATACGATTACTCTAGGGGCCAGCTGATAGAGGATGTCGAGCACACCTTTG<br>AGGAGATTAAGCCGTTGTACGAACATTTGCACGCCTATGTCAGGGCTAAGCTCATGAACGCTTA<br>TCCGAGTTATATCTCCCCGATAGGATGCTTGCCTGCTCACTTGTTGGGCGATATGTGGGGACGC<br>TTTTGGACCAACTTGTATTCCCTTACGGTACCGTTCGGCCAGAAACCAAATATCGACGTGACAG<br>ACGCAATGGTGGATCAAGCATGGGATGCGCAACGAATCTTCAAGGAGGCAGAAAAATTTTTCGT<br>TTCAGTTGGACTCCCAAACATGACGCAGGGTTTCTGGGAGAACTCAATGTTGACAGATCCAGGT<br>AATGTGCAGAAAGCGGTTTGCCACCCTACTGCATGGGATCTTGGTAAAGGGGACTTCCGCATAC<br>TCATGTGTACGAAAGTAACTATGGACGACTTTCTTACTGCGCACCACGAGATGGGGCACATACA<br>ATACGATATGGCGTACGCAGCTCAACCTTTCCTTCTGCGGAACGGGGCGAATGAAGGATTTCAC<br>GAGGCAGTGGGTGAGATTATGTCCCTGTCAGCTGCCACTCCGAAACATCTGAAAAGCATCGGCC<br>TGTTGAGCCCAGACTTCCAAGAAGATAATGAGACCGAAATAAACTTCCTTCTGAAGCAAGCACT<br>GACTATTGTAGGTACCTTGCCCTTTACATACATGCTGGAGAAGTGGAGGTGGATGGTATTTAAG<br>GGGGAGATACCGAAAGATCAATGGATGAAAAAGTGGTGGGAAATGAAAAGGGAGATCGTTGGCG<br>TAGTTGAACCAGTACCGCATGATGAGACGTACTGCGATCCGGCTAGTCTGTTCCATGTCTCTAA<br>TGATTACTCTTTCATCCGCTACTACACCCGCACGCTGTATCAATTCCAGTTCCAAGAAGCTCTC<br>TGTCAGGCTGCCAAGCACGAAGGACCGCTGCACAAATGCGACATTAGCAATTCTACAGAGGCGG<br>GTCAGAAGTTGTTCAATATGCTTAGACTGGGGAAGAGCGAACCGTGGACGCTCGCTTTGGAGAA<br>CGTTGTTGGAGCTAAGAATATGAACGTCAGGCCCTTGCTGAATTACTTTGAACCTCTGTTTACG<br>TGGTTGAAAGACCAAAATAAAAACTCCTTTGTTGGGTGGAGTACTGACTGGTCCCCCTATGCGG<br>ACCAAAGCATCAAAGTGAGGATAAGCCTAAAATCAGCTCTTGGAGATAAAGCATATGAATGGAA<br>CGACAATGAAATGTACCTGTTCCGATCATCTGTTGCATATGCTATGAGGCAGTACTTTTTAAAA<br>GTAAAAAATCAGATGATTCTTTTTGGGGAGGAGGATGTGCGAGTGGCTAATTTGAAACCAAGAA<br>TCTCCTTTAATTTCTTTGTCACTGCACCTAAAAATGTGTCTGATATCATTCCTAGAACTGAAGT<br>TGAAAAGGCCATCAGGATGTCCCGGAGCCGTATCAATGATGCTTTCCGTCTGAATGACAACAGC<br>CTAGAGTTTCTGGGGATACAGCCAACACTTGGACCTCCTAACCAGCCCCCTGTTTCC |
| 310 | CAATCTACCATCGAAGAGCAGGTTAAATATTTCCTCGACAAGTTTAATGCTGAGGCTGAAGACC<br>TTGATTACCAATCAAGTCTGGCTAGCTGGAATTACAATACAAACATTACAGAGGAGAACGTACA<br>AAACATGAATAACGCAGGGGACAAGTGGAGCGCATTCCTTAAGGAACAAAGTACCCTTGCGCAA<br>ATGTATCCGCTGCAAGAGATTCAAAACCTGACGGTTAAGCTGCAACTTCAGGCCCTCCAACAAA<br>ATGGATCCTCAGTCTTGTCAGAAGACAAAAGCAAGCGACTGAACACCATCCTTAACACCATGTC<br>AACCATATATTCAACAGGTAAAGTTTGCAATCCGGATAACCCCCAAGAATGTTTGCTTCTTGAA<br>CCCGGTCTCAACGAAATTATGGCCAACAGTCTTGATTACAACGAGCGATTGTGGGCATGGGAAA<br>GTTGGAGGAGTGAGGTAGGCAAACAGTTGAGACCTCTTTATGAAGAGTACGTTGTCCTTAAAAA<br>TGAAATGGCTCGCGCGAATCATTATGAAGACTATGGTGACTACTGGAGGGGGGATTATGAGGTG<br>AACGGGGTGGACGGATACGATTACTCTAGGGGCCAGCTGATAGAGGATGTCGAGCACACCTTTG<br>AGGAGATTAAGCCGTTGTACGAACATTTGCACGCCTATGTCAGGGCTAAGCTCATGAACGCTTA<br>TCCGAGTTATATCTCCCCGATAGGATGCTTGCCTGCTCACTTGTTGGGCGATATGTGGGGACGC<br>TTTTGGACCAACTTGTATTCCCTTACGGTACCGTTCGGCCAGAAACCAAATATCGACGTGACAG<br>ACGCAATGGTGGATCAAGCATGGGATGCGCAACGAATCTTCAAGGAGGCAGAAAAATTTTTCGT<br>TTCAGTTGGACTCCCAAACATGACGCAGGGTTTCTGGGAGAACTCAATGTTGACAGATCCAGGT |

| | |
|---|---|
| | AATGTGCAGAAAGCGGTTTGCCTCCCTACTGCATGGGATCTTGGTAAAGGGGACTTCCGCATAC<br>TCATGTGTACGAAAGTAACTATGGACGACTTTCTTACTGCGCACCACGAGATGGGGCACATACA<br>ATACGATATGGCGTACGCAGCTCAACCTTTCCTTCTGCGGAACGGGGCGAATGAAGGATTTCAC<br>GAGGCAGTGGGTGAGATTATGTCCCTGTCAGCTGCCACTCCGAAACATCTGAAAAGCATCGGCC<br>TGTTGAGCCCAGACTTCCAAGAAGATAATGAGACCGAAATAAACTTCCTTCTGAAGCAAGCACT<br>GACTATTGTAGGTACCTTGCCCTTTACATACATGCTGGAGAAGTGGAGGTGGATGGTATTTAAG<br>GGGGAGATACCGAAAGATCAATGGATGAAAAAGTGGTGGGAAATGAAAAGGGAGATCGTTGGCG<br>TAGTTGAACCAGTACCGCATGATGAGACGTACTGCGATCCGGCTAGTCTGTTCCATGTCTCTAA<br>TGATTACTCTTTCATCCGCTACTACACCCGCACGCTGTATCAATTCCAGTTCCAAGAAGCTCTC<br>TGTCAGGCTGCCAAGCACGAAGGACCGCTGCACAAATGCGACATTAGCAATTCTACAGAGGCGG<br>GTCAGAAGTTGTTCAATATGCTTAGACTGGGGAAGAGCGAACCGTGGACGCTCGCTTTGGAGAA<br>CGTTGTTGGAGCTAAGAATATGAACGTCAGGCCCTTGCTGAATTACTTTGAACCTCTGTTTACG<br>TGGTTGAAAGACCAAAATAAAAACTCCTTTGTTGGGTGGAGTACTGACTGGTCCCCCTATGCGG<br>ACCAAAGCATCAAAGTGAGGATAAGCCTAAAATCAGCTCTTGGAGATAAAGCATATGAATGGAA<br>CGACAATGAAATGTACCTGTTCCGATCATCTGTTGCATATGCTATGAGGCAGTACTTTTTAAAA<br>GTAAAAAATCAGATGATTCTTTTTGGGGAGGAGGATGTGCGAGTGGCTAATTTGAAACCAAGAA<br>TCTCCTTTAATTTCTTTGTCACTGCACCTAAAAATGTGTCTGATATCATTCCTAGAACTGAAGT<br>TGAAAAGGCCATCAGGATGTCCCGGAGCCGTATCAATGATGCTTTCCGTCTGAATGACAACAGC<br>CTAGAGTTTCTGGGGATACAGCCAACACTTGGACCTCCTAACCAGCCCCCTGTTTCC |
| 311 | CAATCTACCATCGAAGAGCAGGCCAAAACATTCCTCGACTATTTTAATCACGAGGCTGAAGACC<br>TTTTCTACCAATCAAGTCTGGCTAGCTGGAATTACAATACAAACATTACAGAGGAGAACGTACA<br>AAACATGAATAACGCAGGGGACAAGGTTAGCGCATTCCTTAAGGAACAAAGTACCACTGCGCAA<br>ATGTATCCGCTGCAAGAGATTCAAAACCCAACGGTTAAGCTGCAACTTCAGGCCCTCCAACAAA<br>ATGGATCCTCAGTCTTGTCAGAAGACAAAAGCAAGCGACTGAACACCATCCTTAACACCATGTC<br>AACCATATATTCAACAGGTAAAGTTTGCAATCCGGATAACCCCCAAGAATGTTTGCTTCTTGAA<br>CCCGGTCTCAACGAAATTATGGCCAACAGTCTTGATTACAACGAGCGATTGTGGGCATGGGAAA<br>GTTGGAGGAGTGAGGTAGGCAAACAGTTGAGACCTCTTTATGAAGAGTACGTTGTCCTTAAAAA<br>TGAAATGGCTCGCGCGAATCATTATGAAGACTATGGTGACTACTGGAGGGGGGATTATGAGGTG<br>AACGGGGTGGACGGATACGATTACTCTAGGGGCCAGCTGATAGAGGATGTCGAGCACACCTTTG<br>AGGAGATTAAGCCGTTGTACGAACATTTGCACGCCTATGTCAGGGCTAAGCTCATGAACGCTTA<br>TCCGAGTTATATCTCCCCGATAGGATGCTTGCCTGCTCACTTGTTGGGCGATATGTGGGGACGC<br>TTTTGGACCAACTTGTATTCCCTTACGGTACCGTTCGGCCAGAAACCAAATATCGACGTGACAG<br>ACGCAATGGTGGATCAAGCATGGGATGCGCAACGAATCTTCAAGGAGGCAGAAAAATTTTTCGT<br>TTCAGTTGGACTCCCAAACATGACGCAGGGTTTCTGGGAGAACTCAATGTTGACAGATCCAGGT<br>AATGTGCAGAAAGCGGTTTGCCTCCCTACTGCATGGGATCTTGGTAAAGGGGACTTCCGCATAC<br>TCATGTGTACGAAAGTAACTATGGACGACTTTCTTACTGCGCACCACGAGATGGGGCACATACA<br>ATACGATATGGCGTACGCAGCTCAACCTTTCCTTCTGCGGAACGGGGCGAATGAAGGATTTCAC<br>GAGGCAGTGGGTGAGATTATGTCCCTGTCAGCTGCCACTCCGAAACATCTGAAAAGCATCGGCC<br>TGTTGAGCCCAGACTTCCAAGAAGATAATGAGACCGAAATAAACTTCCTTCTGAAGCAAGCACT<br>GACTATTGTAGGTACCTTGCCCTTTACATACATGCTGGAGAAGTGGAGGTGGATGGTATTTAAG<br>GGGGAGATACCGAAAGATCAATGGATGAAAAAGTGGTGGGAAATGAAAAGGGAGATCGTTGGCG<br>TAGTTGAACCAGTACCGCATGATGAGACGTACTGCGATCCGGCTAGTCTGTTCCATGTCTCTAA<br>TGATTACTCTTTCATCCGCTACTACACCCGCACGCTGTATCAATTCCAGTTCCAAGAAGCTCTC<br>TGTCAGGCTGCCAAGCACGAAGGACCGCTGCACAAATGCGACATTAGCAATTCTACAGAGGCGG<br>GTCAGAAGTTGTTCAATATGCTTAGACTGGGGAAGAGCGAACCGTGGACGCTCGCTTTGGAGAA<br>CGTTGTTGGAGCTAAGAATATGAACGTCAGGCCCTTGCTGAATTACTTTGAACCTCTGTTTACG<br>TGGTTGAAAGACCAAAATAAAAACTCCTTTGTTGGGTGGAGTACTGACTGGTCCCCCTATGCGG |

| | |
|---|---|
| | ACCAAAGCATCAAAGTGAGGATAAGCCTAAAATCAGCTCTTGGAGATAAAGCATATGAATGGAA<br>CGACAATGAAATGTACCTGTTCCGATCATCTGTTGCATATGCTATGAGGCAGTACTTTTTAAAA<br>GTAAAAAATCAGATGATTCTTTTTGGGGAGGAGGATGTGCGAGTGGCTAATTTGAAACCAAGAA<br>TCTCCTTTAATTTCTTTGTCACTGCACCTAAAAATGTGTCTGATATCATTCCTAGAACTGAAGT<br>TGAAAAGGCCATCAGGATGTCCCGGAGCCGTATCAATGATGCTTTCCGTCTGAATGACAACAGC<br>CTAGAGTTTCTGGGGATACAGCCAACACTTGGACCTCCTAACCAGCCCCCTGTTTCC |
| 312 | CAATCTACCATCGAAGAGCAGGCCAAATATTTCCTCGACAAGTTTAATGCTGAGGCTGAAGACC<br>TTTTCTACCAATCAAGTCTGGCTAGCTGGAATTACAATACAAACATTACAGAGGAGAACGTACA<br>AAACATGAATAACGCAGGGGACAAGTGGAGCGCATTCCTTAAGGAACAAAGTACCCTTGCGCAA<br>ATGTATCCGCTGCAAGAGATTCAACAACTGACGGTTAAGCTGCAACTTCAGGCCCTCCAACAAA<br>ATGGATCCTCAGTCTTGTCAGAAGACAAAAGCAAGCGACTGAACACCATCCTTAACACCATGTC<br>AACCATATATTCAACAGGTAAAGTTTGCAATCCGGATAACCCCCAAGAATGTTTGCTTCTTGAA<br>CCCGGTCTCAACGAAATTATGGCCAACAGTCTTGATTACAACGAGCGATTGTGGGCATGGGAAA<br>GTTGGAGGAGTGAGGTAGGCAAACAGTTGAGACCTCTTTATGAAGAGTACGTTGTCCTTAAAAA<br>TGAAATGGCTCGCGCGAATCATTATGAAGACTATGGTGACTACTGGAGGGGGGATTATGAGGTG<br>AACGGGGTGGACGGATACGATTACTCTAGGGGCCAGCTGATAGAGGATGTCGAGCACACCTTTG<br>AGGAGATTAAGCCGTTGTACGAACATTTGCACGCCTATGTCAGGGCTAAGCTCATGAACGCTTA<br>TCCGAGTTATATCTCCCCGATAGGATGCTTGCCTGCTCACTTGTTGGGCGATATGTGGGGACGC<br>TTTTGGACCAACTTGTATTCCCTTACGGTACCGTTCGGCCAGAAACCAAATATCGACGTGACAG<br>ACGCAATGGTGGATCAAGCATGGGATGCGCAACGAATCTTCAAGGAGGCAGAAAAATTTTTCGT<br>TTCAGTTGGACTCCCAAACATGACGCAGGGTTTCTGGGAGAACTCAATGTTGACAGATCCAGGT<br>AATGTGCAGAAAGCGGTTTGCCTCCCTACTGCATGGGATCTTGGTAAAGGGGACTTCCGCATAC<br>TCATGTGTACGAAAGTAACTATGGACGACTTTCTTACTGCGCACCACGAGATGGGGCACATACA<br>ATACGATATGGCGTACGCAGCTCAACCTTTCCTTCTGCGGAACGGGGCGAATGAAGGATTTCAC<br>GAGGCAGTGGGTGAGATTATGTCCCTGTCAGCTGCCACTCCGAAACATCTGAAAAGCATCGGCC<br>TGTTGAGCCCAGACTTCCAAGAAGATAATGAGACCGAAATAAACTTCCTTCTGAAGCAAGCACT<br>GACTATTGTAGGTACCTTGCCCTTTACATACATGCTGGAGAAGTGGAGGTGGATGGTATTTAAG<br>GGGGAGATACCGAAAGATCAATGGATGAAAAAGTGGTGGGAAATGAAAAGGGAGATCGTTGGCG<br>TAGTTGAACCAGTACCGCATGATGAGACGTACTGCGATCCGGCTAGTCTGTTCCATGTCTCTAA<br>TGATTACTCTTTCATCCGCTACTACACCCGCACGCTGTATCAATTCCAGTTCCAAGAAGCTCTC<br>TGTCAGGCTGCCAAGCACGAAGGACCGCTGCACAAATGCGACATTAGCAATTCTACAGAGGCGG<br>GTCAGAAGTTGTTCAATATGCTTAGACTGGGGAAGAGCGAACCGTGGACGCTCGCTTTGGAGAA<br>CGTTGTTGGAGCTAAGAATATGAACGTCAGGCCCTTGCTGAATTACTTTGAACCTCTGTTTACG<br>TGGTTGAAAGACCAAAATAAAAACTCCTTTGTTGGGTGGAGTACTGACTGGTCCCCCTATGCGG<br>ACCAAAGCATCAAAGTGAGGATAAGCCTAAAATCAGCTCTTGGAGATAAAGCATATGAATGGAA<br>CGACAATGAAATGTACCTGTTCCGATCATCTGTTGCATATGCTATGAGGCAGTACTTTTTAAAA<br>GTAAAAAATCAGATGATTCTTTTTGGGGAGGAGGATGTGCGAGTGGCTAATTTGAAACCAAGAA<br>TCTCCTTTAATTTCTTTGTCACTGCACCTAAAAATGTGTCTGATATCATTCCTAGAACTGAAGT<br>TGAAAAGGCCATCAGGATGTCCCGGAGCCGTATCAATGATGCTTTCCGTCTGAATGACAACAGC<br>CTAGAGTTTCTGGGGATACAGCCAACACTTGGACCTCCTAACCAGCCCCCTGTTTCC |
| 313 | CAATCTACCATCGAAGAGCAGGCCAAAACATTCCTCGACTTCTTTGATAGCCAGGCTGAAGACC<br>TTTTCTACCAATCAAGTCTGGCTAGCTGGAATTACAATACAAACATTACAGAGGAGAACGTACA<br>AAACATGAATAACGCAGGGGACAAGTGGAGCGCATTCCTTAAGGAACAAAGTACCCTTGCGCAA<br>ATGTATCCGCTGCAAGAGATTCAAAACCTGACGGTTAAGCTGCAACTTCAGGCCCTCCAACAAA<br>ATGGATCCTCAGTCTTGTCAGAAGACAAAAGCAAGCGACTGAACACCATCCTTAACACCATGTC<br>AACCATATATTCAACAGGTAAAGTTTGCAATCCGGATAACCCCCAAGAATGTTTGCTTCTTGAA<br>CCCGGTCTCAACGAAATTATGGCCAACAGTCTTGATTACAACGAGCGATTGTGGGCATGGGAAA |

| | |
|---|---|
| | GTTGGAGGAGTGAGGTAGGCAAACAGTTGAGACCTCTTTATGAAGAGTACGTTGTCCTTAAAAA<br>TGAAATGGCTCGCGCGAATCATTATGAAGACTATGGTGACTACTGGAGGGGGGATTATGAGGTG<br>AACGGGGTGGACGGATACGATTACTCTAGGGGCCAGCTGATAGAGGATGTCGAGCACACCTTTG<br>AGGAGATTAAGCCGTTGTACGAACATTTGCACGCCTATGTCAGGGCTAAGCTCATGAACGCTTA<br>TCCGAGTTATATCTCCCCGATAGGATGCTTGCCTGCTCACTTGTTGGGCGATATGTGGGGACGC<br>TTTTGGACCAACTTGTATTCCCTTACGGTACCGTTCGGCCAGAAACCAAATATCGACGTGACAG<br>ACGCAATGGTGGATCAAGCATGGGATGCGCAACGAATCTTCAAGGAGGCAGAAAAATTTTTCGT<br>TTCAGTTGGACTCCCAAACATGACGCAGGGTTTCTGGGAGAACTCAATGTTGACAGATCCAGGT<br>AATGTGCAGAAAGCGGTTTGCCTCCCTACTGCATGGGATCTTGGTAAAGGGGACTTCCGCATAC<br>TCATGTGTACGAAAGTAACTATGGACGACTTTCTTACTGCGCACCACGAGATGGGGCACATACA<br>ATACGATATGGCGTACGCAGCTCAACCTTTCCTTCTGCGGAACGGGGCGAATGAAGGATTTCAC<br>GAGGCAGTGGGTGAGATTATGTCCCTGTCAGCTGCCACTCCGAAACATCTGAAAAGCATCGGCC<br>TGTTGAGCCCAGACTTCCAAGAAGATAATGAGACCGAAATAAACTTCCTTCTGAAGCAAGCACT<br>GACTATTGTAGGTACCTTGCCCTTTACATACATGCTGGAGAAGTGGAGGTGGATGGTATTTAAG<br>GGGGAGATACCGAAAGATCAATGGATGAAAAAGTGGTGGGAAATGAAAAGGGAGATCGTTGGCG<br>TAGTTGAACCAGTACCGCATGATGAGACGTACTGCGATCCGGCTAGTCTGTTCCATGTCTCTAA<br>TGATTACTCTTTCATCCGCTACTACACCCGCACGCTGTATCAATTCCAGTTCCAAGAAGCTCTC<br>TGTCAGGCTGCCAAGCACGAAGGACCGCTGCACAAATGCGACATTAGCAATTCTACAGAGGCGG<br>GTCAGAAGTTGTTCAATATGCTTAGACTGGGGAAGAGCGAACCGTGGACGCTCGCTTTGGAGAA<br>CGTTGTTGGAGCTAAGAATATGAACGTCAGGCCCTTGCTGAATTACTTTGAACCTCTGTTTACG<br>TGGTTGAAAGACCAAAATAAAAACTCCTTTGTTGGGTGGAGTACTGACTGGTCCCCCTATGCGG<br>ACCAAAGCATCAAAGTGAGGATAAGCCTAAAATCAGCTCTTGGAGATAAAGCATATGAATGGAA<br>CGACAATGAAATGTACCTGTTCCGATCATCTGTTGCATATGCTATGAGGCAGTACTTTTTAAAA<br>GTAAAAAATCAGATGATTCTTTTTGGGGAGGAGGATGTGCGAGTGGCTAATTTGAAACCAAGAA<br>TCTCCTTTAATTTCTTTGTCACTGCACCTAAAAATGTGTCTGATATCATTCCTAGAACTGAAGT<br>TGAAAAGGCCATCAGGATGTCCCGGAGCCGTATCAATGATGCTTTCCGTCTGAATGACAACAGC<br>CTAGAGTTTCTGGGGATACAGCCAACACTTGGACCTCCTAACCAGCCCCCTGTTTCC |
| 353 | CAATCTACCATCGAAGAGCAGGCCAAAGCATTCCTCGACTTCTTTGATAGCCAGGCTGAAGACC<br>TTTTCTACCAATCAAGTCTGGCTAGCTGGAATTACAATACAAACATTACAGAGGAGAACGTACA<br>AGACATGAATAACGCAGGGGACAGGTGGAGCGCATTCCTTAAGGAACAAAGTACCCCTGCGCAA<br>ATGTATCCGCTGCAAGAGATTCAAAACCTGACGGTTAAGCTGCAACTTCAGGCCCTCCAACAAA<br>ATGGATCCTCAGTCTTGTCAGAAGACAAAAGCAAGCGACTGAACACCATCCTTAACACCATGTC<br>AACCATATATTCAACAGGTAAAGTTTGCAATCCGGATAACCCCCAAGAATGTTTGCTTCTTGAA<br>CCCGGTCTCAACGAAATTATGGCCAACAGTCTTGATTACAACGAGCGATTGTGGGCATGGGAAA<br>GTTGGAGGAGTGAGGTAGGCAAACAGTTGAGACCTCTTTATGAAGAGTACGTTGTCCTTAAAAA<br>TGAAATGGCTCGCGCGAATCATTATGAAGACTATGGTGACTACTGGAGGGGGGATTATGAGGTG<br>AACGGGGTGGACGGATACGATTACTCTAGGGGCCAGCTGATAGAGGATGTCGAGCACACCTTTG<br>AGGAGATTAAGCCGTTGTACGAACATTTGCACGCCTATGTCAGGGCTAAGCTCATGAACGCTTA<br>TCCGAGTTATATCTCCCCGATAGGATGCTTGCCTGCTCACTTGTTGGGCGATATGTGGGGACGC<br>TTTTGGACCAACTTGTATTCCCTTACGGTACCGTTCGGCCAGAAACCAAATATCGACGTGACAG<br>ACGCAATGGTGGATCAAGCATGGGATGCGCAACGAATCTTCAAGGAGGCAGAAAAATTTTTCGT<br>TTCAGTTGGACTCCCAAACATGACGCAGGGTTTCTGGGAGAACTCAATGTTGACAGATCCAGGT<br>AATGTGCAGAAAGCGGTTTGCCTCCCTACTGCATGGGATCTTGGTAAAGGGGACTTCCGCATAC<br>TCATGTGTACGAAAGTAACTATGGACGACTTTCTTACTGCGCACCACGAGATGGGGCACATACA<br>ATACGATATGGCGTACGCAGCTCAACCTTTCCTTCTGCGGAACGGGGCGAATGAAGGATTTCAC<br>GAGGCAGTGGGTGAGATTATGTCCCTGTCAGCTGCCACTCCGAAACATCTGAAAAGCATCGGCC<br>TGTTGAGCCCAGACTTCCAAGAAGATAATGAGACCGAAATAAACTTCCTTCTGAAGCAAGCACT |

| | |
|---|---|
| | GACTATTGTAGGTACCTTGCCCTTTACATACATGCTGGAGAAGTGGAGGTGGATGGTATTTAAG<br>GGGGAGATACCGAAAGATCAATGGATGAAAAAGTGGTGGGAAATGAAAAGGGAGATCGTTGGCG<br>TAGTTGAACCAGTACCGCATGATGAGACGTACTGCGATCCGGCTAGTCTGTTCCATGTCTCTAA<br>TGATTACTCTTTCATCCGCTACTACACCCGCACGCTGTATCAATTCCAGTTCCAAGAAGCTCTC<br>TGTCAGGCTGCCAAGCACGAAGGACCGCTGCACAAATGCGACATTAGCAATTCTACAGAGGCGG<br>GTCAGAAGTTGTTCAATATGCTTAGACTGGGGAAGAGCGAACCGTGGACGCTCGCTTTGGAGAA<br>CGTTGTTGGAGCTAAGAATATGAACGTCAGGCCCTTGCTGAATTACTTTGAACCTCTGTTTACG<br>TGGTTGAAAGACCAAAATAAAAACTCCTTTGTTGGGTGGAGTACTGACTGGTCCCCCTATGCGG<br>ACCAAAGCATCAAAGTGAGGATAAGCCTAAAATCAGCTCTTGGAGATAAAGCATATGAATGGAA<br>CGACAATGAAATGTACCTGTTCCGATCATCTGTTGCATATGCTATGAGGCAGTACTTTTTAAAA<br>GTAAAAAATCAGATGATTCTTTTTGGGGAGGAGGATGTGCGAGTGGCTAATTTGAAACCAAGAA<br>TCTCCTTTAATTTCTTTGTCACTGCACCTAAAAATGTGTCTGATATCATTCCTAGAACTGAAGT<br>TGAAAAGGCCATCAGGATGTCCCGGAGCCGTATCAATGATGCTTTCCGTCTGAATGACAACAGC<br>CTAGAGTTTCTGGGGATACAGCCAACACTTGGACCTCCTAACCAGCCCCCTGTTTCC |
| 354 | CAATCTACCATCGAAGAGCAGGCCAAAACATTCCTCGACTTCTTTGATGCCCAGGCTGAAGACC<br>TTTTCTACCAATCAAGTCTGGCTAGCTGGGATTACAGTACAAGCATTACAGAGGGGAACGTGCA<br>AAACATGAATGACGCAGGGGACAAGTGGAGCGCATTCCTTAAGGAGCAAAGTACCCTTGCGCAA<br>ATGTATCCGCTGCAAGAGATTCAAAACCTGACGGTTAAGCTGCAACTTCAGGCCCTCCAGCAAA<br>ATGGATCCTCAGTCTTGTCAGAAGACAAAAGCAAGCGACTGAACACCATCCTTAACACCATGTC<br>AACCATATATTCAACAGGTAAAGTTTGCAATCCGGATAACCCCCAAGAATGTTTGCTTCTTGAA<br>CCCGGTCTCAACGAAATTATGGCCAACAGTCTTGATTACAACGAGCGATTGTGGGCATGGGAAA<br>GTTGGAGGAGTGAGGTAGGCAAACAGTTGAGACCTCTTTATGAAGAGTACGTTGTCCTTAAAAA<br>TGAAATGGCTCGCGCGAATCATTATGAAGACTATGGTGACTACTGGAGGGGGGATTATGAGGTG<br>AACGGGGTGGACGGATACGATTACTCTAGGGGCCAGCTGATAGAGGATGTCGAGCACACCTTTG<br>AGGAGATTAAGCCGTTGTACGAACATTTGCACGCCTATGTCAGGGCTAAGCTCATGAACGCTTA<br>TCCGAGTTATATCTCCCCGATAGGATGCTTGCCTGCTCACTTGTTGGGCGATATGTGGGGACGC<br>TTTTGGACCAACTTGTATTCCCTTACGGTACCGTTCGGCCAGAAACCAAATATCGACGTGACAG<br>ACGCAATGGTGGATCAAGCATGGGATGCGCAACGAATCTTCAAGGAGGCAGAAAAATTTTTCGT<br>TTCAGTTGGACTCCCAAACATGACGCAGGGTTTCTGGGAGAACTCAATGTTGACAGATCCAGGT<br>AATGTGCAGAAAGCGGTTTGCCTCCCTACTGCATGGGATCTTGGTAAAGGGGACTTCCGCATAC<br>TCATGTGTACGAAAGTAACTATGGACGACTTTCTTACTGCGCACCACGAGATGGGGCACATACA<br>ATACGATATGGCGTACGCAGCTCAACCTTTCCTTCTGCGGAACGGGGCGAATGAAGGATTTCAC<br>GAGGCAGTGGGTGAGATTATGTCCCTGTCAGCTGCCACTCCGAAACATCTGAAAAGCATCGGCC<br>TGTTGAGCCCAGACTTCCAAGAAGATAATGAGACCGAAATAAACTTCCTTCTGAAGCAAGCACT<br>GACTATTGTAGGTACCTTGCCCTTTACATACATGCTGGAGAAGTGGAGGTGGATGGTATTTAAG<br>GGGGAGATACCGAAAGATCAATGGATGAAAAAGTGGTGGGAAATGAAAAGGGAGATCGTTGGCG<br>TAGTTGAACCAGTACCGCATGATGAGACGTACTGCGATCCGGCTAGTCTGTTCCATGTCTCTAA<br>TGATTACTCTTTCATCCGCTACTACACCCGCACGCTGTATCAATTCCAGTTCCAAGAAGCTCTC<br>TGTCAGGCTGCCAAGCACGAAGGACCGCTGCACAAATGCGACATTAGCAATTCTACAGAGGCGG<br>GTCAGAAGTTGTTCAATATGCTTAGACTGGGGAAGAGCGAACCGTGGACGCTCGCTTTGGAGAA<br>CGTTGTTGGAGCTAAGAATATGAACGTCAGGCCCTTGCTGAATTACTTTGAACCTCTGTTTACG<br>TGGTTGAAAGACCAAAATAAAAACTCCTTTGTTGGGTGGAGTACTGACTGGTCCCCCTATGCGG<br>ACCAAAGCATCAAAGTGAGGATAAGCCTAAAATCAGCTCTTGGAGATAAAGCATATGAATGGAA<br>CGACAATGAAATGTACCTGTTCCGATCATCTGTTGCATATGCTATGAGGCAGTACTTTTTAAAA<br>GTAAAAAATCAGATGATTCTTTTTGGGGAGGAGGATGTGCGAGTGGCTAATTTGAAACCAAGAA<br>TCTCCTTTAATTTCTTTGTCACTGCACCTAAAAATGTGTCTGATATCATTCCTAGAACTGAAGT |

| | |
|---|---|
| | TGAAAAGGCCATCAGGATGTCCCGGAGCCGTATCAATGATGCTTTCCGTCTGAATGACAACAGC<br>CTAGAGTTTCTGGGGATACAGCCAACACTTGGACCTCCTAACCAGCCCCCTGTTTCC |
| 355 | CAACCAACCATCGAAGAGCAGGCCAAAACATTCCTCGACAAGTTTAATCACGAGGCTGAAGACC<br>TTTTCTACTTGTCAAGTCTGGCTAGCTGGAATTACAATACAAACATTACAGAGGAGAACGTACA<br>AAACATGAATAACGCAGGGGACAAGTGGAGCGCATTCCTTAAGGAACAAAGTACCACTGCGCAA<br>ATGTATCCGCTGCAAGAGATTCAACAGCTGACGGTTAAGCTGCAACTTCAGGCCCTCCAACAAA<br>ATGGATCCTCAGTCTTGTCAGAAGACAAAAGCAAGCGACTGAACACCATCCTTAACACCATGTC<br>AACCATATATTCAACAGGTAAAGTTTGCAATCCGGATAACCCCCAAGAATGTTTGCTTCTTGAA<br>CCCGGTCTCAACGAAATTATGGCCAACAGTCTTGATTACAACGAGCGATTGTGGGCATGGGAAA<br>GTTGGAGGAGTGAGGTAGGCAAACAGTTGAGACCTCTTTATGAAGAGTACGTTGTCCTTAAAAA<br>TGAAATGGCTCGCGCGAATCATTATGAAGACTATGGTGACTACTGGAGGGGGGATTATGAGGTG<br>AACGGGGTGGACGGATACGATTACTCTAGGGGCCAGCTGATAGAGGATGTCGAGCACACCTTTG<br>AGGAGATTAAGCCGTTGTACGAACATTTGCACGCCTATGTCAGGGCTAAGCTCATGAACGCTTA<br>TCCGAGTTATATCTCCCCGATAGGATGCTTGCCTGCTCACTTGTTGGGCGATATGTGGGGACGC<br>TTTTGGACCAACTTGTATTCCCTTACGGTACCGTTCGGCCAGAAACCAAATATCGACGTGACAG<br>ACGCAATGGTGGATCAAGCATGGGATGCGCAACGAATCTTCAAGGAGGCAGAAAAATTTTTCGT<br>TTCAGTTGGACTCCCAAACATGACGCAGGGTTTCTGGGAGAACTCAATGTTGACAGATCCAGGT<br>AATGTGCAGAAAGCGGTTTGCCTCCCTACTGCATGGGATCTTGGTAAAGGGGACTTCCGCATAC<br>TCATGTGTACGAAAGTAACTATGGACGACTTTCTTACTGCGCACCACGAGATGGGGCACATACA<br>ATACGATATGGCGTACGCAGCTCAACCTTTCCTTCTGCGGAACGGGGCGAATGAAGGATTTCAC<br>GAGGCAGTGGGTGAGATTATGTCCCTGTCAGCTGCCACTCCGAAACATCTGAAAAGCATCGGCC<br>TGTTGAGCCCAGACTTCCAAGAAGATAATGAGACCGAAATAAACTTCCTTCTGAAGCAAGCACT<br>GACTATTGTAGGTACCTTGCCCTTTACATACATGCTGGAGAAGTGGAGGTGGATGGTATTTAAG<br>GGGGAGATACCGAAAGATCAATGGATGAAAAAGTGGTGGGAAATGAAAAGGGAGATCGTTGGCG<br>TAGTTGAACCAGTACCGCATGATGAGACGTACTGCGATCCGGCTAGTCTGTTCCATGTCTCTAA<br>TGATTACTCTTTCATCCGCTACTACACCCGCACGCTGTATCAATTCCAGTTCCAAGAAGCTCTC<br>TGTCAGGCTGCCAAGCACGAAGGACCGCTGCACAAATGCGACATTAGCAATTCTACAGAGGCGG<br>GTCAGAAGTTGTTCAATATGCTTAGACTGGGGAAGAGCGAACCGTGGACGCTCGCTTTGGAGAA<br>CGTTGTTGGAGCTAAGAATATGAACGTCAGGCCCTTGCTGAATTACTTTGAACCTCTGTTTACG<br>TGGTTGAAAGACCAAAATAAAAACTCCTTTGTTGGGTGGAGTACTGACTGGTCCCCCTATGCGG<br>ACCAAAGCATCAAAGTGAGGATAAGCCTAAAATCAGCTCTTGGAGATAAAGCATATGAATGGAA<br>CGACAATGAAATGTACCTGTTCCGATCATCTGTTGCATATGCTATGAGGCAGTACTTTTTAAAA<br>GTAAAAAATCAGATGATTCTTTTTGGGGAGGAGGATGTGCGAGTGGCTAATTTGAAACCAAGAA<br>TCTCCTTTAATTTCTTTGTCACTGCACCTAAAAATGTGTCTGATATCATTCCTAGAACTGAAGT<br>TGAAAAGGCCATCAGGATGTCCCGGAGCCGTATCAATGATGCTTTCCGTCTGAATGACAACAGC<br>CTAGAGTTTCTGGGGATACAGCCAACACTTGGACCTCCTAACCAGCCCCCTGTTTCC |
| 373 | CGATCTACCATCGAAGAGCAGGCCAAAACATTCCTCGACTTCTTTGATAGCCAGGCTGAAGACC<br>TTTTCTACCAATCAAGTCTGGCAAGCTGGAATTACAACACAAACATTACAGAGGAGAACGTACA<br>AAACATGAATAACGCAGGGGACAAGCGGAGCGCATTCCTTAAGGAACGAAGTACCCTTGCGCAG<br>ATGTATCCGCTGCAAGAGATTCAAAACCTGACGGTTAAGCTGCAACTTCAGGCCCTCCAACAAA<br>ATGGATCCTCAGTCTTGTCAGAAGACAAAAGCAAGCGACTGAACACCATCCTTAACACCATGTC<br>AACCATATATTCAACAGGTAAAGTTTGCAATCCGGATAACCCCCAAGAATGTTTGCTTCTTGAA<br>CCCGGTCTCAACGAAATTATGGCCAACAGTCTTGATTACAACGAGCGATTGTGGGCATGGGAAA<br>GTTGGAGGAGTGAGGTAGGCAAACAGTTGAGACCTCTTTATGAAGAGTACGTTGTCCTTAAAAA<br>TGAAATGGCTCGCGCGAATCATTATGAAGACTATGGTGACTACTGGAGGGGGGATTATGAGGTG<br>AACGGGGTGGACGGATACGATTACTCTAGGGGCCAGCTGATAGAGGATGTCGAGCACACCTTTG<br>AGGAGATTAAGCCGTTGTACGAACATTTGCACGCCTATGTCAGGGCTAAGCTCATGAACGCTTA |

| | |
|---|---|
| | TCCGAGTTATATCTCCCCGATAGGATGCTTGCCTGCTCACTTGTTGGGCGATATGTGGGGACGC<br>TTTTGGACCAACTTGTATTCCCTTACGGTACCGTTCGGCCAGAAACCAAATATCGACGTGACAG<br>ACGCAATGGTGGATCAAGCATGGGATGCGCAACGAATCTTCAAGGAGGCAGAAAAATTTTTCGT<br>TTCAGTTGGACTCCCAAACATGACGCAGGGTTTCTGGGAGAACTCAATGTTGACAGATCCAGGT<br>AATGTGCAGAAAGCGGTTTGCCTCCCTACTGCATGGGATCTTGGTAAAGGGGACTTCCGCATAC<br>TCATGTGTACGAAAGTAACTATGGACGACTTTCTTACTGCGCACCACGAGATGGGGCACATACA<br>ATACGATATGGCGTACGCAGCTCAACCTTTCCTTCTGCGGAACGGGGCGAATGAAGGATTTCAC<br>GAGGCAGTGGGTGAGATTATGTCCCTGTCAGCTGCCACTCCGAAACATCTGAAAAGCATCGGCC<br>TGTTGAGCCCAGACTTCCAAGAAGATAATGAGACCGAAATAAACTTCCTTCTGAAGCAAGCACT<br>GACTATTGTAGGTACCTTGCCCTTTACATACATGCTGGAGAAGTGGAGGTGGATGGTATTTAAG<br>GGGGAGATACCGAAAGATCAATGGATGAAAAAGTGGTGGGAAATGAAAAGGGAGATCGTTGGCG<br>TAGTTGAACCAGTACCGCATGATGAGACGTACTGCGATCCGGCTAGTCTGTTCCATGTCTCTAA<br>TGATTACTCTTTCATCCGCTACTACACCCGCACGCTGTATCAATTCCAGTTCCAAGAAGCTCTC<br>TGTCAGGCTGCCAAGCACGAAGGACCGCTGCACAAATGCGACATTAGCAATTCTACAGAGGCGG<br>GTCAGAAGTTGTTCAATATGCTTAGACTGGGGAAGAGCGAACCGTGGACGCTCGCTTTGGAGAA<br>CGTTGTTGGAGCTAAGAATATGAACGTCAGGCCCTTGCTGAATTACTTTGAACCTCTGTTTACG<br>TGGTTGAAAGACCAAAATAAAAACTCCTTTGTTGGGTGGAGTACTGACTGGTCCCCCTATGCGG<br>ACCAAAGCATCAAAGTGAGGATAAGCCTAAAATCAGCTCTTGGAGATAAAGCATATGAATGGAA<br>CGACAATGAAATGTACCTGTTCCGATCATCTGTTGCATATGCTATGAGGCAGTACTTTTTAAAA<br>GTAAAAAATCAGATGATTCTTTTTGGGGAGGAGGATGTGCGAGTGGCTAATTTGAAACCAAGAA<br>TCTCCTTTAATTTCTTTGTCACTGCACCTAAAAATGTGTCTGATATCATTCCTAGAACTGAAGT<br>TGAAAAGGCCATCAGGATGTCCCGGAGCCGTATCAATGATGCTTTCCGTCTGAATGACAACAGC<br>CTAGAGTTTCTGGGGATACAGCCAACACTTGGACCTCCTAACCAGCCCCCTGTTTCC |
| 375 | CAACCTACCATCGAAGAGCAGGCCAAAACATTCCTCGACAAGTTTAGTGTCGAGGCTGAAGACC<br>TTCTCTACCAATCAAGTCTGGCTAGCTGGGATTACAACACAAACATTACAGAGGAGAACGTACA<br>AAACATGAATAACGCAGGGGACAAATGGAGCGCATTCCTCAAGGAACAAAGTACCCTTGCGCAA<br>ATGTATCCGCTGCAAGAGATTCAAAACCTGACGGTTAAGCTGCAACTTCAGGCCCCCCAACAAA<br>ATGGATCCTCAGTCTTGTCAGAAGACAAAAGCAAGCGACTGAACACCATCCTTAACACCATGTC<br>AACCATATATTCAACAGGTAAAGTTTGCAATCCGGATAACCCCCAAGAATGTTTGCTTCTTGAA<br>CCCGGTCTCAACGAAATTATGGCCAACAGTCTTGATTACAACGAGCGATTGTGGGCATGGGAAA<br>GTTGGAGGAGTGAGGTAGGCAAACAGTTGAGACCTCTTTATGAAGAGTACGTTGTCCTTAAAAA<br>TGAAATGGCTCGCGCGAATCATTATGAAGACTATGGTGACTACTGGAGGGGGGATTATGAGGTG<br>AACGGGGTGGACGGATACGATTACTCTAGGGGCCAGCTGATAGAGGATGTCGAGCACACCTTTG<br>AGGAGATTAAGCCGTTGTACGAACATTTGCACGCCTATGTCAGGGCTAAGCTCATGAACGCTTA<br>TCCGAGTTATATCTCCCCGATAGGATGCTTGCCTGCTCACTTGTTGGGCGATATGTGGGGACGC<br>TTTTGGACCAACTTGTATTCCCTTACGGTACCGTTCGGCCAGAAACCAAATATCGACGTGACAG<br>ACGCAATGGTGGATCAAGCATGGGATGCGCAACGAATCTTCAAGGAGGCAGAAAAATTTTTCGT<br>TTCAGTTGGACTCCCAAACATGACGCAGGGTTTCTGGGAGAACTCAATGTTGACAGATCCAGGT<br>AATGTGCAGAAAGCGGTTTGCCTCCCTACTGCATGGGATCTTGGTAAAGGGGACTTCCGCATAC<br>TCATGTGTACGAAAGTAACTATGGACGACTTTCTTACTGCGCACCACGAGATGGGGCACATACA<br>ATACGATATGGCGTACGCAGCTCAACCTTTCCTTCTGCGGAACGGGGCGAATGAAGGATTTCAC<br>GAGGCAGTGGGTGAGATTATGTCCCTGTCAGCTGCCACTCCGAAACATCTGAAAAGCATCGGCC<br>TGTTGAGCCCAGACTTCCAAGAAGATAATGAGACCGAAATAAACTTCCTTCTGAAGCAAGCACT<br>GACTATTGTAGGTACCTTGCCCTTTACATACATGCTGGAGAAGTGGAGGTGGATGGTATTTAAG<br>GGGGAGATACCGAAAGATCAATGGATGAAAAAGTGGTGGGAAATGAAAAGGGAGATCGTTGGCG<br>TAGTTGAACCAGTACCGCATGATGAGACGTACTGCGATCCGGCTAGTCTGTTCCATGTCTCTAA<br>TGATTACTCTTTCATCCGCTACTACACCCGCACGCTGTATCAATTCCAGTTCCAAGAAGCTCTC |

| | |
|---|---|
| | TGTCAGGCTGCCAAGCACGAAGGACCGCTGCACAAATGCGACATTAGCAATTCTACAGAGGCGG<br>GTCAGAAGTTGTTCAATATGCTTAGACTGGGGAAGAGCGAACCGTGGACGCTCGCTTTGGAGAA<br>CGTTGTTGGAGCTAAGAATATGAACGTCAGGCCCTTGCTGAATTACTTTGAACCTCTGTTTACG<br>TGGTTGAAAGACCAAAATAAAAACTCCTTTGTTGGGTGGAGTACTGACTGGTCCCCCTATGCGG<br>ACCAAAGCATCAAAGTGAGGATAAGCCTAAAATCAGCTCTTGGAGATAAAGCATATGAATGGAA<br>CGACAATGAAATGTACCTGTTCCGATCATCTGTTGCATATGCTATGAGGCAGTACTTTTTAAAA<br>GTAAAAAATCAGATGATTCTTTTTGGGGAGGAGGATGTGCGAGTGGCTAATTTGAAACCAAGAA<br>TCTCCTTTAATTTCTTTGTCACTGCACCTAAAAATGTGTCTGATATCATTCCTAGAACTGAAGT<br>TGAAAAGGCCATCAGGATGTCCCGGAGCCGTATCAATGATGCTTTCCGTCTGAATGACAACAGC<br>CTAGAGTTTCTGGGGATACAGCCAACACTTGGACCTCCTAACCAGCCCCCTGTTTCC |
